# Comprehensive Biosynthetic Analysis of Human Microbiome Reveals Diverse Protective Ribosomal Peptides

**DOI:** 10.1101/2024.07.16.603675

**Authors:** Jian Zhang, Dengwei Zhang, Yi Xu, Junliang Zhang, Runze Liu, Ying Gao, Yuqi Shi, Peiyan Cai, Zheng Zhong, Beibei He, Xuechen Li, Hongwei Zhou, Muxuan Chen, Yong-Xin Li

## Abstract

The human microbiome holds tremendous potential for generating specialized peptides, specifically ribosomally synthesized and post-translationally modified peptides (RiPPs), significantly affecting human health by mediating interactions with other microbes and the human host. However, the capacity of our microbiome to produce these peptides and their links with human health are poorly understood. This study systematically analyzes 306,481 human microbiota-associated genomes, uncovering a broad array of yet-to-be-discovered RiPPs. These RiPPs are distributed across various body sites but show a specific enrichment in the gut and oral microbiome. Big data omics analysis reveals that numerous RiPP families are inversely related to various diseases, suggesting their potential protective effects on health. For a proof of principle study, guided by biosynthetic prediction, nine autoinducing peptides (AIPs) were chemically synthesized for *in vitro* and *ex vivo* assay. Our findings revealed that five AIPs effectively inhibited the biofilm formation of disease-associated pathogens, with one demonstrating significant anti-inflammatory capabilities. Furthermore, when *ex vivo* testing gut bacteria from mice with inflammatory bowel disease, we observed that two AIPs could regulate the microbial community and reduce harmful species. These findings highlight the vast potential of human microbial RiPPs in regulating microbial communities and maintaining human health, emphasizing their potential for therapeutic development.

## Introduction

Humans and their symbiotic microbes form a complex and diversified holobiont. Perturbation of the human microbiota, known as dysbiosis, has been implicated in the pathogenesis of various diseases^1,2^, which is largely attributed to the chemical interactions mediated by metabolites *de novo* biosynthesized or transformed by the colonized microbiota^3-6^. As a result, extensive efforts have been made to decipher bioactive molecules secreted by the human microbiota^7,8^, revealing a rather diverse array of metabolites such as polyketides (PKs), non-ribosomal peptides (NRPs), and ribosomally synthesized and post-translationally modified peptides (RiPPs). RiPPs have particularly attracted increasing attention because of their diverse structures^9^, various biological functions, and manifold roles in physiology and ecology^10,11^.

RiPPs are a class of diverse and widespread natural products produced by a multitude of producers, including bacteria, fungi, and plants^10^. They often exhibit diverse activities^12^, allowing them to mediate the communications between microbes or between microbes and human hosts. Antibacterial RiPPs can enable commensal producers to combat pre-existing or foreign pathogens^13^. For example, lactocilin, a thiopeptide from vaginal *Lactobacillus gasseri*, demonstrates potent antibacterial effects against vaginal pathogens such as *Corynebacterium aurimucosum*^14^. Nisin-like lantibiotics identified in gut microbiome against both pathogens and human gut commensals could shed light on the future development of therapeutics^15^. Additionally, immunomodulatory RiPPs^16-19^ can interact with the human immune system, exemplified by tonsillar microbiome-derived Salivaricins, which can block the binding of IL-6 and IL-21 to their receptors to disrupt the immune response^16^. Beyond impacting the microbial community or host directly, RiPPs can also act as signaling molecules to regulate human microbiome responses, as evidenced by autoinducing peptides (AIPs) from the skin microbiome, which maintain skin barrier homeostasis by disrupting the *agr* system of the *Staphylococcus aureus* pathogen^20^. These observations underscore the diverse chemical nature of RiPPs and their varied protective roles to human health. Nonetheless, our current understanding in this area is primarily limited to sporadic reports, highlighting the urgent imperative for a more comprehensive investigation.

While over 40 subclasses of RiPPs are differentiated by their unique structures, their biosynthesis follows a standard three-step process controlled by biosynthetic gene clusters (BGCs)^9,21^. The biosynthesis process includes expressing a precursor peptide, installing various structural features to the core peptide through post-translational modifications (PTM), and finalizing with leader cleavage and export to produce active RiPP products. The conserved biosynthetic logic allows the development of computational approaches for RiPP discovery, rendering diverse tools poised for RiPP identification from the human microbiome^22,23^. These tools typically are established on precursor-centric genome mining strategies (e.g., DeepRiPP^24^ and TrRiPP^25^) or primarily targeting the PTM enzymes (e.g., antiSMASH^26^ and DeepBGC^27^). Leveraging various bioinformatics tools, researchers are now focusing on understanding the biosynthetic potential of RiPPs within the complex microbiota. Some studies have concentrated on the biosynthetic potential of specific strains, as illustrated by the discovery of bioactive RiPPs like Ruminococcin C^28^ and Streptosactin^29^ from human commensal bacteria *Ruminococcus gnavus* and *Streptococcus* spp., respectively. Others have centered on in silico investigation of RiPP from a limited number of reference genomes^14,30-32^. However, our understanding of the biosynthetic landscape of RiPPs from human microbiomes is still limited, hindered by a lack of explored reference genomes and imperfect prediction tools. As such, a more thorough examination of RiPPs is needed to improve our understanding of their biosynthesis, laying the groundwork for understanding the crucial biological role of RiPPs within the complex microbiota.

The exponential increase in human microbial genome data provides an opportunity for systematically exploring RiPPs biosynthesis in the human microbiome and their potential role in human health. In this study, using precursor-centric and PTM-centric approaches, we systematically examined the RiPP biosynthetic potential of 306,481 genomes of the human microbiota to identify 12,076 yet-to-be-discovered RiPP families, largely expanding the chemical landscape of RiPPs. For a proof of principle study, we have found 30 RiPP families, through comparative meta-omic analyses, that are negatively associated with multiple diseases such as inflammatory bowel disease (IBD) and colorectal cancer (CRC). Experimental validation of biosynthesis-guided chemically synthesized AIPs demonstrated their potential to counter biofilm formation of disease-related pathogens and exhibit anti-inflammatory activity. Using an *ex vivo* culture to simulate the gut microbiota’s response to AIPs, we identified two AIPs capable of modifying the gut microbial community and reducing pathogenic species from IBD-affected mice. These findings offer valuable insights into the chemical diversity of RiPPs and their protective roles in human health, sparking growing interest in the therapeutic potential of RiPPs released by our microbiota.

## Results

### Mining human microbiomes reveals the untapped biosynthetic potential of RiPPs

Current genome mining for RiPPs can be approached through two main methods: tailoring enzymes-oriented approach (e.g., antiSMASH 6.0^26^) and precursors-centric approach (e.g., DeepRiPP^24^ and TrRiPP^25^). The enzymes-oriented approach is reliable in identifying known RiPPs but may miss out on discovering novel ones. On the other hand, the precursors-centric approach is better suited for identifying novel RiPPs, especially from highly fragmented metagenome-assembled genomes but may have a slightly lower accuracy rate. To comprehensively explore the biosynthetic potential of RiPPs in the human microbiomes, we combined both types of tools to analyze a dataset consisting of 306,481 reference microbial genomes from diverse human body sites^33-35^, including the gut, oral cavity, skin, airways, vagina, and nasal cavity (**Fig. 1, Supplementary Fig. 1**). Results showed that most genomes (78.3% by antiSMASH and 95.3% by DeepRiPP and TrRiPP) could encode RiPPs.

**Fig 1.**
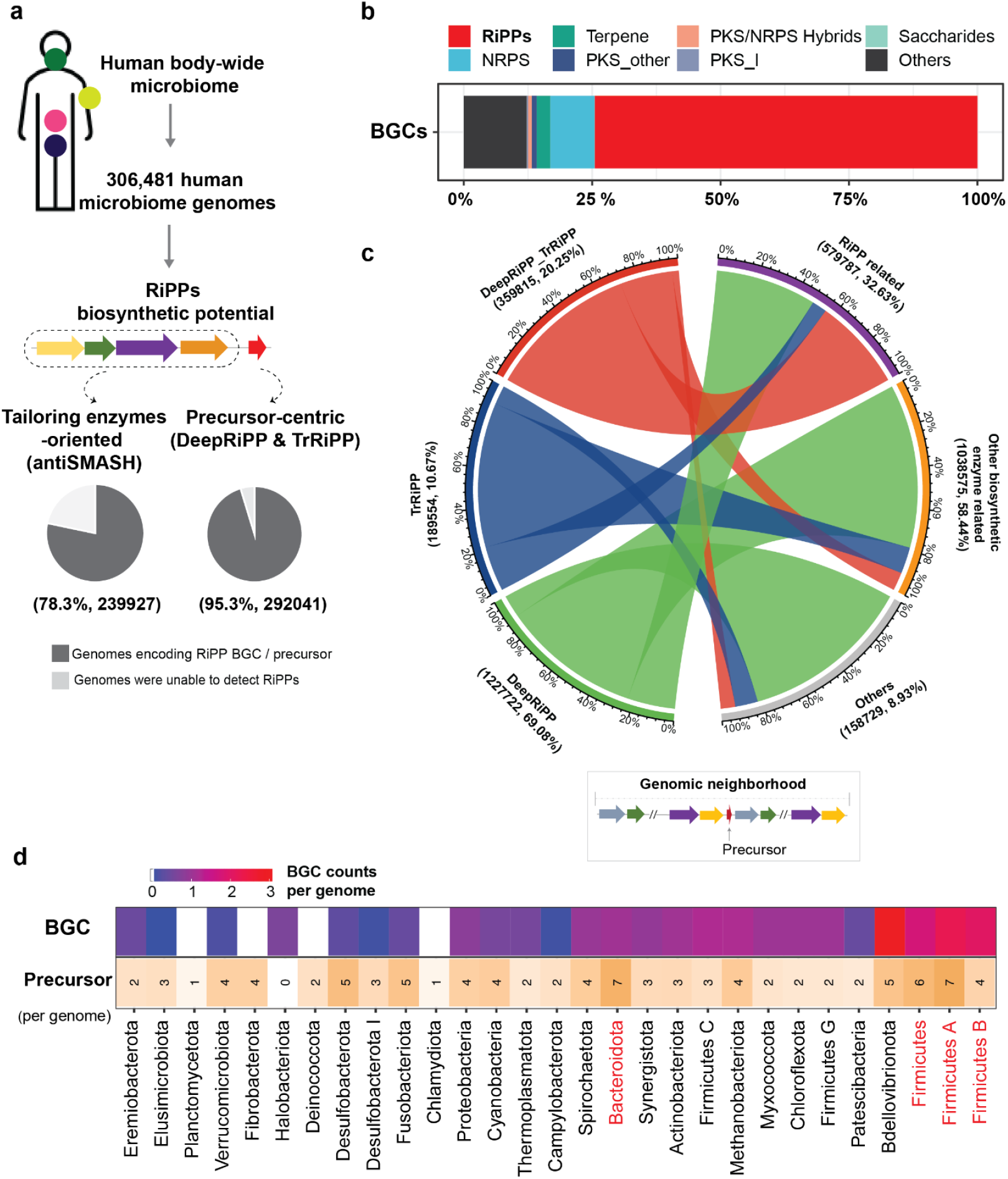
Comprehensive survey of RiPPs in the human microbiome. **a** The workflow of identifying RiPP BGCs and RiPP precursors from 30,6481 human-associated microbial genomes collected from wide body sites. Tailoring enzymes-oriented tool (e.g., antiSMASH) and precursors-centric tools (e.g., DeepRiPP and TrRiPP) are adopted for predicting RiPP BGCs and RiPP precursors, respectively. The pie charts show the percentage of genomes that encode RiPPs. **b** BGCs identified by antiSMASH are grouped into eight BGC classes, including RiPPs (410487, 74.5%), Terpene (14753, 2.7%), PKS/NRPS Hybrids (4067, 0.7%), Saccharides (17, 0.003%), NRPS (47281, 8.6%), PKS_others (5210,0.9%), PKS_I (1506,0.3%), and Others (67328, 12.2%). **c** Genomic context of identified RiPP precursors by DeepRiPP and TrRiPP. The left panel of the Fig. displays the scales representing the proportion of precursors identified by either DeepRiPP or TrRiPP, or both approaches (DeepRiPP_TrRiPP). The right panel of the Fig. illustrates the genomic context associated with the identified precursors. This includes:(1) RiPP related: Genes in the genomic context that are associated with known RiPP biosynthesis. (2) Other biosynthetic enzyme related: Precursors that co-occur with potentially novel RiPP biosynthetic enzyme(s) from a broader PTM enzyme dataset collected by decRiPPter^36^. (3) Others: Precursors located in the biosynthetic gene cluster (BGC) region of other secondary metabolite BGC classes or under other conditions. **d** The top section of the bar chart represents the counts of BGCs per genome in each phylum, while the bottom section represents the counts of RiPP precursors per genome in each phylum. Taxonomic classification was determined based on annotations from the Genome Taxonomy Database (GTDB). Phyla with higher RiPP biosynthetic potential are highlighted in red.

The analysis of antiSMASH detected 410,487 RiPP BGCs in 239,927 genomes, accounting for 74.5% of the predicted secondary metabolite BGCs (**Fig. 1b, Supplementary Fig. 1, Supplementary Data 1**). The other two precursor-centric tools identified 1,777,091 RiPP precursors in 292,041 genomes, with 359,815 precursors overlapping between the two tools (**Fig. 1c, Supplementary Fig. 2, Supplementary Data 2**). Through examining the adjacent genes of RiPP precursors identified by precursor-centric approaches, we found that the genomic contexts of 32.6% of precursors contained genes associated with known RiPP biosynthesis (**Fig. 1c, Supplementary Fig. 3, Supplementary Tables 1-2**), while the genomic contexts of 58.4% of precursors contained potentially novel RiPP biosynthetic enzymes collected in decRiPPter^36^ (**Fig. 1c**). These findings indicated that the precursor-centric approach offered a higher level of biosynthetic novelty. The analysis also showed that antiSMASH-defined RiPP BGCs were more abundant in 25 phyla, expanding to 29 phyla with the precursor-centric approaches (**Fig. 1d, Supplementary Fig. 1**). Notably, the most abundant phyla in the human gut^37^, Bacteroidota and Firmicutes, harbored the most abundant RiPPs. This widely distributed RiPPs across diverse human microbiomes underscores human microbes’ robust RiPP biosynthetic potential.

**Fig 2.**
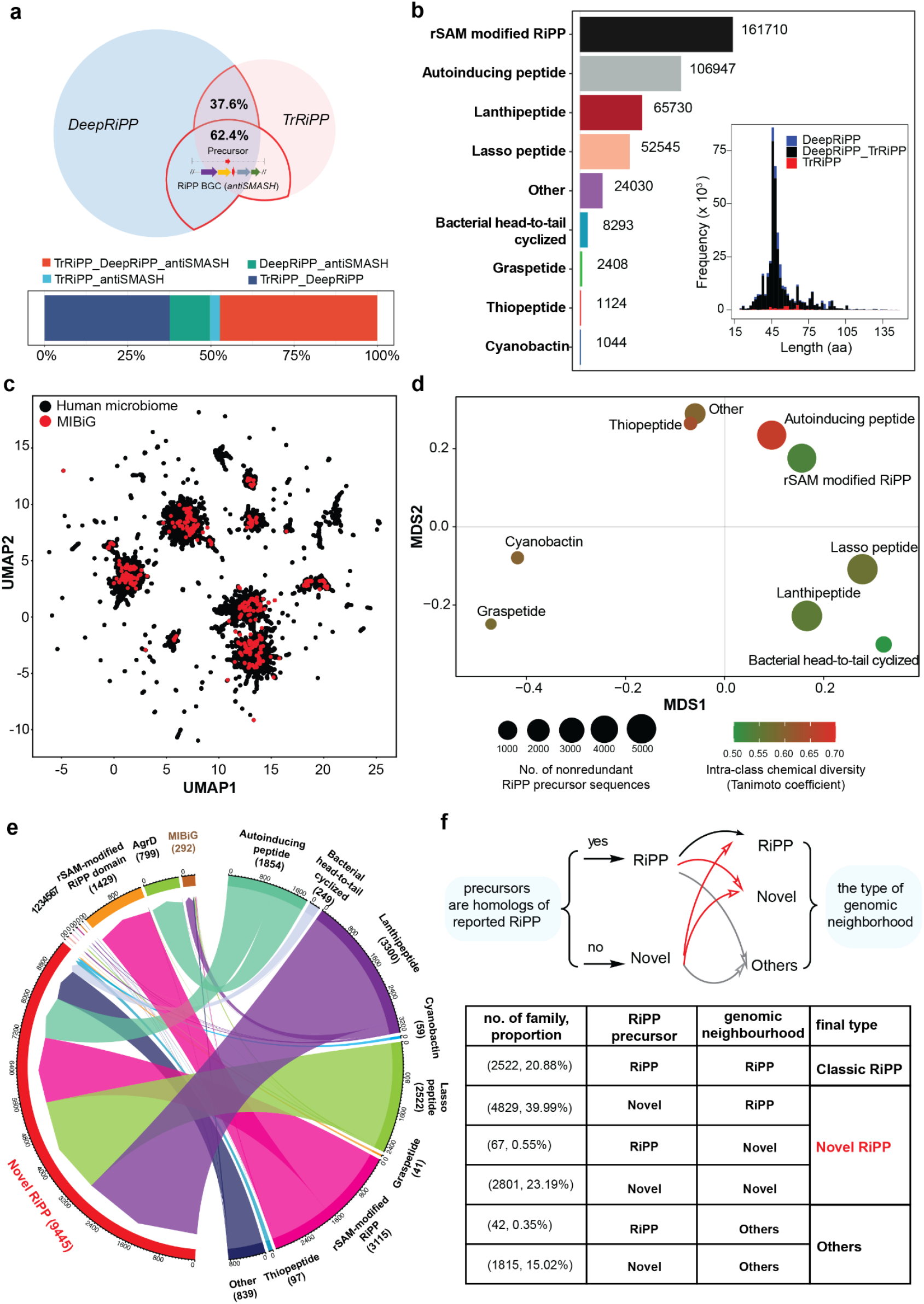
RiPP precursor peptides present a hypervariable chemical diversity and novelty. **a** RiPP precursors retained for downstream analysis. Upper: Two circles in different colors represent the RiPP precursor peptides identified by DeepRiPP and TrRiPP. The red-circled area highlights the precursors that were retained for downstream analysis. Bottom: The stacked barplot illustrates the proportion of RiPP precursors retained for analysis. **b** The outer barplot displays the count of precursor sequences for nine RiPP classes, while the inner stacked barplot represents the distribution of precursor length. **c** Uniform Manifold Approximation and Projection (UMAP) plot showing the chemical space of RiPP precursors obtained from the human microbiome (black dots) and experimentally validated RiPP precursors deposited in the MIBiG 3.0 database (red dots). **d** Multi-Dimensional Scaling (MDS) plot displays the chemical diversity of predicted mature precursors within and between RiPP classes. Dot size signifies the count of unique precursor sequences per class, color indicates median Tanimoto coefficient reflecting class similarity, and distance between dots represents similarity among different RiPP classes. (**e** & **f**) Classify the novelty of RiPP families based on their precursor and genomic neighborhood **(Supplementary Information). e** The chord diagram illustrates the novelty of identified precursor families (left panel) for nine different RiPP classes (right panel). The novelty of RiPP precursor families into “MIBiG” (homologous to characterized precursors in MIBiG), families with RiPP-associated domains (1, Graspetide (3 families), 2, Lanthipeptide (31 families), 3, Lassopeptide (26 families), 4, Thiopeptide (5 families), 5, RiPP-like (25 families), 6, LAP (19 families), and 7, other known RiPPs (2 families)), and “Novel RiPP” families. Numbers in brackets indicate the count per category. **f** RiPP families are classified as classic RiPP (known precursor homology and defined genomic neighborhood), Novel RiPP (no known precursor homology or novel genomic neighborhood), and Others (remaining families).

**Fig 3.**
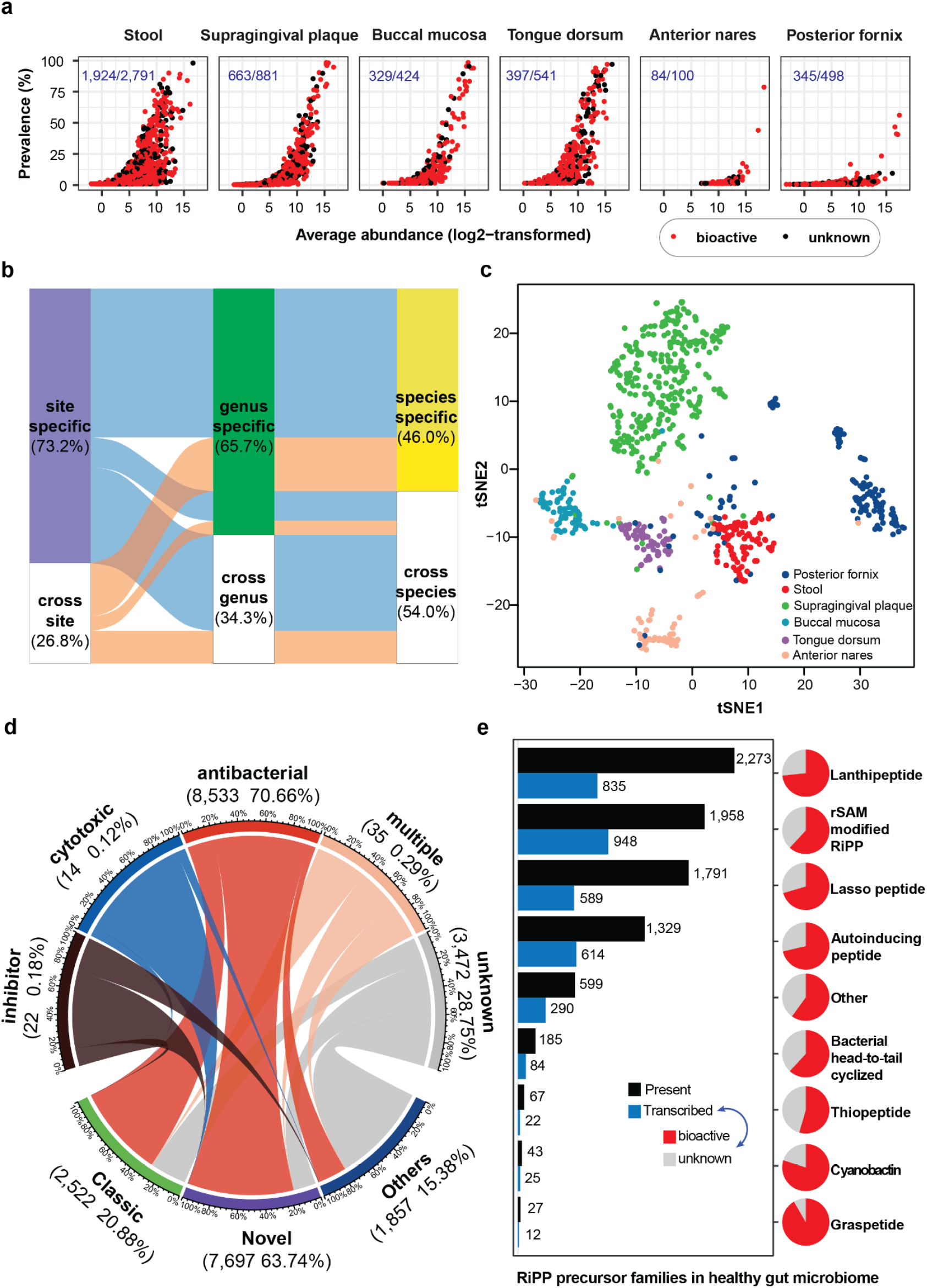
Identified RiPP families are present and transcribed in the healthy human microbiome. **a** The data represents RiPP precursor families’ prevalence and average abundance across six human body sites. Each dot signifies a family, with bioactive ones in red and unknown ones in black. Red numbers denote the count of bioactive and total families per site. **b** The Sankey diagram illustrates the distribution of RiPP families in the healthy human microbiome, categorizing them as either niche-specific (present in one site) or cross-niche (present in more than one site), and further differentiating them as genus/species-specific or genus/species-cross. **c** The t-SNE plot showcases the distinct profiles of RiPP precursor families in different body sites. The clustering of RiPP families within each body site indicates conservation. Each dot on the plot represents one metagenome sample. **d** Predicted activity of RiPP families. The scale represents the proportion of each predicted activity (upper) or RiPP novelty (bottom). The number in brackets indicates the count and percentage of the RiPP family relative to all families. The term “multiple” indicates multi-functional RiPPs. Different colors are used to highlight the connections between RiPP types and their predicted activities. **e** Presence and transcription of all RiPP classes in 281 paired metagenomic and metatranscriptomic data from fecal samples of healthy individuals. The pie charts present the percentage of transcribed RiPP families in each class that predicted bioactivity (red) or unknown (grey).

### Broad diversity and novelty of RiPP precursor peptides in the human microbiome

Given the tremendous potential of RiPP biosynthesis in the human microbiome, we next sought to examine their chemical diversity and novelty. To balance the accuracy and novelty of genome mining in our analysis, we only retained 423,831 RiPP precursors confirmed by at least two mining strategies for downstream analysis, which could effectively reduce the number of false positives while preserving new and novel RiPPs. The 423,831 RiPP precursors were found either located within RiPP BGCs region^38^ identified by an antiSMASH (62.4%) or by both precursors-centric approaches (37.6%) (**Fig. 2a, Supplementary Data 3**), with rSAM-modified RiPPs being the most abundant (**Fig. 2b**). These RiPPs were strongly enriched in 21 phyla, 1,014 genera, and 3,369 species. At the genus level, an appropriate taxonomic rank for assessing secondary metabolite biosynthetic diversity^39^, we found that RiPP biosynthetic potential considerably varied from 1 to 33 RiPP precursors per genome, with *Elizabethkingia* (n = 33), *Chryseobacterium* (n = 18) and *Tissierella* (n = 15) having the highest number of RiPP precursors **(Supplementary Fig. 4, Supplementary Information)**.

**Fig 4.**
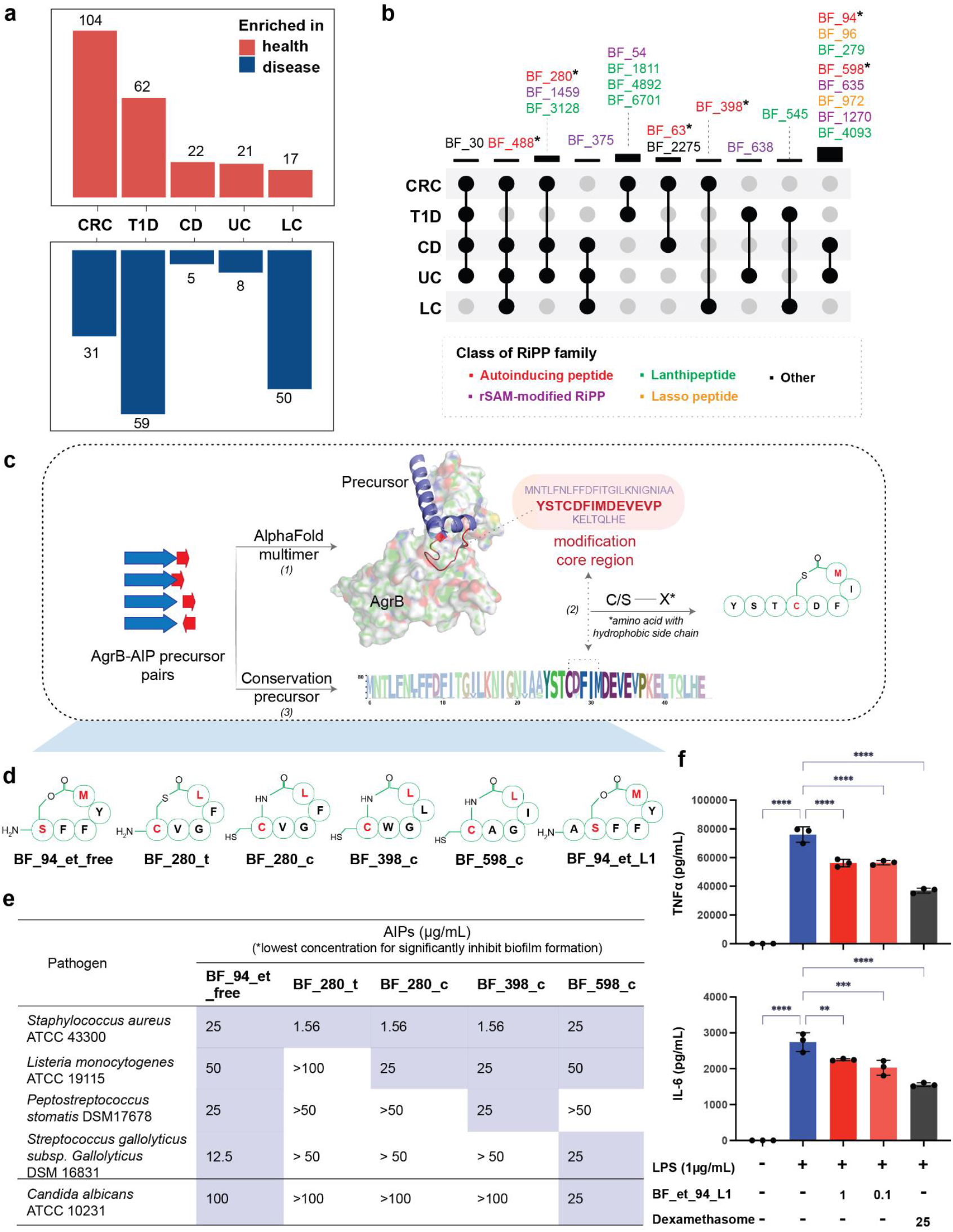
Differentially transcribed RiPP families in multi-disease case-control cohorts. **a** The bar plot displays the number of RiPP precursor families that are significantly enriched in the healthy group (top) or the disease group (bottom). Statistical significance was determined using criteria of |log2 fold change| ≥1 and adjusted *p* values ≤ 0.05. **b** The intersection of differentially transcribed RiPP precursor families that are enriched in healthy groups compared to multiple disease groups. The top bar plot and connecting lines represent the differential RiPP precursor families identified in the corresponding disease case-control cohorts. Precursor families are highlighted in different colors based on their RiPP classes.*Families chosen to tested bioactivity in this study. **c** The workflow for predicting the mature AIP uses the reported AIP-I from *Staphylococcus aureus* as an illustrative example **(Method). d** Chemically synthesized AIPs with antibiofilm or anti-inflammatory activity in this study. **e** Anti-biofilm activity of AIPs against pathogenic biofilm by pathogens. *n = 3 biologically independent samples. **f** Anti-inflammatory activity of BF_94_et_L1 in RAW264.7 cell lines. (E & F)n = 3 biological samples for each experiment, 2 independent experiments. Significance was determined using one-way ANOVA test. Bars represent mean ± standard error. For all *p* values, *p* < 0.05 mean significant difference compared with the control group.

To evaluate RiPP sequence diversity, we calculated the Extended-connectivity fingerprints (ECFPs)^40^ of each RiPP precursor, a topological fingerprint for molecular characterization utilized in two-dimensional chemical space. Comparing them with experimentally validated RiPP precursors in the MIBiG database (3.0)^6^, we observed a notably expanded chemical space (**Fig. 2c, Supplementary Data 4**), suggesting that human microbiome-derived RiPPs could significantly enlarge the chemical diversity within the RiPP superfamily. Next, we explored the chemical diversity of nine RiPP classes based on the core precursor sequences predicted by the cleavage prediction module of DeepRiPP^24^ **(Fig. 2d, Supplementary Data 4)**. Pairwise Tanimoto coefficient was used to measure the similarity between precursors. Notably, autoinducing peptides exhibited the highest median Tanimoto coefficient, suggesting that their sequences were comparatively conserved within the class. In contrast, bacterial head-to-tail cyclized precursors had the lowest Tanimoto coefficient, indicating high intra-class sequence diversity. Moreover, we found that different RiPP classes were far apart, indicating their sequence uniqueness. Overall, RiPP precursor peptides in the human microbiome exhibit a remarkably diverse chemical landscape characterized by both intra-class and inter-class variations.

To further assess the novelty of RiPPs in the human microbiome, we examined their precursor sequences and tailoring enzymes, which can dictate the chemical structure of mature RiPP products (**Fig. 2e and 2f, Supplementary Figs. 5-6, Supplementary Information**). When evaluating the chemical novelty of RiPPs based on their precursors, we first clustered 423,831 RiPP precursors into 12,076 families with a threshold of 50% identity (**Fig. 2e, Supplementary Data 5)**. Through querying these families against known RiPP precursors deposited in the MIBiG database, we surprisingly found that only 292 (2.41%) families were homologous. For the remaining families, we used RPS-BLAST to search protein domains against the Conserved Domains Database (CDD), detecting 2,339 families (19.37%) containing well-defined RiPP precursor domains, such as AgrD domain and rSAM-modified RiPP domain. Notably, most families (9,445, 78.21%) remained uncharacterized or exhibited significant distinctions from known RiPPs **(Fig. 2e)**. We examined the adjacent genomic context of RiPP precursor families when evaluating the novelty based on tailoring enzymes. We found that the genomic contexts of 60.9% RiPP precursor families harbored classical RiPP biosynthetic-related genes, while the genomic contexts of 23.8% RiPP precursor families contained novel RiPP biosynthetic-related enzymes defined by DeepBGC^27^ (**Fig. 2f**). Considering both aspects, 63.7% of RiPP precursor families have the potential to produce novel RiPP products (**Fig. 2f**). This finding suggests the possibility of novel enzymology and chemistry in these RiPPs families.

**Fig 5.**
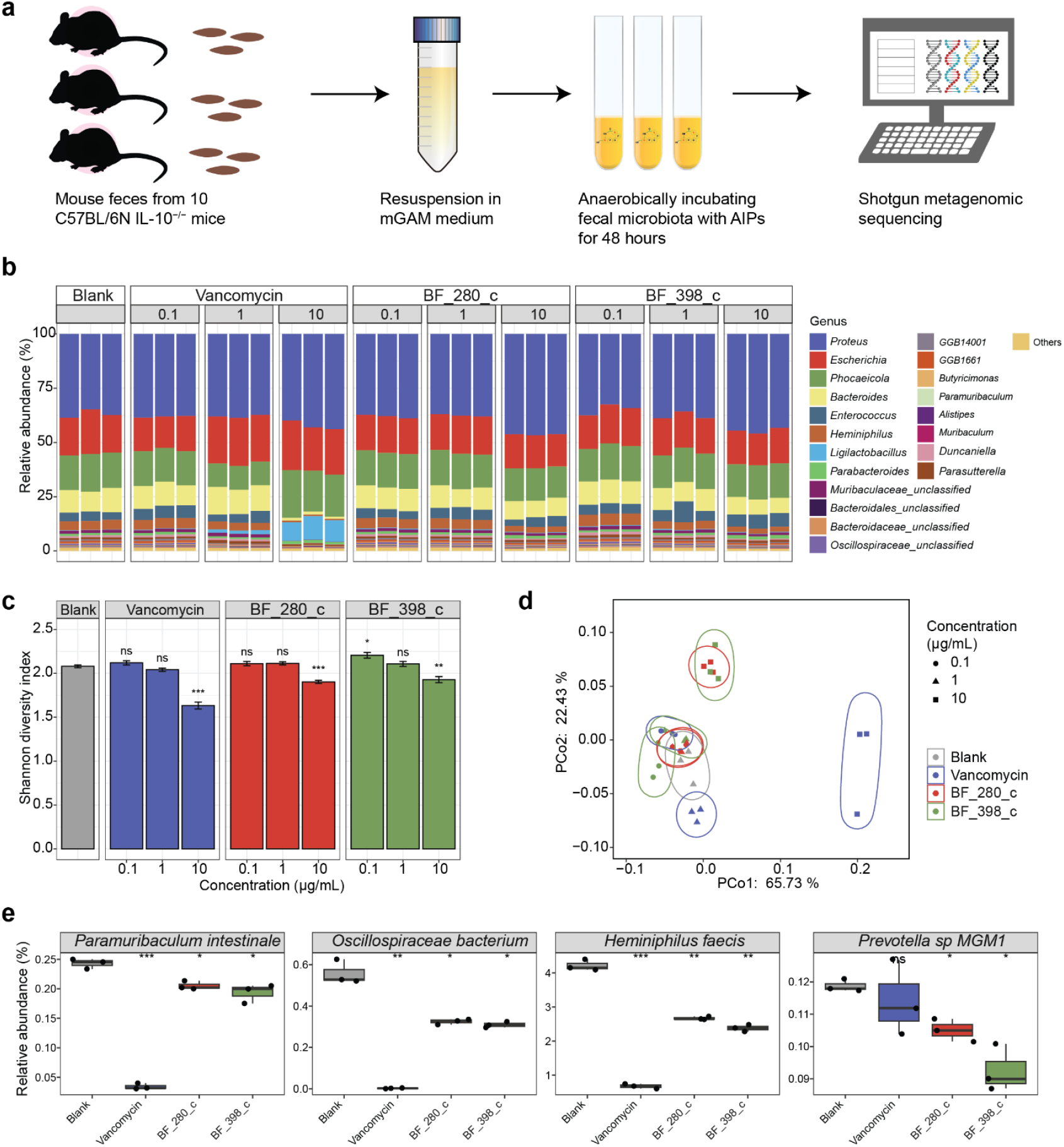
AIPs regulate IBD mouse fecal-derived ex vitro microbial community. **a** Graphical depiction of the *ex vivo* experimental setup. **b** Relative abundance of microbial genera after 48 h of treatment with vancomycin, BF_280_c, and BF_398_c at concentrations of 0.1, 1, and 10 μg/mL. DMSO was used as a blank control. Only 20 abundant species are displayed. **c** Bar plots show the alpha diversity of the microbial community at the species level, as represented by the Shannon diversity index. Data are mean ± standard deviation. Significances between treatment groups and blank groups were indicated by using two-sided Welch’s *t*-test. **d** Principal coordinate analysis (PCoA) of microbial community based on the Bray-Curtis dissimilarity at the species level. The ellipses represent the distinct clustering of groups. **e** Relative abundance of four representative taxa. All box plots include center lines representing the median, box limits representing upper and lower quartiles, whiskers representing the 1.5x interquartile range, and points representing outliers. MaAsLin2 was used to compare the species abundance among groups. \**p* < 0.05; \*\**p* < 0.01; \*\*\**p* < 0.001; ns, not significant.

### RiPPs are niche-specific and actively transcribed in the healthy human microbiome

We then attempted to assess RiPP distribution in the human microbiome by examining their profile in healthy human microbiomes. We analyzed 748 metagenomic samples from the Human Microbiome Project (HMP)^41^, covering six human body sites: gut, buccal mucosa (oral), supragingival plaque (oral), tongue dorsum (oral), anterior nares (skin), and posterior fornix (vaginal) **(Supplementary Table 3)**. Metagenomic analysis revealed 28.63% (3,457/12,076) of RiPP precursor families were detected in healthy individuals, varying considerably from 100 families in anterior nares to 2,791 families in the stool **(Fig. 3a)**. Accumulation curves showed a steep increase, indicating that more RiPP precursor families could be identified in each body site with larger sample sizes **(Supplementary Fig. 7a)**. Compared to the sporadic presence in the skin and vagina, RiPP precursor families exhibited higher diversity, prevalence, and abundance in the oral and gut **(Fig. 3a** and **Supplementary Fig. 7b)**. Among these detected families, 73.2% (2,530/3,457) were niche-specific, while 0.9% (31/3,457) were present across six body sites in at least one sample **(Supplementary Fig. 7c)**. This could be largely attributed to the fact that the producers of niche-specific RiPPs were habitat-specific **(Fig. 3b)**. Furthermore, two-dimensional visualization demonstrated distinct patterns of RiPP precursor families among different body sites, indicating their niche-specificity **(Fig. 3c)**. Activity prediction showed that most RiPPs (70.7%, 8,533/12,076) were antibacterial **(Fig. 3d)**, possibly enabling their producers for niche adaption.

To further confirm whether RiPPs are potentially active in the human microbiome, we looked into their transcription in 281 pairs of metagenomic and metatranscriptomic data^42^ **(Supplementary Table 3)**. Among 8,278 (68.55%, 8,278/12,076) RiPP precursor families detected in metagenome samples, 3,525 (29.19%, 3,525/12,076) were found to be actively transcribed in metatranscriptome data. Furthermore, a significant proportion of them in each class (ranging from 54.5% of thiopeptide to 91.7% of graspetide) were predicted to exhibit bioactivity (**Fig. 3e**). These findings necessitate investigating the potential role of bioactive RiPPs in human health.

### Differentially transcribed RiPP families are associated with multi-diseases

Inspired by numerous functional RiPPs from human microbiota linked to mediate microbe-microbe interaction or microbe-host interaction, we aimed to explore the potential impact of RiPPs on human health. For a proof of principle study, we conducted comparative metatranscriptome analyses to identify underlying bioactive RiPPs associated with multiple diseases, including type 1 diabetes mellitus (T1D), obesity (OB), liver cirrhosis with hepatitis C infection (LC), colorectal cancer (CRC), Parkinson’s disease (PD), and inflammatory bowel disease (IBD, encompassing Crohn’s disease (CD) and ulcerative colitis (UC)) **(Supplementary Table 3)**. The metatranscriptomic analysis showed significant differences in RiPP diversity between the health and disease groups **(Supplementary Fig. 8)**. Specifically, the healthy groups had a lower diversity than patients with OB. Permutational multivariate analysis of variance (PERMANOVA) based on Bray-Curtis dissimilarity showed a significant difference (*p* < 0.05) in the overall composition of transcribed RiPP precursor families between health and disease in CRC and IBD case-control cohorts. In total, we identified 195 precursor families depleted while 146 enriched in disease groups within these disease case-control cohorts **(Fig. 4a, Supplementary Data 6)**. Remarkably, 23 families exhibited depletion across multiple disease groups, while 7 families were enriched in multiple disease groups **(Fig. 4b, Supplementary Fig. 9)**. Some families (e.g., BF_63 and BF_94) significantly differentiated CD patients from healthy individuals in a random forest classification model based on the abundance of transcribed precursor families (**Supplementary Fig. 10)**, suggesting their important roles in maintaining human health. Together, these differentially transcribed RiPPs might contribute to the disease pathogenesis or promote human health^16,43^.

### Autoinducing peptides negatively associated with diseases exhibit anti-pathogenic biofilm and anti-inflammatory activity

Considering the comparative conciseness and predictability of the AIP product, we next sought to experimentally validate the potential functions of six AIP families (BF_63, BF_94, BF_280, BF_398, BF_488, and BF_598), which were enriched in the healthy microbiome **(Fig. 4b, Supplementary Figs. 11-16, Supplementary Table 4)**. The AIP biosynthesis involves the removal of a C-terminal follower and modifications, which are achieved by the AgrB enzyme. The resulting intermediate then undergoes a second cleavage in the N-terminus, ultimately producing mature AIP with variable lengths of the exotail^44^. Although we intended to carry out heterologous expression of six AIPs, this proved unsuccessful. This might be because the protease involved in the second cleavage of AIPs’ biosynthetic pathway is still not fully understood, and the native producers of these six AIPs are unavailable. We, therefore, alternatively attempted to chemically synthesize the mature chemicals of AIPs. Here, we employed a combination of structural prediction and conservation analysis of RiPP precursor to deduce the ultimate structure of AIPs **(Fig. 4c)**. To do so, we initially established this approach on a well-studied AIP-I from *Staphylococcus aureus*. Specifically, we utilized AlphaFold-Multimer to predict the interaction between precursor peptide and AgrB protein and found that the modification core region precisely docks into the catalytic pocket. In the core region, thiolactone or lactone will be installed between Cysteine or Serine and another amino acid containing a hydrophobic side chain by AgrB. Additionally, we can deduce the exocyclic tail region of the final AIPs, as the proteolysis step recognizes and occurs at conserved residues. AIPs containing thiolactone, without the exocyclic region, may rearrange to form homodetic cyclopeptides (cAIPs)^45,46^. Therefore, we could infer the structure of mature AIPs by combining the prediction of core region, modified residues, and exocyclic tail. The accuracy and reliability of this approach were further validated by another two reported AIPs (**Supplementary Fig. 17**).

According to the biosynthetic logics of AIPs, we deduced the potential mature chemicals of six AIP families (**Fig. 4b, Supplementary Fig. 18**) and chemically synthesized 9 AIP chemicals, including 3 variants. These were confirmed through high-resolution MS/MS and NMR **(Fig. 4d, Supplementary Figs. 19-36, Supplementary Information)**. After observing that none of them exhibited significant antibacterial activity against a panel of 17 human pathogens **(Supplementary Table 5)**, even at concentrations of 100 or 50 μg/mL, we proceeded to evaluate their inhibitory activities against the biofilm formation of these pathogens **(Supplementary Figs. 37-38)**. We observed that the 5 AIPs (BF_94_et_free, BF_280_t, BF_280_c, BF_398_c, and BF_598_c) demonstrated significant inhibition against at least one pathogenic biofilm **(Fig. 4e, Supplementary Fig. 38, Supplementary data 7)**, 4 of which showed extended-spectrum activities against pathogenic biofilm across different species. For example, BF_398_c and BF_94_et_free from *Lachnospira* genus, showed relatively potent anti-biofilm activities to *Staphylococcus aureus, Listeria monocytogenes*, and *Peptostreptococcus stomati*. We further explored their immunomodulatory effects *in vitro* in lipopolysaccharide (LPS)-induced mouse macrophage RAW264.7 cells. Using an enzyme-linked immunosorbent assay (ELISA), we observed that BF_94_et_L1 could significantly decrease two pro-inflammatory cytokines, tumor necrosis factor-α (TNF-α) and interleukins 6 (IL-6), at a physiologically relevant low concentration of 0.1 μg/mL (**Fig. 4f, Supplementary data 8**). Collectively, their multiple functions implied that human microbiome-derived AIPs potentially provide a multifaceted protective role in human health.

### AIPs affect the composition of IBD mouse fecal-derived *ex vivo* microbial community

Efforts to investigate the role of AIPs in disease contexts would greatly benefit from studying a system that mimics disease-derived microbial communities. We, thus, adopted an IBD mouse fecal-derived *ex vivo* microbial community to model how the gut microbiota responds to AIPs (**Fig. 5a, Supplementary data 9**). We selected two functional AIPs (i.e., BF_280_c and BF_398_c) for assay, given their potent inhibition against biofilm formation, which is a major virulence factor in IBD^47,48^. Antibiotic vancomycin was used as a positive control due to its clinical use in IBD treatment^49^. After incubating fecal-derived microbiota with AIPs or vancomycin for 48 hours, we found, except for vancomycin (10 μg/mL), no marked difference in major taxa when compared with the blank group (**Fig. 5b, Supplementary Fig. 39, Supplementary data 9**). Nevertheless, despite their lack of antibacterial activity in our tests, both two AIPs (10 μg/mL) significantly reduced the microbial diversity, as reflected by the Shannon diversity index (**Fig. 5c, Supplementary data 9**). Beta diversity analysis utilizing principal coordinate analysis based on Bray-Curtis distances revealed distinct clustering between blank and treatment, with notable differences observed in groups of AIPs (10 μg/mL) and vancomycin (1 μg/mL and 10 μg/mL) (**Fig. 5d, Supplementary Fig. 40, Supplementary data 9**). These results indicated that BF_280_c and BF_398_c could affect the overall microbial community, possibly by (1) manipulating the production of other metabolites such as antibiotics^50^, which could inhibit the growth of pathogens and reconstitution of the protective gut microbiota; (2) regulating (stimulating or inhibiting) the quorum sensing pathway^51,52^, thus affecting the communication between microbes.

Using MaAsLin2^53^ (microbiome multivariable associations with linear models), we found variable number of significantly differentially abundant species (FDR-adjusted *p* value < 0.05) in groups of vancomycin (1 μg/mL and 10 μg/mL), BF_280_c (10 μg/mL), and BF_398_c (0.1 μg/mL and 10 μg/mL) when compared with blank group (**Supplementary Fig. 41, Supplementary data 9**). At the same concentration of 10 μg/mL, vancomycin affected more species (n = 73) than the other two AIPs (n = 16 and 40). Nevertheless, most differentially abundant species were affected in a similar pattern by vancomycin or AIPs at 10 μg/mL concentration (**Supplementary Fig. 41c, Supplementary data 9**), some of which are closely associated with IBD. As depicted in **Fig. 5e**, they inhibited the growth of pathobiont species from genera *Paramuribaculum*^*54*^ (*Paramuribaculum intestinale*), and *Oscillospiraceae*^*55,56*^ (*Oscillospiraceae bacterium*), which are known to be enriched in patients with IBD. Another species, *Heminiphilus faecis*^*57,58*^, which could produce IBD pathogenic factors, was also found to be reduced. Additionally, both AIPs decreased the abundance of pathobiont species from the *Prevotella* genus^59,60^ (*Prevotella sp MGM1*), which are linked to increased susceptibility to mucosal inflammation in IBD. These findings implied that AIPs could modulate the fecal microbial composition and reduce potential pathogens in IBD, justifying their possibly protective effects on human health.

## Discussion

Extensive studies have highlighted the profound functional impact of the host-microbiome interaction on human health, as well as the etiology and progression of various diseases^61^. These research findings underscore the crucial role that human microbiota plays in health maintenance by producing a wide variety of unique metabolites^62^. These metabolites possess various biological activities and mediate interactions between microbes or between microbes and the host. Yet, a significant knowledge gap persists in understanding microbial ribosomal peptides’ diversity, abundance, prevalence, and role in maintaining microbial and human homeostasis. In this study, we investigated the biosynthetic potential of RiPPs within the human-associated microbiota, demonstrating their remarkable chemical diversity, novelty, and potential bioactivity. However, it is important to recognize that the scope of RiPP discovery is constrained by the availability of microbial genome datasets, predominantly from the gut microbiome, particularly bacteria. Concurrently, while utilizing a combination of deep learning-based and rule-based methods can help strike a balance between precision and novelty in studying RiPP biosynthesis, this strategy may result in a somewhat constrained depiction of the entire RiPP family from the human microbiome, as part of actual RiPPs may be excluded from the analysis. Nonetheless, our study thoroughly uncovers the profile of RiPP biosynthesis in the human microbiome and compiles an atlas of RiPPs for future research on innovative antimicrobial strategies and therapeutic interventions targeting microbiome^63,64^.

The ecological dynamics of the human microbiome are greatly shaped by specialized microbial metabolites that play protective roles and can potentially impact human health. Recent analyses of the human microbiome have revealed hidden potential for antibiotics, with RiPPs being the most commonly predicted compounds across various microbial environments within the body^13^. Concurrently, peptides with potentially harmful effects were also identified, including a lasso peptide from the oral bacterium *Rothia aeria* and a potentially toxic lantibiotic from translocating *Streptococcus*^65,66^. Our study observed that RiPPs are enriched in the human gut and oral microbiome, most of which are niche-specific bioactive molecules and actively transcribed in healthy human gut microbiomes. RiPPs with protective roles can mediate microbe-microbe interaction (e.g., antibacterial activities^13^, quorum-sensing or quorum-quenching^20^), or microbiome-host interaction (e.g., immunomodulatory activities^16-19^, cytotoxicity^67^). These interactions could influence the balance of the human-microbe holobiont and potentially substantially impact human health. We believe that these protective RiPPs, sourced from the human microbiome, represent innovative antimicrobial tactics and therapeutic interventions that are yet to be fully explored.

In microbe-microbe interactions, quorum-sensing is used for social coordination, often via peptide-mediated mechanisms in Gram-positive bacteria. RiPPs, particularly AIPs, can act as signaling molecules, regulate their production, and respond to quorum-sensing signals^10^. Our study identified six AIP families enriched in healthy microbiomes, most exhibiting significant antibiofilm activity against IBD-or CRC-related pathogens. Polymicrobial biofilms, particularly in IBD and CRC^68^, complicate treatment as antimicrobials need to target all biofilm pathogens. The combination of AIPs with antibiotics, specifically AIP-I, has shown potential in enhancing biofilm infection treatment^69,70^, as it can trigger MRSA biofilm dispersal and increase the susceptibility of detached cells to antibiotics. Our study revealed five AIPs with anti-pathogenic biofilm activity and found that two antibiofilm AIPs could adjust the gut microbial community in an *ex vivo* assay and hereby reduced pathogenic species linked to IBD, suggesting an alternative treatment option. These AIPs may assist in microbial communication through quorum-sensing or indirectly influencing the microbiome by controlling the production of antibacterial metabolites^50^. Considering their increased prevalence in healthy individuals compared to patients with IBD or CRC, our findings suggest that they could provide protective roles by inhibiting the growth or biofilm formation of pathogens^71^. While exploring AIPs’ impact on harmful biofilms and microbiome homeostasis could open up new therapeutic avenues, their exact mechanisms remain to be understood, and their bioactivity needs to be confirmed *in vivo* animal models.

Our study provides a comprehensive analysis of the biosynthetic landscape of RiPPs in the largely unexplored human microbiome, using (meta)genome mining and extensive omics analysis. In a proof of principle study, we linked RiPP profiles with various human diseases and pinpointed several RiPP candidates that may potentially impact human health. Despite certain limitations, our study offers valuable insights into the diversity and potential functions of RiPPs in the human microbiome. We also identified protective RiPPs that can combat pathogenic biofilms and possess the ability to maintain the balance of microbial communities, thus offering alternative antimicrobial tactics or therapeutic interventions that target the microbiome.

## Methods

### Biosynthetic gene cluster analysis and RiPP precursor identification from human microbiome reference genomes

We retrieved reference genomes of body-wide human microbes from two available datasets: 289,232 genomes from Unified Human Gastrointestinal Genome dataset (UHGG, v2.0^34,35^), and 17,249 genomes from CIBIO^33^ (**Supplementary Data 10)**. To normalize taxonomic annotations for all the genomes, we re-annotated all of them using GTDB-Tk^72^ (v2.0.0) against GTDB (rev207 version). All genomes were analyzed by antiSMASH 6.0^26^ for BGC detection using default parameters. To identify the RiPP precursor peptide, all open reading frames (ORFs) in each genome were annotated by prodigal-short^73^. Subsequently, small ORFs (≤150 amino acids) containing a start and stop codon and ribosome-binding site motifs were subjected to TrRiPP^25^ and DeepRiPP^24^ for RiPP precursor identification with default parameters. Notably, DeepRiPP can identify RiPP products even when the precursor genes are distant from the tailoring enzymes, while TrRiPP can detect RiPPs from highly fragmented metagenomes. Combining these tools can effectively identify more potential RiPP precursor peptides in metagenome-assembled genomes, which are predominant in our collection.

### Analysis of genomic context of predicted RiPP precursor peptides

For all identified RiPP precursors, we first checked whether they were within the region of antiSMASH-defined BGCs. For those precursors outside antiSMASH-defined BGC region, we defined 10 genes of the precursor upstream and downstream as genomic neighborhoods. We used two methods to examine the characteristics of these genomic neighborhoods associated with RiPP. First, we analyzed the protein domain of precursors and their 10 flanking genes downstream and upstream by a domain-based approach: RPS-BLAST. A domain was significantly assigned with a default CDD *e*-value threshold (< 0.01), and the protein aligns to at least 80% of the PSSM’s length. The assigned domain belonging to a dataset of known RiPP precursor domains and RiPP-related biosynthetic enzyme domains was noted as RiPP-related domains (**Supplementary Tables 1 and 2**). This database was collected and expanded based on the hmm-rule-parser of antiSMASH and previous studies^9,21,26,74^. For the remaining precursors, the domains were paired with a broader dataset of biosynthetic enzyme domains, which were used in the decRiPPter^36^ pipeline for exploring any novel biosynthetic enzyme candidates for RiPP.

### Chemical space comparison of RiPP precursor peptides

Known RiPP precursor peptides were collected from The Minimum Information about a Biosynthetic Gene cluster database (MIBiG 3.0)^75^. Extended-connectivity fingerprints (ECFPs) are used to predict and gain insight into RiPPs precursors’ chemical diversity^40^. Then ECFP6 fingerprints for each unique RiPP precursor were compared with each other to generate Tanimoto coefficient matrices^76^ and visualize the chemical space by uniform manifold approximation and projection (UMAP). Specifically, The ECFP6 chemical fingerprints of the domain representatives were calculated using The Chemistry Development Kit^77^. Subsequently, we generated a Jaccard distance matrix based on the chemical fingerprints and performed dimension reduction using densMAP^78^ with n_neighbors= 15, implemented in the UMAP Python package. Furthermore, we calculated the pairwise Tanimoto coefficient of each precursor using NumPy and scikit-learn. The average Tanimoto coefficient was obtained by averaging all of these coefficients. Next, core precursor sequences predicted by the cleavage predictions module of DeepRiPP were used to explore the chemical diversity of nine RiPP classes. Pairwise Tanimoto coefficient was further adopted for measuring the similarity between precursors. The precursor with the highest median within-class Tanimoto coefficients was chosen as representative structures to generate Tanimoto coefficient matrices for intra-classes and further calculate the diversity for intra-classes. Tanimoto score ranges from 0 to 1. The higher Tanimoto score means higher chemical similarity between two precursors or two subclasses.

### Novelty examination of RiPP precursor families

To trade off the novelty and accuracy, 423,831 RiPP precursors that were either within RiPP BGCs region^38^ identified by enzymes-oriented approach or identified by both two precursors-centric approaches were retained for further analyses. These precursor peptides were grouped by MMseqs2^79^ with the following parameters: easy-cluster clusterRes tmp --min-seq-id 0.5 --single-step-clustering --cluster-mode 2 --cov-mode 2 -c 0.95. The precursors within a family are more likely to share a similar function^80^. The classification of each RiPP precursor family was further determined based on the novelty of their precursor and their genomic contexts **(Supplementary Information)**. The novelty of RiPP families was further classified into three types: (1) “classic RiPP families”, which are identified by having precursors that exhibit similarity to known RiPP precursors and are located in a genomic context associated with typical RiPP biosynthetic gene clusters. (2) “Novel RiPP families”, which consist of novel precursors and/or genomic neighborhoods containing novel biosynthetic genes. (3) “Others”.

### Bioactivity prediction of RiPP families

We utilized the DeepBGC^27^ tool to identify the potential bioactivity of RiPP precursors. This approach allowed us to assess and classify the potential bioactivity of RiPPs based on the predicted functional genes within their genomic neighborhoods. Our approach involved collecting gene regions within 10 genes upstream and downstream of RiPP precursor genes, which we referred to as genomic neighborhoods. We then utilized DeepBGC to predict the potential function of each RiPP. This tool could account for the four most common compound activity classes: antibacterial, cytotoxic, inhibitor, and antifungal. Precursors exhibiting multiple predicted activities were labeled as “multiple”, while the activity of each RiPP family was determined by considering the activities of over 50% of the precursors within the family.

### Analysis of metagenomic and metatranscriptomic data

The metagenomic and metatranscriptomic samples were downloaded from sequence read archive (SRA) of the NCBI (**Supplementary Table 3**). Raw metagenome and metatranscriptome sequencing reads were quality filtered using *bbduk*.*sh* with the following parameters: qtrim=rl ktrim=r mink=11 trimq=10 minlen=40. The resulting metatranscriptomic reads were subjected to *bbmap*.*sh* for removing reads derived from ribosomal RNAs. To remove the human host contamination, the high-quality metagenomic and metatranscriptomic sequencing reads were further searched against the human reference genome (GRCh38.p13) from NCBI, and unmapped reads were mapped to microbe reference genomes using BWA (0.7.17-r1188)^81^ with default parameters. The reads mapped to RiPP precursor gene were counted by featurecounts^82^ with the following parameters: -f -t CDS -M -O -g transcript_id -F GTF -s 0 -p --fracOverlap 0.25 -Q 10 -primary. In metatranscriptomic datasets, the count file with the absolute abundance for each family was imported, and differential gene abundance between healthy and diseased subjects was normalized and analyzed by implementing DESeq2 pipeline in R. Of note, the abundance of each family was calculated by the sum of the abundances of all genes in the family. Additionally, MetaPhlAn 4^83^ was used to compute the relative abundance of microbial species. In addition, accumulation curves, alpha-diversity, beta diversity and PERMANOVA were performed using the R package *vegan*. Beta diversity was performed to quantify the relative abundance differences in the overall composition of RiPP precursor families between the disease and the control groups. PERMANOVA was performed to show the encoding and expression profile differences of RiPP precursor families between disease and control groups. Specifically, families with a prevalence ≥ 5% in a cohort were subjected to further analysis (Beta diversity, differential analysis, and classifier) for investigating potential causality in human disease. In metagenomic datasets, we calculated the significance of prevalence using a two-sided Fisher’s exact test, *p* < 0.05. BiG-SCAPE^84^ was used to visualize the genomic context of differential RiPP precursors families. Besides, only 50 members with formative genomic context (larger gene sizes) in RiPP families with larger members (≥ 50 genes) were chosen for analysis. Each representation biosynthetic gene cluster was chosen to show the conserved domain and products in each family. Multiple sequence alignment of all precursor families was conducted using MAFFT^85^, followed by Jalview for visualization.

### Chemical synthesis of AIPs

To begin, we initiated the calculation and analysis of the precursor-AgrB complex (**Supplementary Table 4**). The calculation was performed using the following command: colabfold_batch --amber --templates --num-recycle 3 --use-gpu-relax --model-type alphafold2_multimer_v3 input_path output_path. The resulting structures were visualized using PyMol. During our analysis of the reported AIP precursor and paired AgrB, we observed that the binding of the Precursor-AgrB complex can be partially buried. In other words, the core peptide of the precursor should be fully accommodated within the catalytic pocket of AgrB. Based on this observation, we manually examined the predicted complexes. Specifically, we focused on two aspects: 1. We inspected whether the conserved C-terminal core region is captured and effectively buried within the catalytic pocket of AgrB. By conducting this analysis, we aimed to gain insights into the interaction between the precursor and AgrB, shedding light on the potential core peptide of AIP precursors. Next, we aimed to comprehensively assess the conservation patterns of precursor sequences within each AIP family, incorporating both our study data and relevant sequences available in public databases. Thus, we first collected nonredundant precursor sequences from our study. Additionally, we included similar sequences (>50% similarity) obtained from NCBI as additional sources of AIPs found in nature. Next, Logo sequences for each clustered short peptide family were generated by makelogo.py, which is available at https://github.com/yxllab-hku/cluser_to_logo/blob/main/makelogo.py. With the bioinformatic prediction, we deduced the potential mature chemicals of six AIP families and chemically synthesized 9 potential AIPs **(Supplementary Information)**.

### Antibiofilm assay

Biofilm formation was assessed using the crystal violet method. The strains listed in **Supplementary Table 5** were employed as the tested strains. To initiate the experiment, the inocula from overnight cultures were diluted at a ratio of 1:100 using the corresponding culture medium. To perform dose-response assays, the chosen peptide was prepared as a stock solution and serially diluted in DMSO. Following this, 1 μL of each concentration was carefully dispensed into the corresponding wells. Subsequently, 99 μL of the bacterial culture, diluted accordingly, was added to the respective wells. Additionally, vehicle controls consisting of 1 μL of DMSO were included to establish the baseline biofilm formation of the strains. The microtiter plate was then incubated under appropriate conditions. Following the incubation period, the biofilm was gently washed with PBS to remove any unattached cells and then allowed to air dry. Next, 100 μL of a 0.1% crystal violet solution was added to each well to stain the biofilm. The plate was further incubated at room temperature for 15 minutes to ensure proper staining. After the incubation, each well was washed three times with water to completely remove the dye. The plate was then inverted and allowed to air-dry at ambient temperature throughout the night. To quantify the biofilm formation, 100 μL of a 30% acetic acid solution was added to each well to dissolve the crystal violet. Following an additional incubation period of 10-15 minutes at room temperature, the absorbance was measured at 550 nm. Three replicated wells were used for each group to ensure accuracy and reproducibility.

### Anti-inflammatory activity by ELISA experiment

The RAW264.7 cells were seeded at a density of 15 × 10^4^ cells/well in 24-well plates and cultured at 37 °C and 5% CO2 overnight. Cells were treated with AIPs (1 μg/mL and 0.1 μg/mL) or 25 μg/mL of Dexamethasone for 3 hours and then stimulated with 1 μg lipopolysaccharides (LPS) for 24 hours. The secretion of TNFα and IL6 was measured according to the ELISA kit (Elabscience). ELISA methods were applied according to the manufacturer’s instructions without any modifications by using mouse IL-6 (Interleukin 6) ELISA Kit (E-EL-M0044), mouse TNF-α (tumor necrosis factor alpha) ELISA Kit (E-EL-M3063). Optical densities were read on a plate reader set at 450 nm a microplate reader (BioTerk, Winooski, VT, USA). The concentration of each parameter in the samples was calculated from the standard curve, multiplied by the dilution factor and was expressed as mean ± standard error of the mean (SEM).

### *Ex vivo* screening study

#### Mice

Mouse studies were performed in accordance with all relevant ethical regulations and were approved by the Ethics Committee of the Animal Experimental Center of ZhuJiang Hospital, Southern Medical University (LAEC-2022-059). IL-10-deficient (IL-10^−/−^) male mice with C57BL/6N background were purchased from Cyagen Biosciences (Guangzhou, China). All the mice were housed in specific pathogen free (SPF) conditions with a 12 hours light/dark cycle, and were provided sterilized water and food ad libitum (temperature 23 ± 2°C, humidity 45 ± 5%). IL-10-deficient mice were used to model spontaneous chronic colitis, which closely mimics human inflammatory bowel disease (IBD)^86^.

#### Feces collection

Fecal samples were collected from 10 IL-10-deficient C57BL/6J male mice (16 weeks old), with an average of 7-8 droppings per mouse. The collected pellets were combined and resuspended in 65 mL of a rich medium mGAM. Following resuspension, the tubes were gently centrifuged at 1,000 rpm for 2 minutes, and the supernatants were then retained for further assay.

Two selected AIPs were pre-dissolved in DMSO to final concentrations of 0.1 g/mL, 1 g/mL, and 10 g/mL. Subsequently, 1 µL of AIPs with varying concentrations and 49 µL of fecal supernatant were inoculated into 950 µL of mGAM media, resulting in a final volume of 1 mL. The mixtures were then anaerobically incubated at 37°C for 48 hours. Following the incubation period, the bacterial growth media were centrifuged at 12,000 rpm for 10 minutes. The supernatants were carefully removed, and the resulting pellets were washed twice with 1 mL of PBS before being collected for DNA extraction. The pelleted samples were extracted using the QIAamp® DNA Micro Kit (Qiagen) according to the manufacturer’s instructions. Finally, all DNA samples were prepared for shotgun metagenomics sequencing.

#### Metagenomic sequencing

All DNA samples were submitted to Novogene for shotgun metagenomics sequencing using a 150bp paired-end protocol. Initially, 1 µg of DNA per sample was fragmented via sonication to achieve a size of 350bp. Subsequently, the fragmented DNA underwent end-polishing, A-tailing, and ligation with a full-length adaptor for library construction. The constructed libraries were assessed for quality and quantity using Qubit and real-time PCR for quantification, as well as a bioanalyzer for size distribution analysis. The quantified libraries were then pooled and sequenced on the Illumina NovaSeq X Plus platform, generating approximately 10.7 Gb of data per sample.

#### Metagenomic analysis

Raw reads from metagenomic sequencing were processed using Fastp v0.21.1 for quality control, with parameter of “--detect_adapter_for_pe -l 50 -5 3 -3 3”. High-quality metagenomic sequencing reads were further subjected to KneadData (https://github.com/biobakery/kneaddata) for detecting and removing reads belonging to the human genome, though searching against the mouse reference genome (GRCm39) from GENCODE^87^. MetaPhlAn version 4.1.1^83^ was used to generate taxonomic profiles of metagenomes.

Alpha diversity, as measured by Shannon diversity, and beta diversity, evaluated through Bray-Curtis dissimilarity, were computed for species-level taxonomic profiles utilizing the Vegan package in R^88^. Differential abundance analysis between groups was conducted using MaAsLin2^53^ (microbiome multivariable associations with linear models). Species with a false discovery rate (FDR)-adjusted *p*-value of less than 0.05 were deemed significantly different between the two groups.

## Data availability

Raw read sequences of the shotgun metagenomic sequences were deposited at Sequence Read Archive (SRA, NCBI) under accession of PRJNA1166984. Reference genomes of body-wide human microbes from two available datasets: 289,232 genomes from Unified Human Gastrointestinal Genome dataset (UHGG, v2.0, https://ftp.ebi.ac.uk/pub/databases/metagenomics/mgnify_genomes/human-gut/v2.0/), and 17,249 genomes from CIBIO (http://segatalab.cibio.unitn.it/data/Pasolli_et_al.html). The genomes used in this study are provided in **Supplementary Data 10**. The raw data for omics data analysis were collected from NCBI datasets through the accession numbers provided in **Supplementary Table 3**.

## Code availability

Codes related to the analyses in this study are available at https://github.com/ZHANGJianArya/RiPPs_human-microbiome.

## Acknowledgments

The authors would like to thank Zhiman Song and Cunlei Cai for their help in MS and NMR analysis.

## Funding

This work is partially funded by three Hong Kong Research Grants Council General research grants (HKU27107320, HKU17115322, and HKU17102123).

## Author contributions

Conceptualization: Y.-X.L., J.Z., DW.Z., Bioinformatics analysis: J.Z., DW.Z., Y.G., Z.Z, B.B.H. Experimental validation: J.Z., Y.X., J.L.Z., R.-Z.L., Y.G., Y.Q.S., P.Y.C, X.C.L., H.-W.Z. Visualization: J.Z., DW.Z., Writing—original draft: J.Z. Writing—review & editing: Y.-X.L., D.W.Z. and J.Z. Investigation: Y.-X. L. and D.W.Z. Supervision: M.-X.C., Y.-X. L.

## Competing interests

The authors declare that they have no competing interests.

## Notes

### Competing Interest Statement

The authors have declared no competing interest.

### Summary of Updates

1. Added new result:AIPs affect the composition of IBD mouse fecal-derived ex vivo microbial community 2. Added author information who participated in: AIPs affect the composition of IBD mouse fecal-derived ex vivo microbial community"; Supplemental files updated. 3. Supplemental files updated

