## Supplementary information and Supplementary figures for "Comprehensive Biosynthetic Analysis of Human Microbiome Reveals Diverse Protective Ribosomal Peptides"

Supplementary Figure 22 NMR assignment of BF_94_et_L1 ... 39
Supplementary Figure 23 UPLC-HRMS/MS analysis of synthesized BF_94_et_free ... 40
Supplementary Figure 24 NMR assignment of BF_94_et_free ... 42
Supplementary Figure 25 UPLC-HRMS/MS analysis of synthesized BF_280_t ... 43
Supplementary Figure 26 NMR assignment of BF_280_t ... 45
Supplementary Figure 27 UPLC-HRMS/MS analysis of synthesized BF_280_c ... 45
Supplementary Figure 28 NMR assignment of BF_280_c ... 46
Supplementary Figure 29 UPLC-HRMS/MS analysis of synthesized BF_398_et_L1 ... 47
Supplementary Figure 30 NMR assignment of BF_398_et_L1 ... 48
Supplementary Figure 31 UPLC-HRMS/MS analysis of synthesized BF_398_c ... 49
Supplementary Figure 32 NMR assignment of BF_398_c ... 50
Supplementary Figure 33 UPLC-HRMS/MS analysis of synthesized BF_488_et_free ... 51
Supplementary Figure 34 NMR assignment of BF_488_et_free ... 53
Supplementary Figure 35 UPLC-HRMS/MS analysis of synthesized BF_598_c ... 54
Supplementary Figure 36 NMR assignment of BF_598_c ... 55
Supplementary Figure 37 Potential quorum sensing or quorum quenching function of AIPs in mediating biofilm formation ... 56
Supplementary Figure 38 AIPs inhibit clinical pathogenic biofilm formation ... 57
Supplementary Figure 39 Relative abundance at the bacterial taxonomic levels of different treatments ... 59
Supplementary Figure 40 Coordinates of PCo1 and PCo2 axes of PCoA plot ... 60
Supplementary Figure 41 Differentially abundant species between blank and treatment groups ... 61

**Reference ... 62**

### Supplementary Information

#### Supplementary Notes

##### Taxonomic distribution of prioritized RiPP

To uncover additional phylogenetic trends, we inspected the taxonomic distribution of detected RiPPs. A strong enrichment of RiPP precursors was observed in 21 phyla and 3,369 species. Given genus is an appropriate taxonomic rank for comparison by their stably representative secondary metabolite biosynthetic diversity[1], we specifically examined RiPPs biosynthetic capacity in 1,014 genera based on the average number of RiPPs identified in each of them. We found that RiPP biosynthetic potential at the genus level considerably varied from 1 to 33 RiPP precursors per genome, with *Elizabethkingia* (n= 33 RiPP precursors), *Chryseobacterium* (n= 18 RiPP precursors) and *Tissierella* (n= 15 RiPP precursors) having the highest number of RiPPs **(Supplementary Fig 4)**. Of note, 285 genera (284 bacteria and 1 archaea) contain 2 precursors per genome. Besides, 10 genera harbor diverse RiPP with more than eight RiPP precursor classes, including *Phocaeicola*, *Lachnospira*, *Agathobacter*, *Parabacteroides*, *Roseburia*, *Bacteroides*, *Prevotella*, *Bifidobacterium*, *CAG-110*, and *Dialister*. However, 332 genera only encode one type of RiPP precursor. Overall, RiPP precursors are widely distributed across a multitude of taxa from the human microbiome.

##### Novelty of the RiPP precursor in each family

**Refer to Figure 2e and 2f. e** The chord diagram illustrates the novelty of identified precursor families (left panel) for nine different RiPP classes (right panel). Left panel: The "MIBiG" category represents precursor families that exhibit homology to experimentally characterized RiPP precursors deposited in the MIBiG database. Precursor families that contain well-defined RiPP-associated domains are highlighted in black. These domains include AgrD, rSAM-modified RiPPs-associated domains, and other domains such as 1, Graspetide-associated (3 families), 2, Lanthipeptide-associated (31 families), 3, Lassopeptide-associated (26 families), 4, Thiopeptide-associated (5 families), 5, RiPP like-associated (25 families), 6, LAP-associated (19 families), and 7, other known RiPPs-associated (2 families). Those precursor families that do not belong to the above categories as "Novel RiPP". The numbers in brackets represent the number of precursor families within each category. **f** The number and proportion of RiPP families under each condition integrating RiPP precursor and genomic neighborhood. The novelty of RiPP families was further classified into classic RiPP, Novel

RiPP, and Others. The classic RiPP category includes families with known precursor homology and a well-defined genomic neighborhood. The Novel RiPP category includes families with no known precursor homology and/or with a novel genomic neighborhood. The Others category includes families that do not fit into either of the above categories.

#### Experiment materials and methods

##### Phylogenetic tree construction.

The representative genome of each genus was used to generate maximum-likelihood trees via GTDB-Tk (v2)[2] with the following parameters (refer to the GTDB-Tk user guide: https://ecogenomics.github.io/GTDBTk/commands/index.html): 1. GTDB-Tk reference data release 207 was used. 2. Parameters of classify: gtdbtk classify_wf --genome_dir --out_dir --force. 3. Parameters of infer tree from multiple sequence alignment: gtdbtk infer --msa_file MSA_FILE --out_dir OUT_DIR. 4. Using the convert_to_itol command to make the tree suitable for visualization in iTOL: gtdbtk convert_to_itol --input_tree --output_tree. Finally, we annotate their phylogenetic trends using iTOL[3].

##### Novelty examination of RiPP precursor families

To trade off the novelty and accuracy, 423,831 RiPP precursors that were either within RiPP BGCs region[4] identified by enzymes-oriented approach or identified by both two precursors-centric approaches were retained for further analyses. These precursor peptides were grouped by MMseqs2[5] with the following parameters: easy-cluster clusterRes tmp --min-seq-id 0.5 --single-step-clustering --cluster-mode 2 --cov-mode 2 -c 0.95. The precursors within a family are more likely to share a similar function[6]. The classification of each RiPP precursor family was further determined based on the novelty of their precursor and their genomic contexts **(Supplementary Fig. 5)**. (1) Novelty of precursors. The known RiPP precursors collected from MIBiG 3.0[7] were quired against 12,076 representative precursor sequences by BLASTp. A significant hit was considered with an e-value < 0.05, the alignment spans ≥ 90% of the peptide, and the length of the hit was 90%–110% of the length of the known RiPP precursor peptide. For the remaining families, each family was assigned a “domain representative” if at least 80% of its members contained a specific RiPP precursor domain. Otherwise, the family was classified as a novel family. (2) Novelty of genomic context of precursors. To assess the novelty of the genomic context surrounding RiPP precursors, we employed antiSMASH or DeepBGCs[8] to identify the presence of biosynthetic genes. Our analysis began with individual precursors, examining their genomic neighborhoods. If a precursor's genomic context was found to be associated with a known RiPP BGC, it was categorized as "RiPP". If the genomic context was novel recognized by DeepBGC, it was categorized as "Novel". Alternatively, if the genomic neighborhood showed evidence of hybridization with other secondary metabolite BGCs or exhibited other conditions, it was placed in the "Others" category. Furthermore, to provide a more comprehensive classification, we grouped and classified the types of genomic neighborhoods for each RiPP precursor family. If over 50% of the family members belonged to a specific type of genomic neighborhood, the entire family was assigned to that respective type. This allowed for a more refined understanding of the genomic diversity within each RiPP precursor family. (3) Novelty of RiPP families. The novelty of RiPP families was further classified into three types: The first type is referred to as "classic RiPP" families, which are identified by having precursors that exhibit similarity to known RiPP precursors and are located in a genomic context associated with typical RiPP biosynthetic gene clusters. The second type is "Novel RiPP families", which consist of novel precursors and/or genomic neighborhoods containing novel biosynthetic genes. Lastly, the third type is "Others".

**Statistical modeling for classifier**

To further investigate whether the abundance of RiPP precursor families can discriminate patients from health control, the relative/normalized abundance was further subjected to the cross-validated random forest classification model, and the receiver operating characteristic (ROC) curve for the predictive disease model was calculated. The average area under the ROC curve value was used to evaluate the performance of the classifier. A higher average AUROC represents the stronger predictive ability of the model to distinguish one group from another group. Following we further rank individual translated RiPP precursor peptide families' contribution to the predictive accuracy of the model[9].

**Chemical synthesis of AIPs**

With the bioinformatic prediction, we deduced the potential mature chemicals of six AIP families **(Fig.4b, Supplementary Table 4)**. Specifically, these substances typically give rise to a lactone (BF_63, BF_94, and BF_488) or thiolactone (BF_398, BF_280, BF_598) between a Ser or Cys residue and another amino acid with a hydrophobic side chain. Of note, it is plausible that mature products of BF_280, BF_598, and exocyclic region-free BF_398 families could lead to the creation of cAIPs (**Fig. 4d, Supplementary Fig. 18**). Furthermore,

BF_94 exhibits sequence similarity to the previously reported exocyclic region-free AIP ( [SFFIF] ) from *C. thermocellum* Clo1313_2818[10]. We therefore synthesized a homologous variant, BF_94_et_free, for the BF_94 family to further investigate the potential bioactivity of AIPs. Finally, BF_94_et_L1, BF_280_c, BF_398_c, BF_598_c were chemically synthesized by Sangon Biotech (Shanghai, China) and the five predicted AIPs of BF_63_et_L1, BF_94_et_free, BF_280_t, BF_398_et_L1, BF_488_et_free were chemically synthesized by Yuan peptide (Nanjing, China).

**LC-HRMS analysis of synthesized peptides**

For LC-HRMS analysis, the samples were desalted and prepared at a final concentration of 0.01 mg/mL. LC-MS analysis was carried out using a Waters ACQUITY UPLC BEH C18, 130Å, 1.7 µm column, coupled with a Thermo Scientific UltiMate 3000 UHPLC system and Bruker impact Mass Spectrometer. The LC method employed a gradient for chromatographic separation, starting at 5% B for 2 minutes, followed by 5% to 95% B for 15 minutes, and finally held at 95% for 4 minutes. The column was re-equilibrated to 5% B for 1 minute before the next run started. The column was maintained at 40˚C, and the flow rate was set at 0.2 mL/min, using 0.1% formic acid in $H_2O$ as solvent A and 0.1% formic acid in acetonitrile as solvent B. The MS system was tuned using a sodium formate standard, and all samples were analyzed in positive polarity with m/z range from 150 to 1500 Th, using data-dependent acquisition mode.

**NMR analysis**

$^1$H NMR and $^1$H-$^1$H COSY spectra were acquired on a Bruker Avance 600 MHz spectrometer with Cryoprobe, using dimethyl sulfoxide-$d_6$ as solvent. The dimethyl sulfoxide-d6 chemical shifts were used as the internal reference.

**Visualize the genomic neighborhood of each family**

First, in each family, 10 genes away from the small genes encoded precursors were extracted and annotated using prokka[11]. Then the architectures were predicted using BiG-SCAPE.[12] Besides, only 50 members with formative genomic context (larger gene sizes) in RiPP families with larger members (≥ 50 genes) were chosen for analysis. Each representation biosynthetic gene cluster was chosen to show the conserved domain and products in each family. Multiple sequence alignment of all precursor families that share amino acid sequence homology with the family was conducted using MAFFT[13], and shown using Jalview.

#### Supplementary Figures

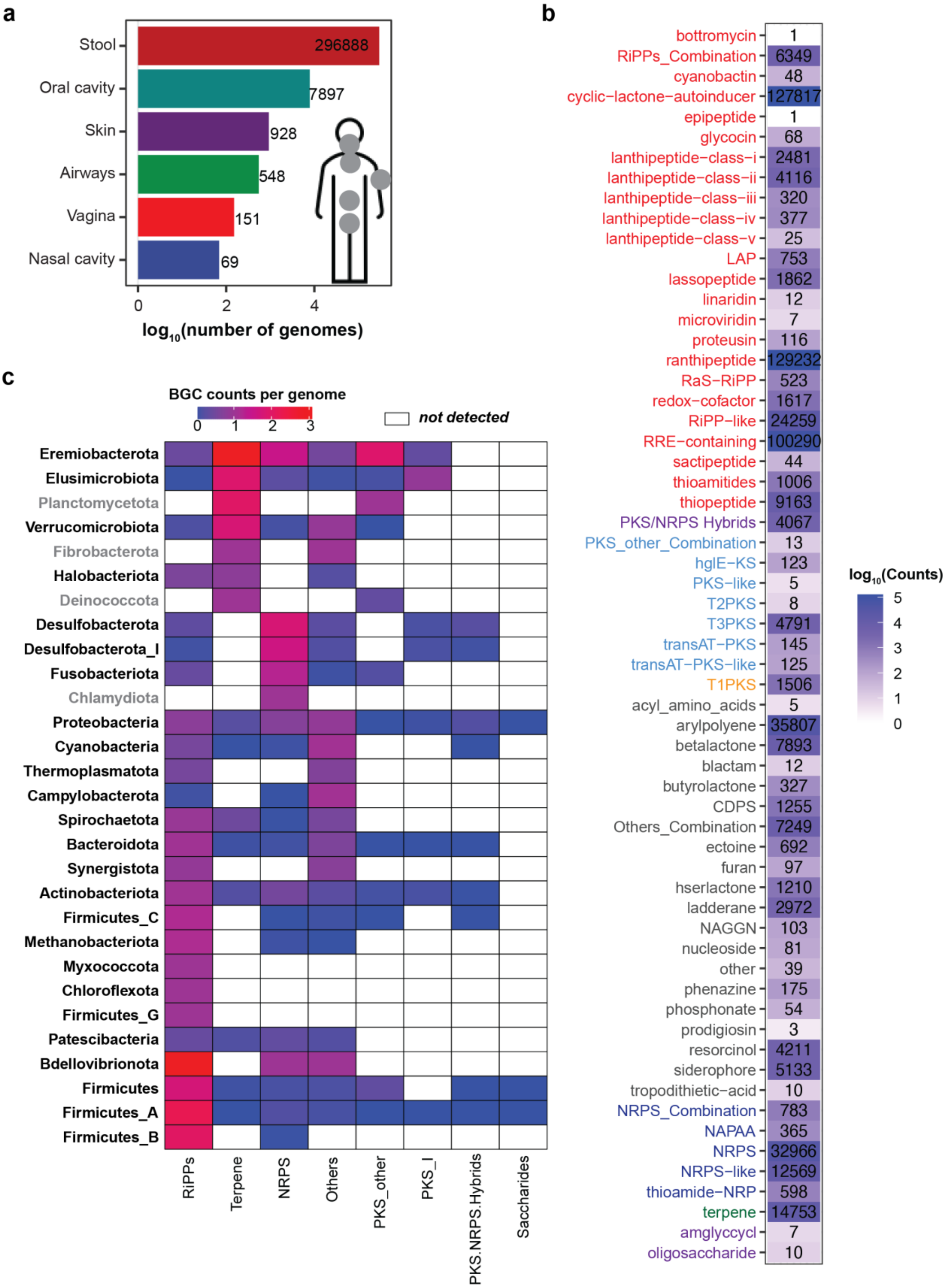

#### Supplementary Figure 1 Overview of BGCs detected by rule-based antiSMASH

**a** The distribution of genomes in human body sites. **b** The heat map displays the number of each BGC type identified from human gut bacteria, with different BGC types depicted in various colors corresponding to their respective BGC classes. **c** Heatmap showing the RiPP BGC counts per genome across various microbial phyla, categorized by BGC classes. For clarity, the RiPP BGC counts have been logarithmically transformed ($log_{10}$). Taxonomic classification was determined according to annotations from The Genome Taxonomy Database (GTDB). Notably, the species that encoded RiPPs BGCs were highlighted in black and bold.

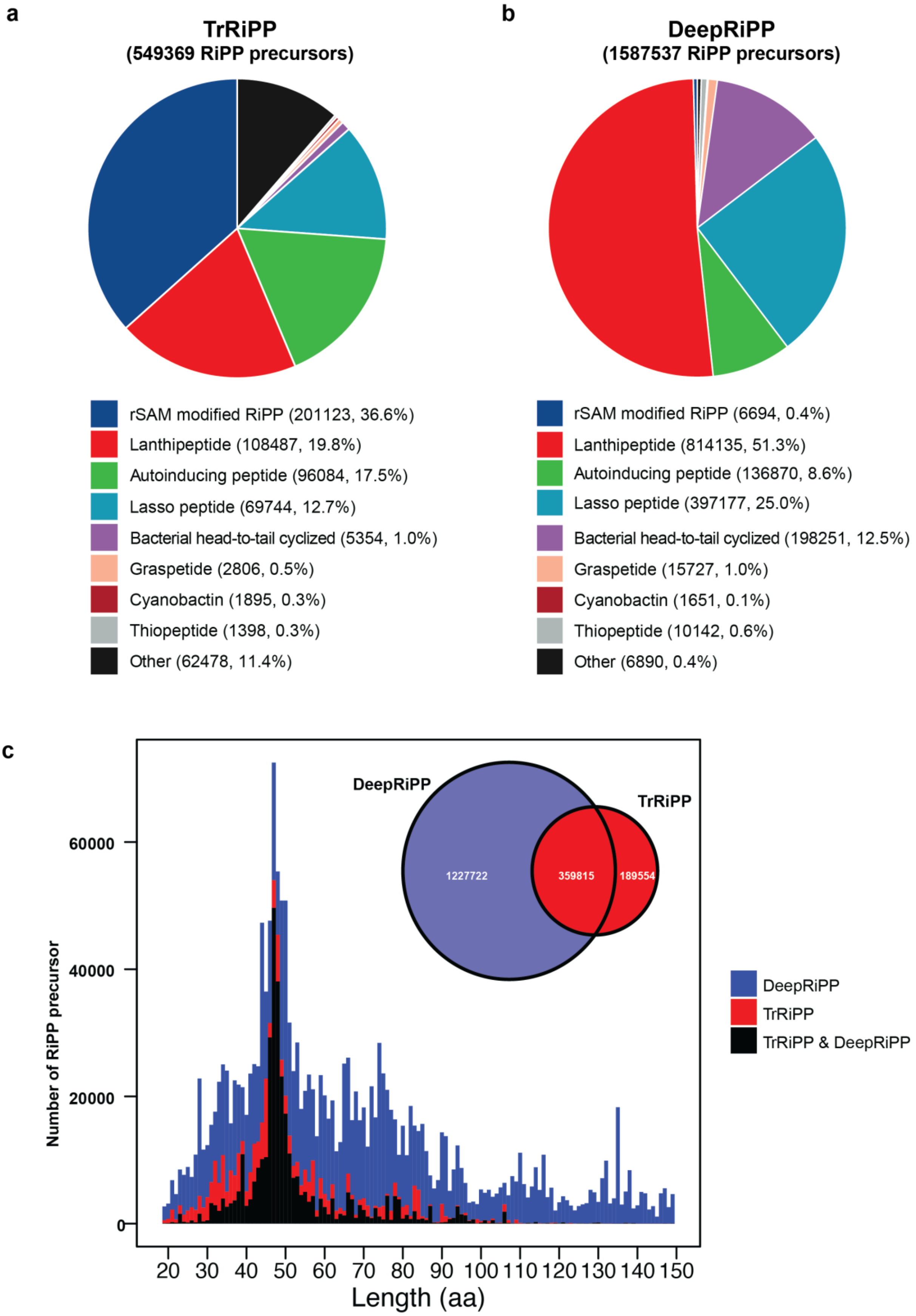

a
TrRiPP
(549369 RiPP precursors)
rSAM modified RiPP (201123, 36.6%)
Lanthipeptide (108487, 19.8%)
Autoinducing peptide (96084, 17.5%)
Lasso peptide (69744, 12.7%)
Bacterial head-to-tail cyclized (5354, 1.0%)
Graspetide (2806, 0.5%)
Cyanobactin (1895, 0.3%)
Thiopeptide (1398, 0.3%)
Other (62478, 11.4%)
b
DeepRiPP
(1587537 RiPP precursors)
rSAM modified RiPP (6694, 0.4%)
Lanthipeptide (814135, 51.3%)
Autoinducing peptide (136870, 8.6%)
Lasso peptide (397177, 25.0%)
Bacterial head-to-tail cyclized (198251, 12.5%)
Graspetide (15727, 1.0%)
Cyanobactin (1651, 0.1%)
Thiopeptide (10142, 0.6%)
Other (6890, 0.4%)
c
DeepRiPP
TrRiPP
1227722
359815
189554
Number of RiPP precursor
60000
40000
20000
0
20 30 40 50 60 70 80 90 100 110 120 130 140 150
Length (aa)
DeepRiPP
TrRiPP
TrRiPP & DeepRiPP

#### Supplementary Figure 2 Overview of RiPP precursors detected by two precursor-centric approaches

**a** Pie chart illustrating the distribution of RiPP precursors identified by TrRiPP. **b** Pie chart showing the distribution of RiPP precursors identified by DeepRiPP. The numbers within parentheses represent the respective counts and percentages of each RiPP class. **c** Inner: Venn diagram depicting the intersection of RiPP precursor peptides identified by DeepRiPP and TrRiPP. Outside: The length distribution of precursors.

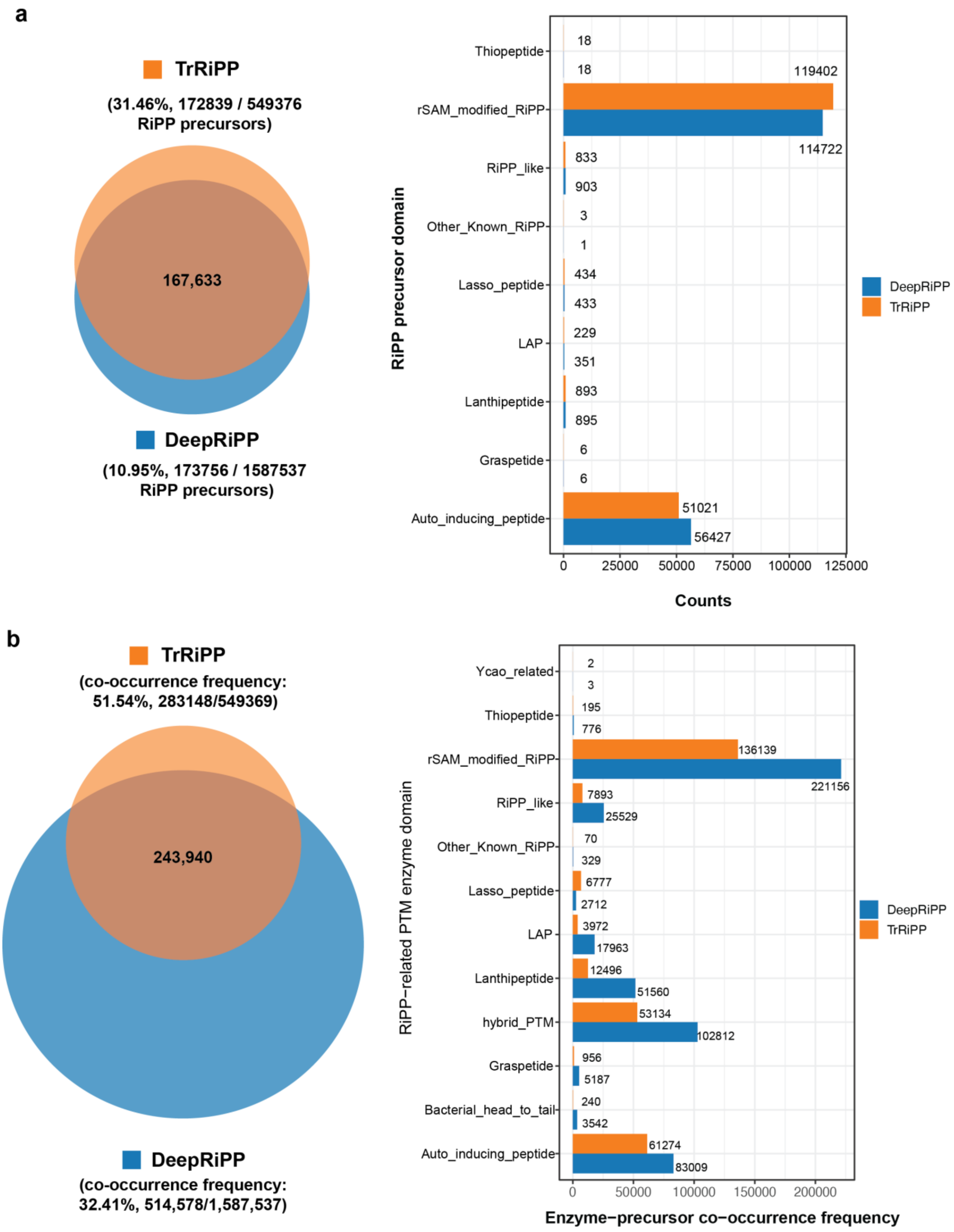

**Supplementary Figure 3 Domain annotation and genetic context annotation of RiPP precursors**

**a** Venn diagram (left) depicting the intersection of RiPP precursors with known RiPP precursor peptide domain by querying against the CDD database. In total, 172,839 precursors detected by TrRiPP and 173,756 detected by DeepRiPP can be assigned to the known RiPP precursor peptide domain in the CDD database, with 167,633 precursors being overlapped. Barplot (right) displays the respective number of RiPP precursors with known domains for each RiPP class. **b** Frequency of co-occurrence between reported RiPP post-modification enzymes domain and predicted precursors within ten downstream and upstream genes of precursors**.** Left panel**:** Total frequency of co-occurrence of PTM enzymes domain with predicted precursors predicted by TrRiPP (highlighted in orange) and DeepRiPP (highlighted in blue). Of note, the frequency of well-known RiPP post-modification enzyme domain co-occurring with RiPP precursors is 51.54% (283,148/549,369) and 32.41% (514,578/1,587,537) for precursors predicted by TrRiPP and DeepRiPP, respectively. Right panel: Identified RiPP PTM enzyme domains were classified into 12 families according to the classification of the final product in which the enzyme is involved in the modification. Notably, hybrid RiPP modification enzyme domains encoded by genes near the predicted RiPP precursor genes are more than 18% (hybrid_PTM).

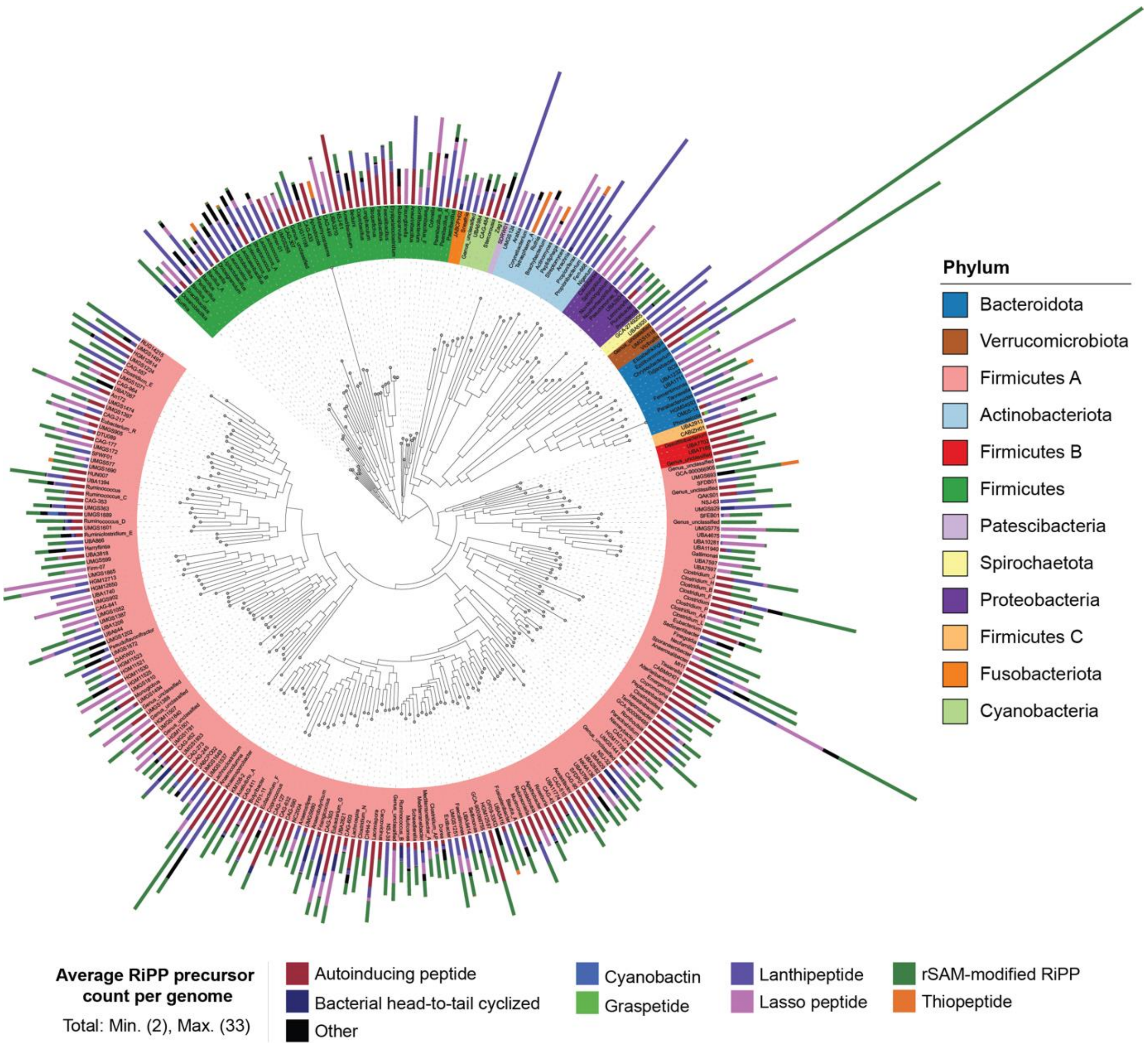

#### Supplementary Figure 4 Taxonomic distribution of RiPP precursors

The figure compares RiPP biosynthetic diversity among 284 bacterial genera containing at least two RiPP precursors per genome. The layers from inner to outer are: (1) The innermost layer shows the maximum likelihood phylogenetic tree of 284 representative bacterial genera associated with the human microbiome. Taxonomic classification is based on the GTDB. (2) The next layer illustrates the average representation of each RiPP subclass encoded within each genus per genome. This layer provides an overview of the RiPP biosynthetic potential within each bacterial genus.

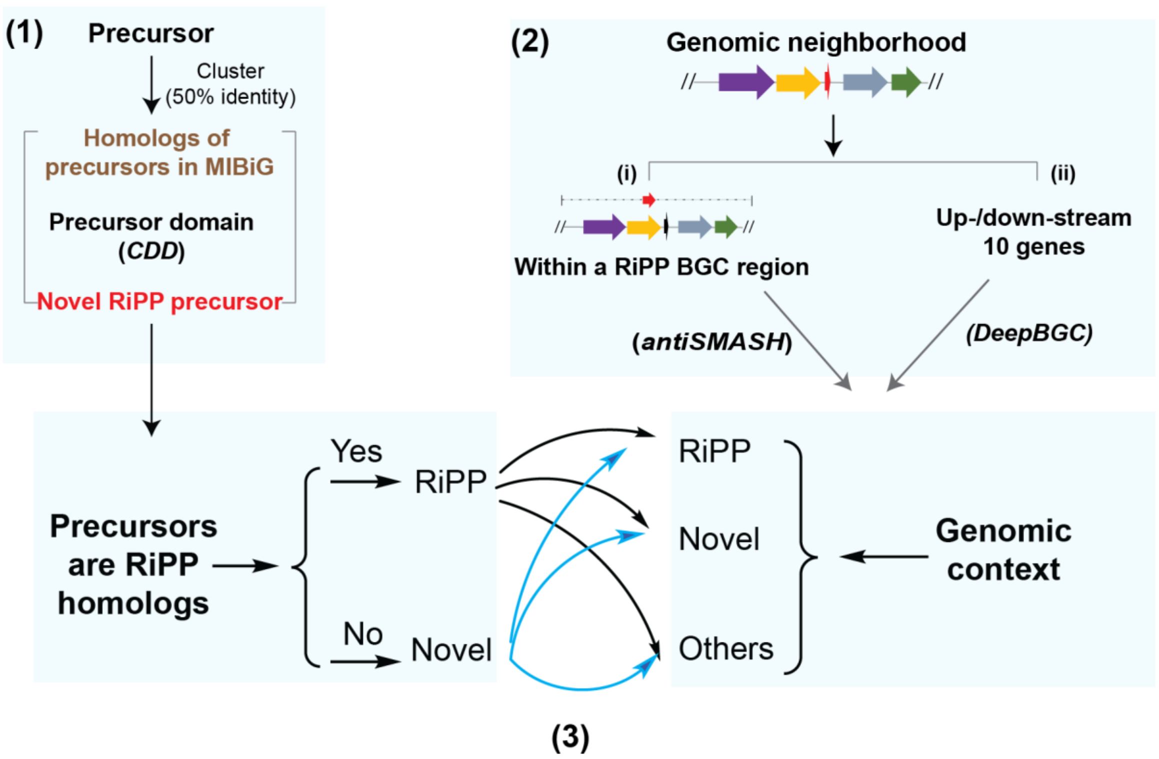

**Supplementary Figure 5 Pipeline for determining the novelty of RiPP families based on precursor and genomic context**

The pipeline involves determining (1) the homology of precursors, (2) analyzing the genomic neighborhood using antiSMASH and DeepBGC, and (3) integrating the information to classify the RiPP precursor families. (1) To determine whether the RiPP precursors have homology to known RiPP precursors (known RiPP) or not (novel). (2) It is important to note that 62.4% of RiPP precursors were found to be located within RiPP BGC regions identified by antiSMASH **(Fig. 2)**. To gain insights into the remaining subset of precursors (37.6%) and their genomic context, we employed an additional approach DeepBGC[8]. DeepBGC utilizes a machine learning algorithm to extrapolate and identify unknown BGCs, thereby complementing the information obtained from antiSMASH. The genomic neighborhood of the precursors (10 genes upstream and downstream) could be captured by DeepBGC. The genomic neighborhoods for each RiPP precursor family were further grouped and classified. (3) Finally, the classification of RiPP precursor families is determined by integrating the novelty of the precursor sequence and the genomic context.

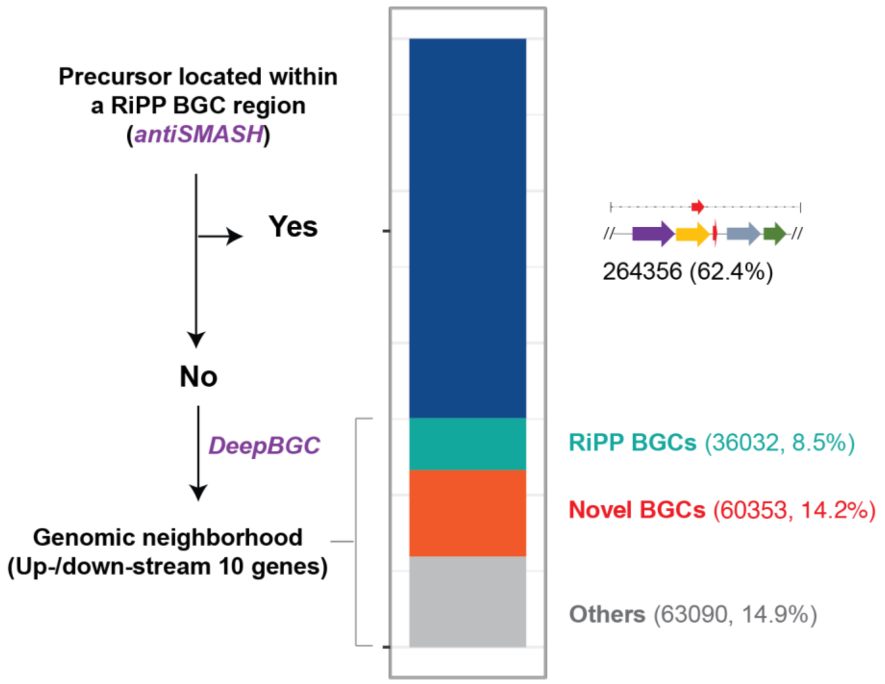

**Supplementary Figure 6 Genomic context of RiPP precursor**

The proportion of RiPP precursor located in a specific genomic context. In detail, 62.4% of RiPP precursors are within a RiPP BGC region identified by antiSMASH. We obtained upstream and downstream ten genes of the precursor as genomic neighborhoods for the remaining precursors and predicted their biosynthetic gene clusters using DeepBGC. Precursors located in the genomic context contain RiPP BGCs (8.5%), novel BGCs (14.2 %), and others (14.9%)

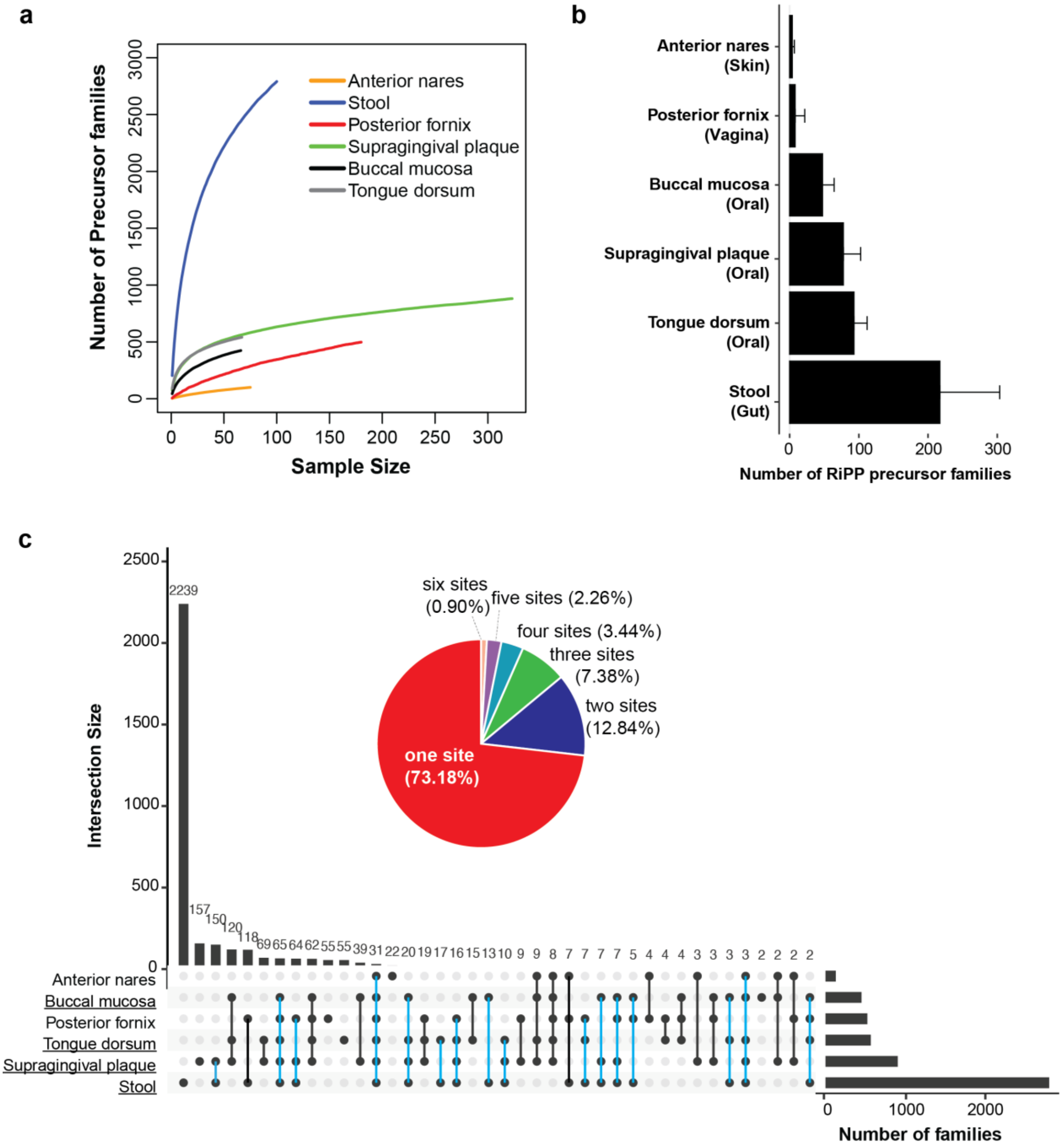

#### Supplementary Figure 7 Variable prevalence of RiPP precursor families detected across six body sites

**a** Accumulation curves of RiPP precursor families were identified from six body sites. The metagenome samples collected from six body sites are stool, 100; supragingival plaque, 281; posterior fornix, 169; tongue dorsum, 66; buccal mucosa, 66; anterior nares, 66. **b** Bar plot showing the number of RiPP families detected in the oral cavity, skin, vagina, and gut samples. The data are presented as mean ± standard deviation. **c** Intersections of RiPP families were detected across six body sites. Inner: The pie chart illustrates the distribution of RiPP families across different body sites, represented as a percentage. Outside: The diagram displays the intersection of RiPP precursor families detected in each body site. The right bar plot provides the total count of RiPP precursor families for each corresponding body site. The top bar plot represents the number of RiPP precursor families in each intersection. Connecting lines are drawn to indicate intersections present in multiple sites. The RiPP families detected in both the oral and gut sites are highlighted in blue lines. The presence of single dots suggests niche-specific RiPP precursor families.

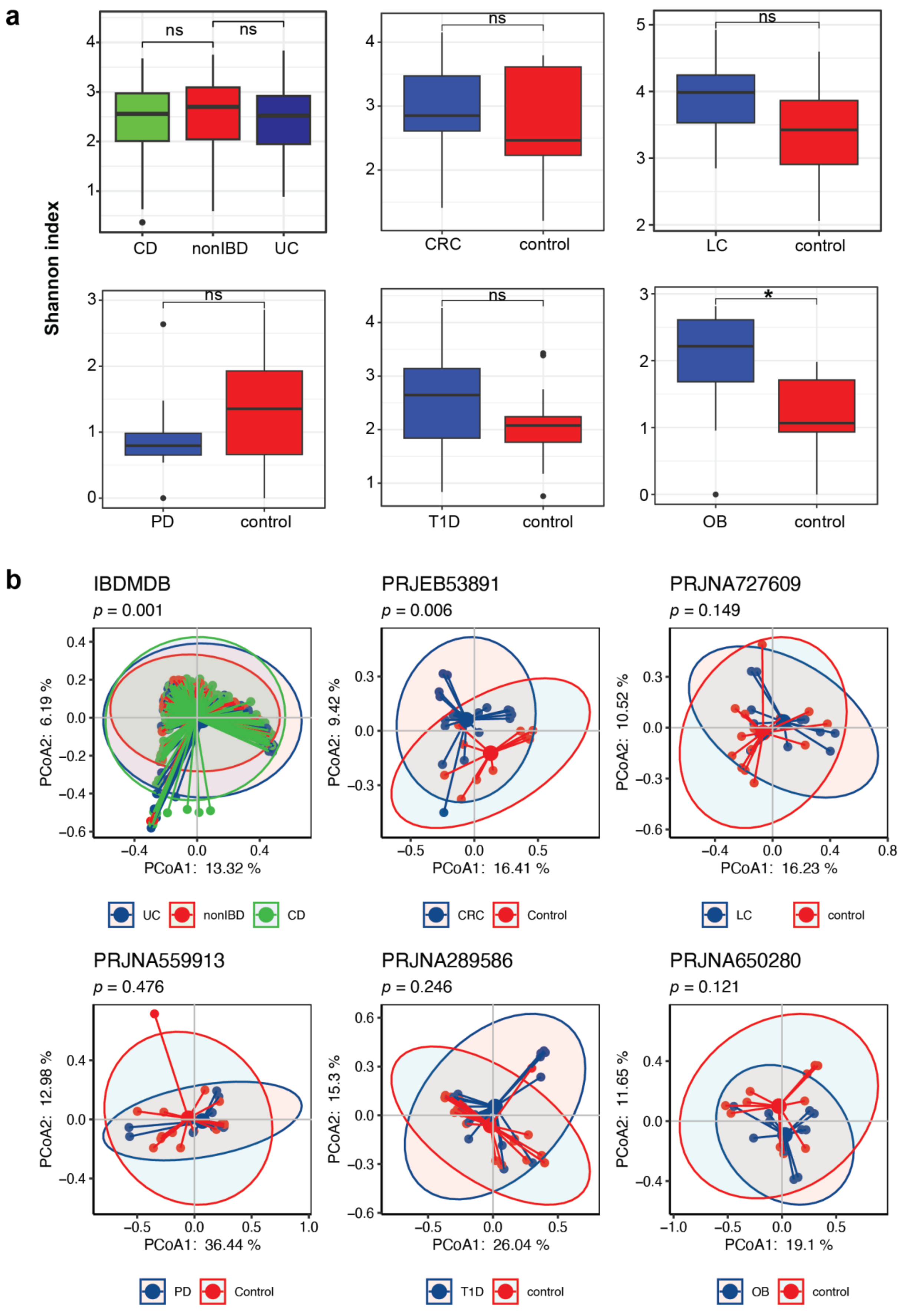

**Supplementary Figure 8 RiPP diversity in healthy individuals and patients with diseases**

**a** Alpha diversity measures are presented using the Shannon index. Box plots a depict the median, lower, and upper quartiles. The whiskers represent the minimum and maximum spread of the data. Dots represent the outliers. Asterisks indicate significance from the Wilcoxon rank sum test (* $p < 0.05$, *** $p < 0.001$). **b** The dissimilarity of samples for beta diversity was assessed using the Bray-Curtis distance, demonstrating the abundance differences of RiPP precursor families detected in metatranscriptomic samples across patients and healthy individuals in disease cohorts. Ellipses in the plot represent a 95% confidence level. P values indicate significance from the significance of PERMANOVA.

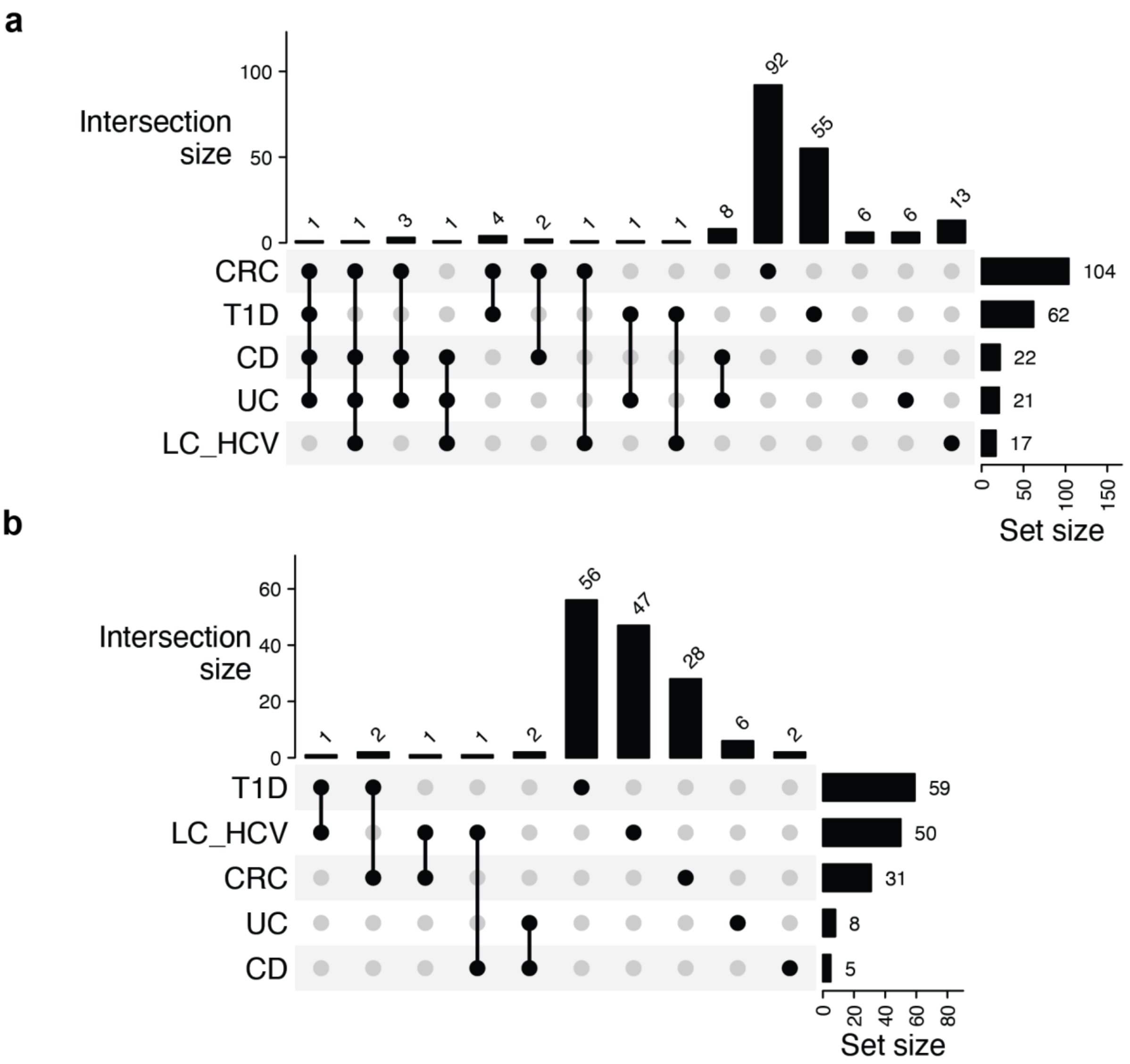

**Supplementary Figure 9 Intersection of differentially abundant RiPP precursor families in metatranscriptome data**

**a** The intersection of the number of transcribed RiPP precursor families significantly enriched in healthy groups. **b** The intersection of transcribed RiPP precursor families significantly enriched disease groups. The bar plot on the right displays the number of differentially abundant RiPP clusters related to the corresponding disease. The bar plot on the top refers to the number of differentially abundant RiPP precursor families identified in each disease intersection. Connecting lines are drawn if an intersection exists in more than one cohort.

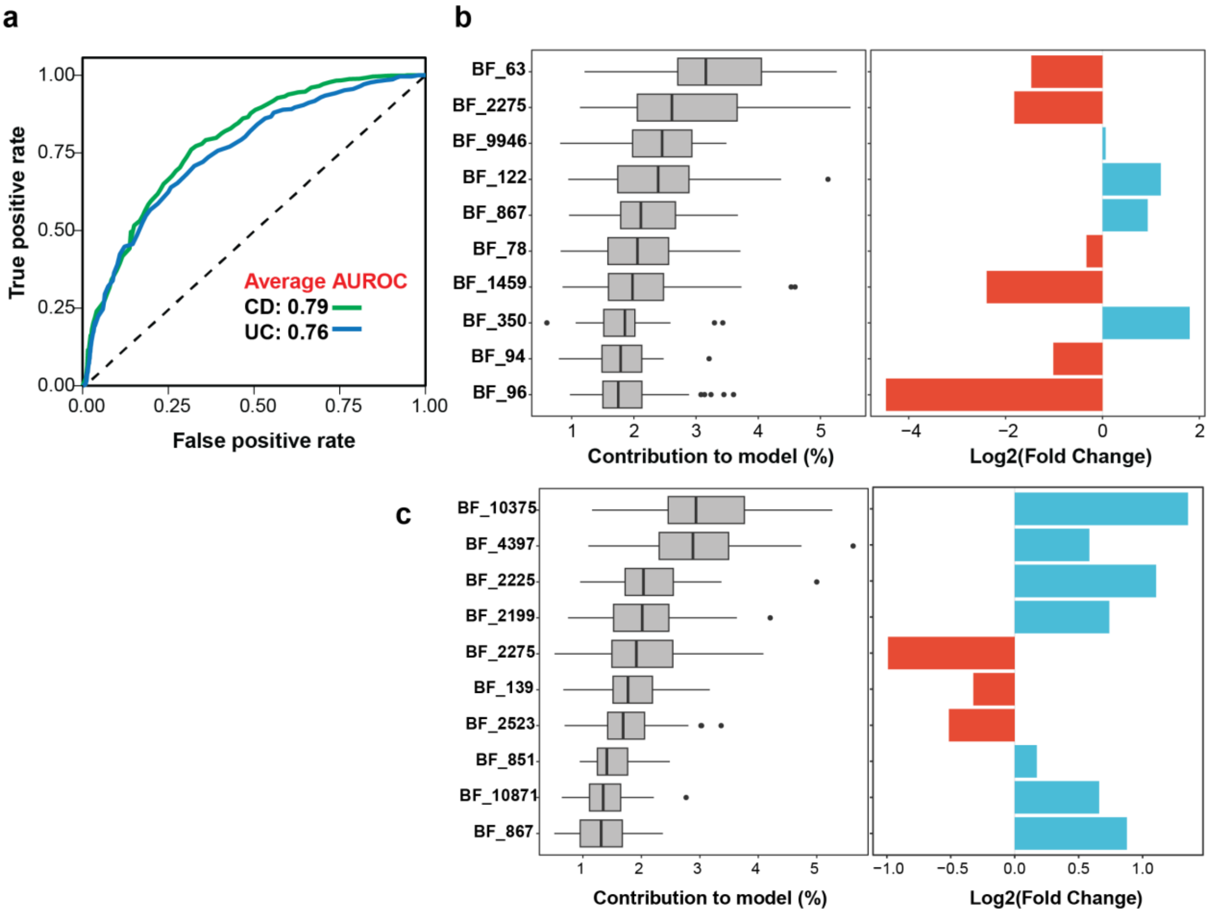

#### Supplementary Figure 10 Biomarker families in IBD disease

**a** The performance of the random forest classifiers based on RiPP abundance in discriminating the health group from the disease group. (**b & c**) The box plot on the left illustrates the top 10 families with the greatest contribution to the discriminative model for CD **b** and UC **c**. The corresponding log2 fold change is depicted in the bar plot on the right side. The red bar represents precursor families enriched in the health group, while the blue bar represents RiPP families enriched in the CD group **b** or UC group **c**.

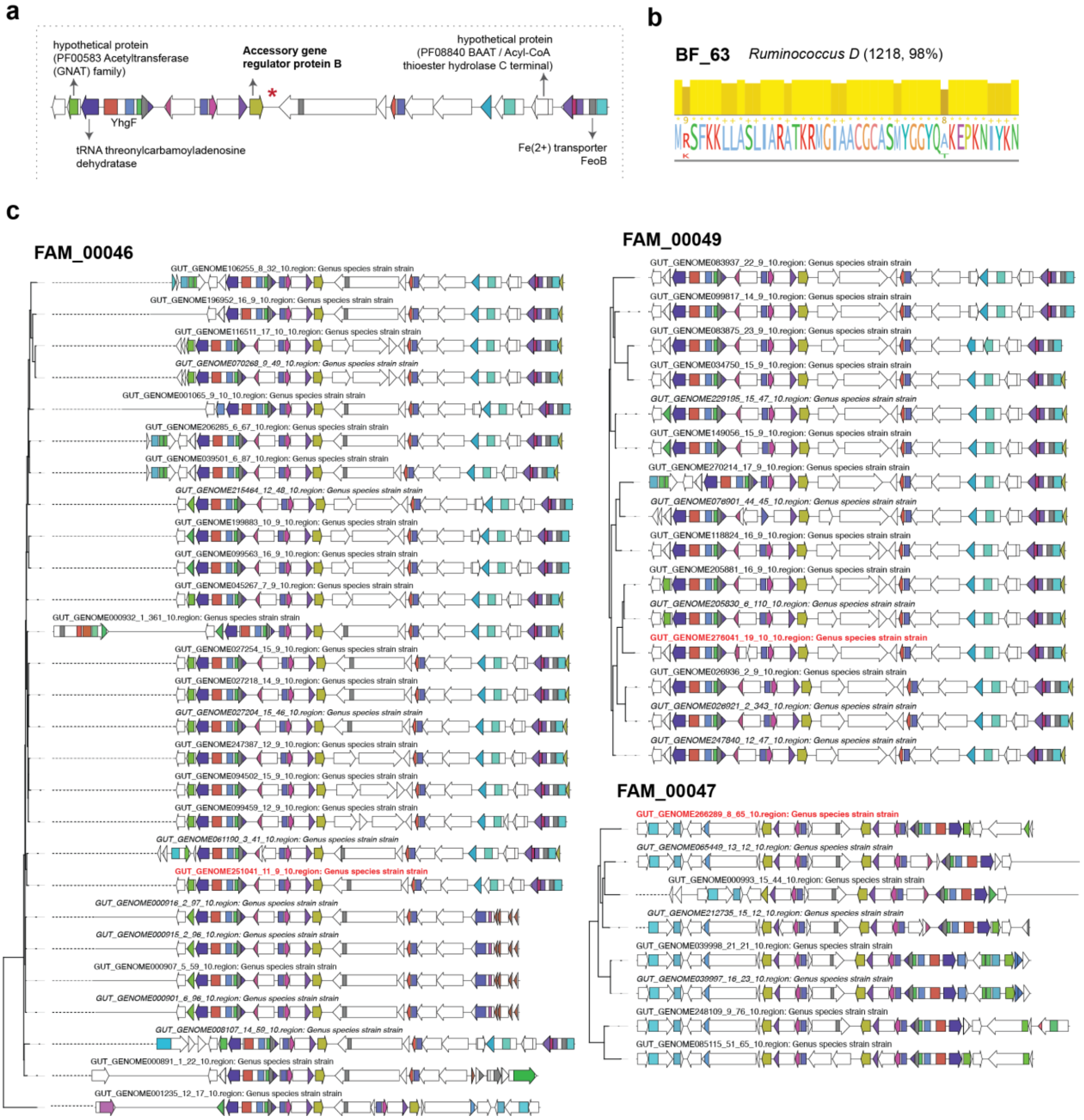

#### Supplementary Figure 11 Representative genomic contexts of RiPP family BF_63

**a** Representative of biosynthetic gene cluster was chosen to show the conserved domain and protein products annotated by BiG-SCAPE in the genomic neighborhood. The location of the precursor was noted with a red asterisk. **b** The sequence log shows a multiple-sequence alignment of family members' shared amino acid residues (sequence logo) and their conservation (bar plot). **c** The top 50 formative genomic contexts (larger gene sizes) of members in the family were chosen for analysis, which were further annotated and grouped into different clades by BiG-SCAPE. The members within this family are predominately encoded by Ruminococcus D (1218, 98%).

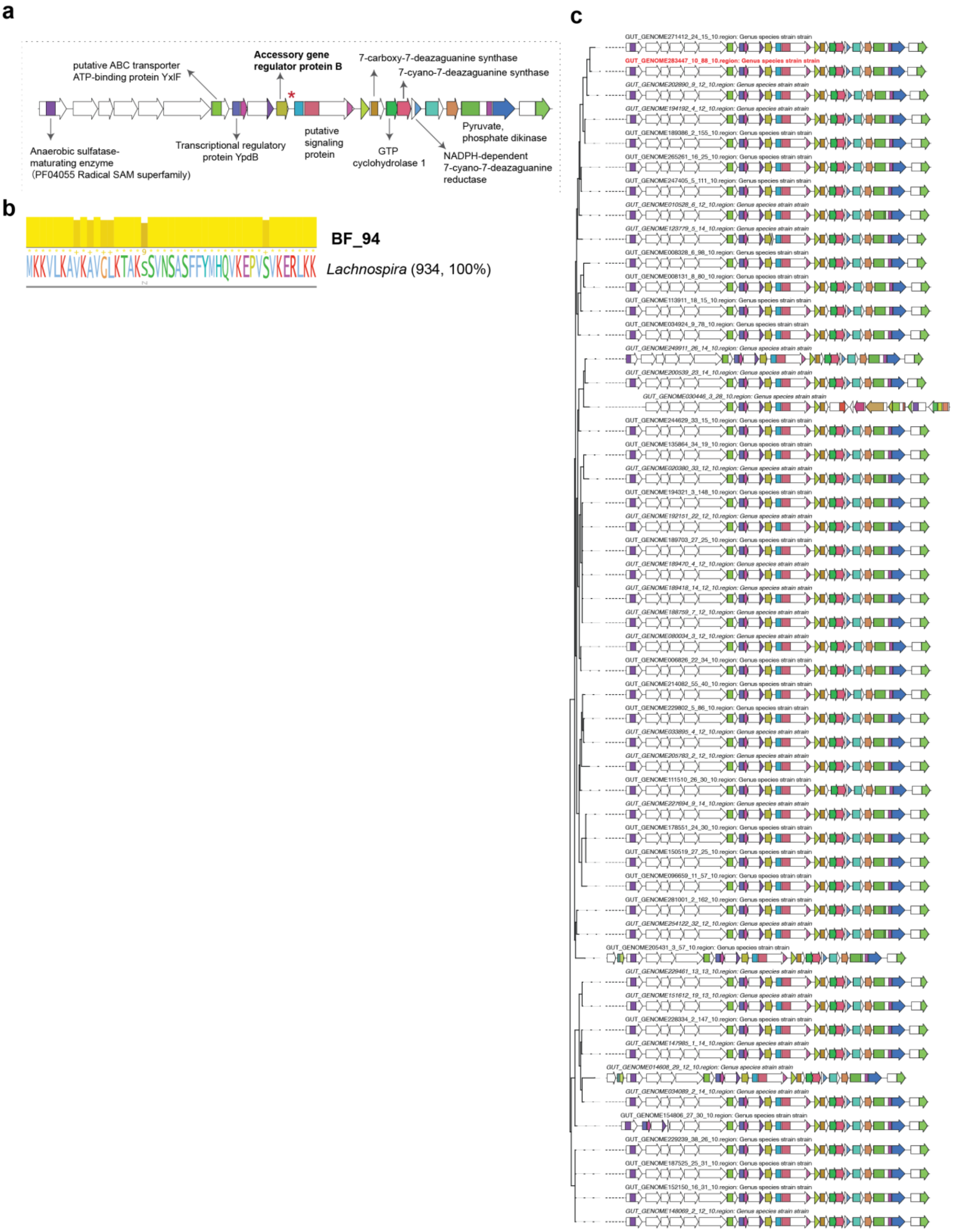

#### Supplementary Figure 12 Representative genomic contexts of RiPP family BF_94

**a** A representative biosynthetic gene cluster was chosen to show the conserved domain and protein products annotated by BiG-SCAPE in the genomic neighborhood. The location of the precursor was noted with a red asterisk. **b** The sequence log shows a multiple-sequence alignment of family members' shared amino acid residues (sequence logo) and their conservation (bar plot). **c** The top 50 formative genomic contexts (larger gene sizes) of members in the family were chosen for analysis, which were further annotated and grouped into different clades by BiG-SCAPE. The members within this family are predominately encoded by *Lachnospira* (934, 100%).

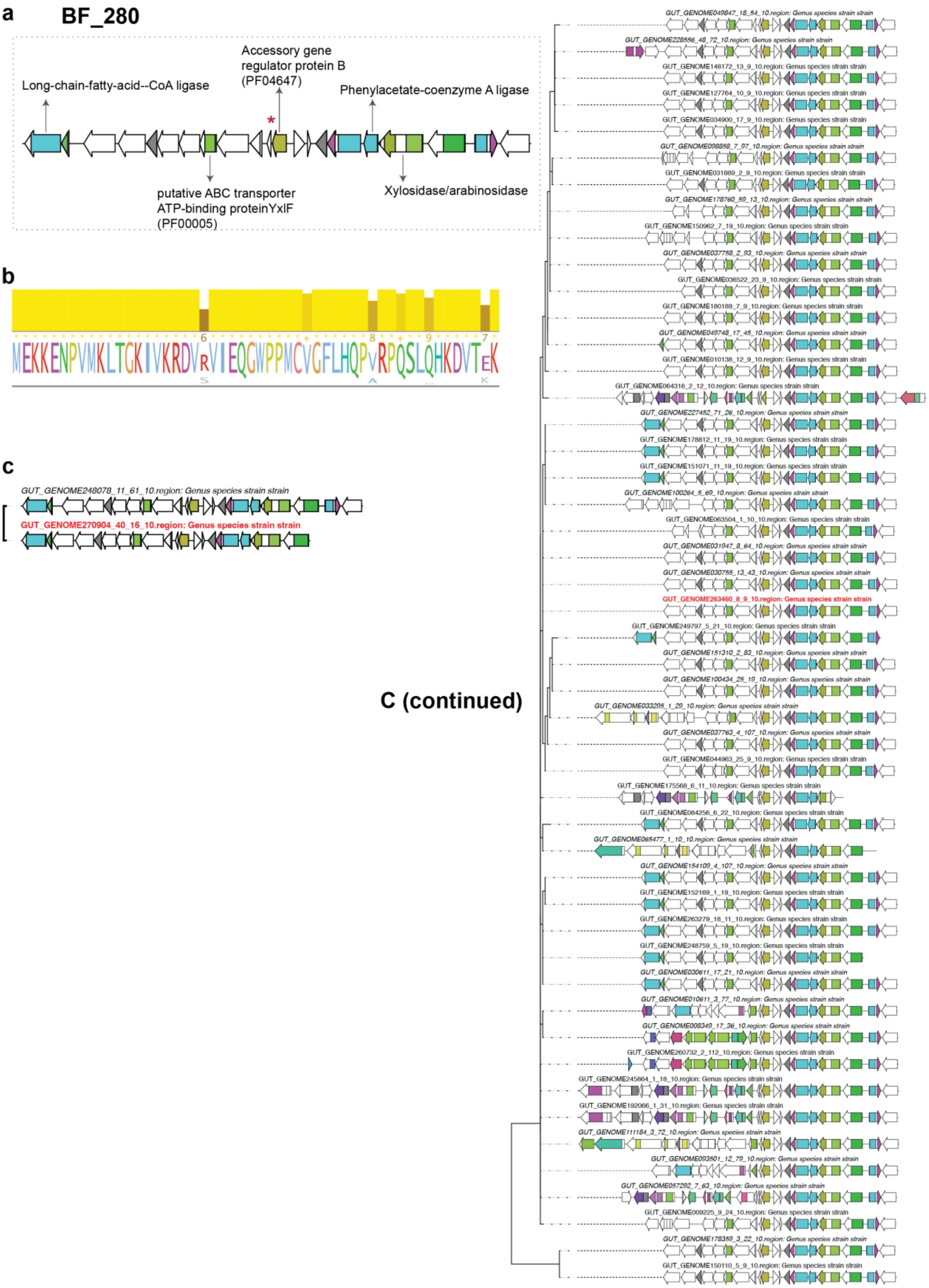

#### Supplementary Figure 13 Representative genomic contexts of RiPP family BF_280

**a** A representative biosynthetic gene cluster was chosen to show the conserved domain and protein products annotated by BiG-SCAPE in the genomic neighborhood. The location of the precursor was noted with a red asterisk. **b** The sequence log shows a multiple-sequence alignment of family members' shared amino acid residues (sequence logo) and their conservation (bar plot). **c** The top 50 formative genomic contexts (larger gene sizes) of members in the family were chosen for analysis, which were further annotated and grouped into different clades by BiG-SCAPE.

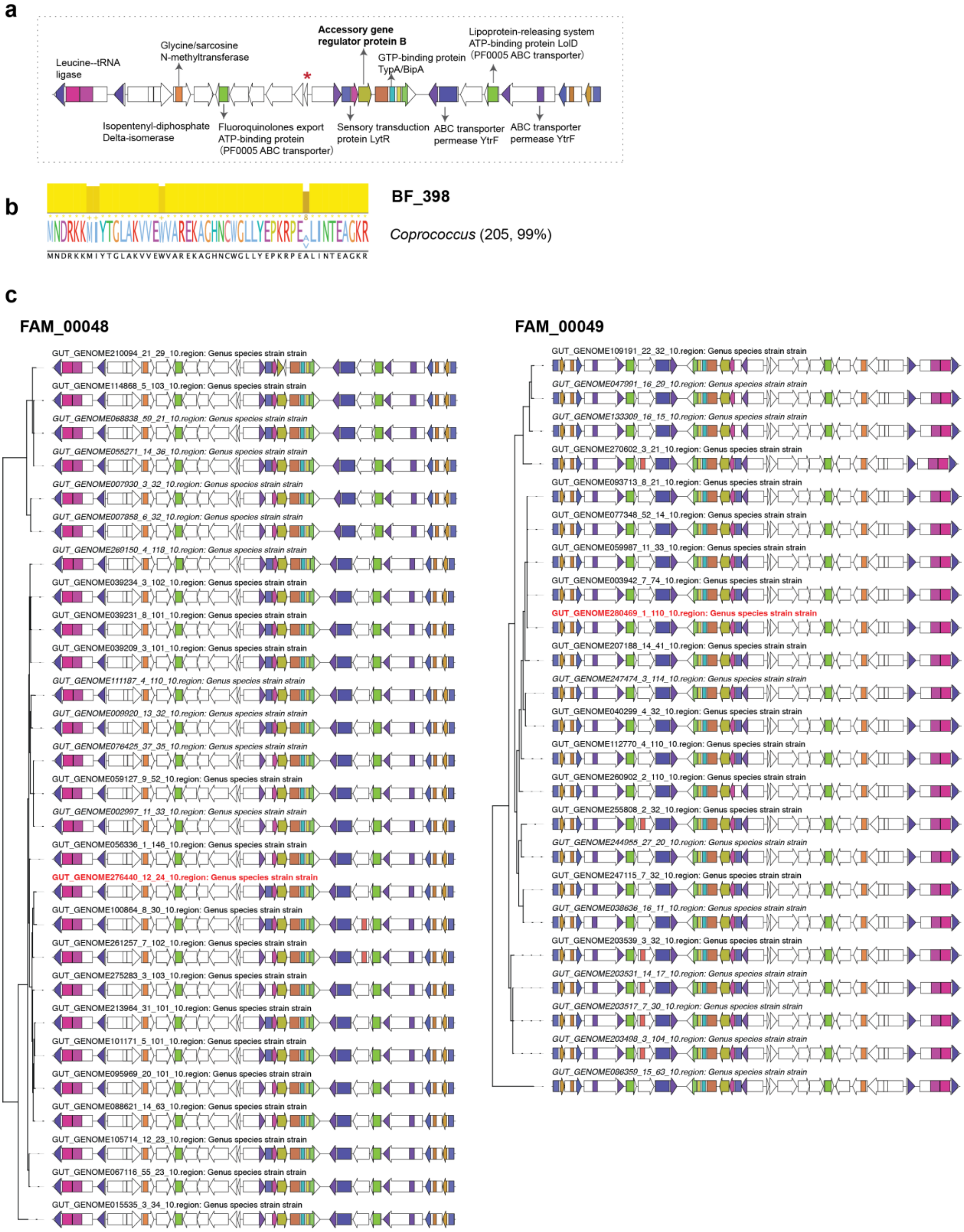

**Supplementary Figure 14 Representative genomic contexts of RiPP family BF_398**

**a** A representative biosynthetic gene cluster was chosen to show the conserved domain and protein products annotated by BiG-SCAPE in the genomic neighborhood. The location of the precursor was noted with a red asterisk. **b** The sequence log shows a multiple-sequence alignment of family members' shared amino acid residues (sequence logo) and their conservation (bar plot). **c** The top 50 formative genomic contexts (larger gene sizes) of members in the family were chosen for analysis, which were further annotated and grouped into different clades by BiG-SCAPE. The members within this family are predominately encoded by *Coprococcus* (205, 99%).

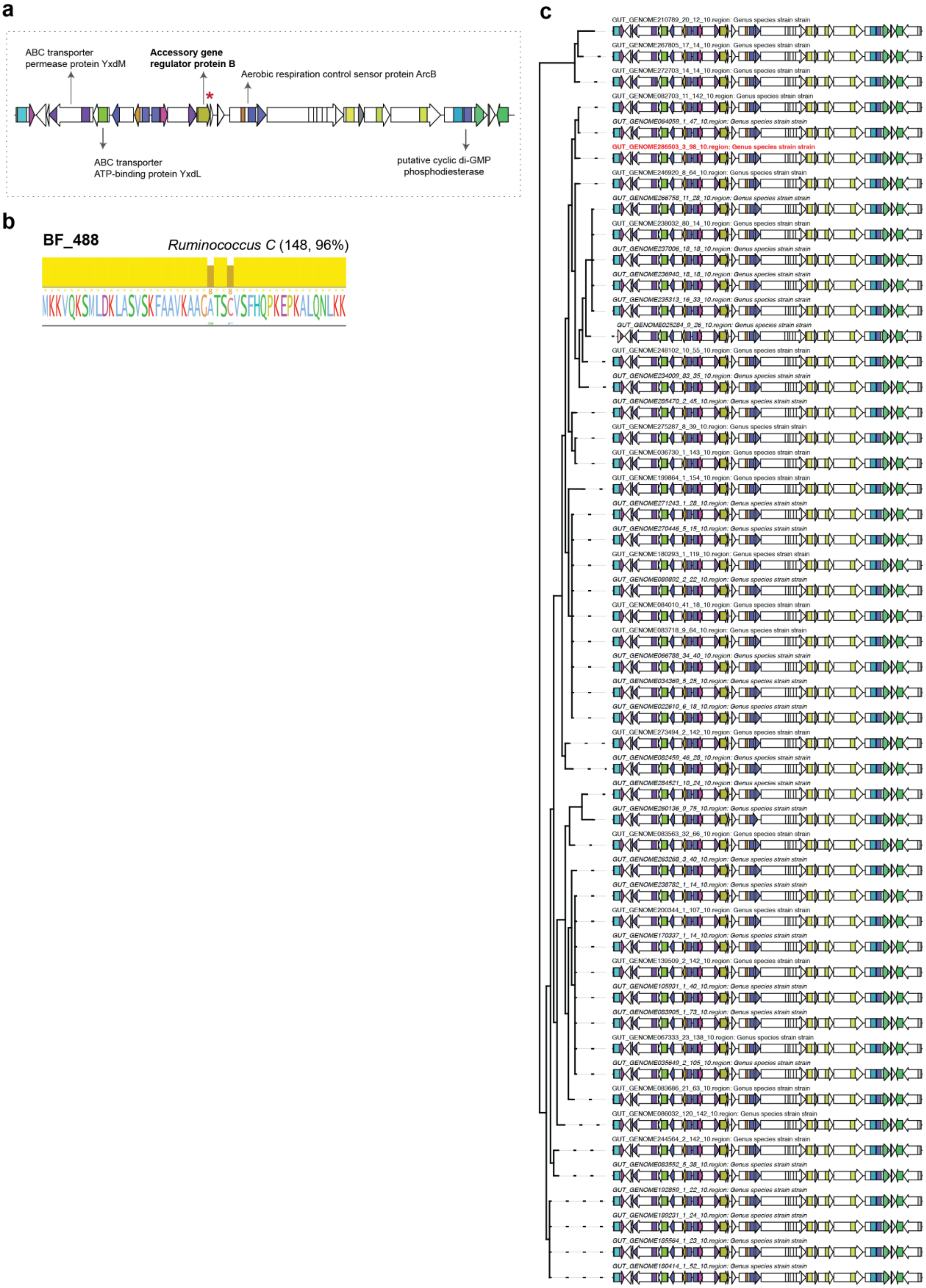

#### Supplementary Figure 15 Representative genomic contexts of RiPP family BF_488

**a** A representative biosynthetic gene cluster was chosen to show the conserved domain and protein products annotated by BiG-SCAPE in the genomic neighborhood. The location of the precursor was noted with a red asterisk. **b** The sequence log shows a multiple-sequence alignment of family members' shared amino acid residues (sequence logo) and their conservation (bar plot). **c** The top 50 formative genomic contexts (larger gene sizes) of members in the family were chosen for analysis, which were further annotated and grouped into different clades by BiG-SCAPE. The members within this family are predominately encoded by *Ruminococcus* C (148, 96%).

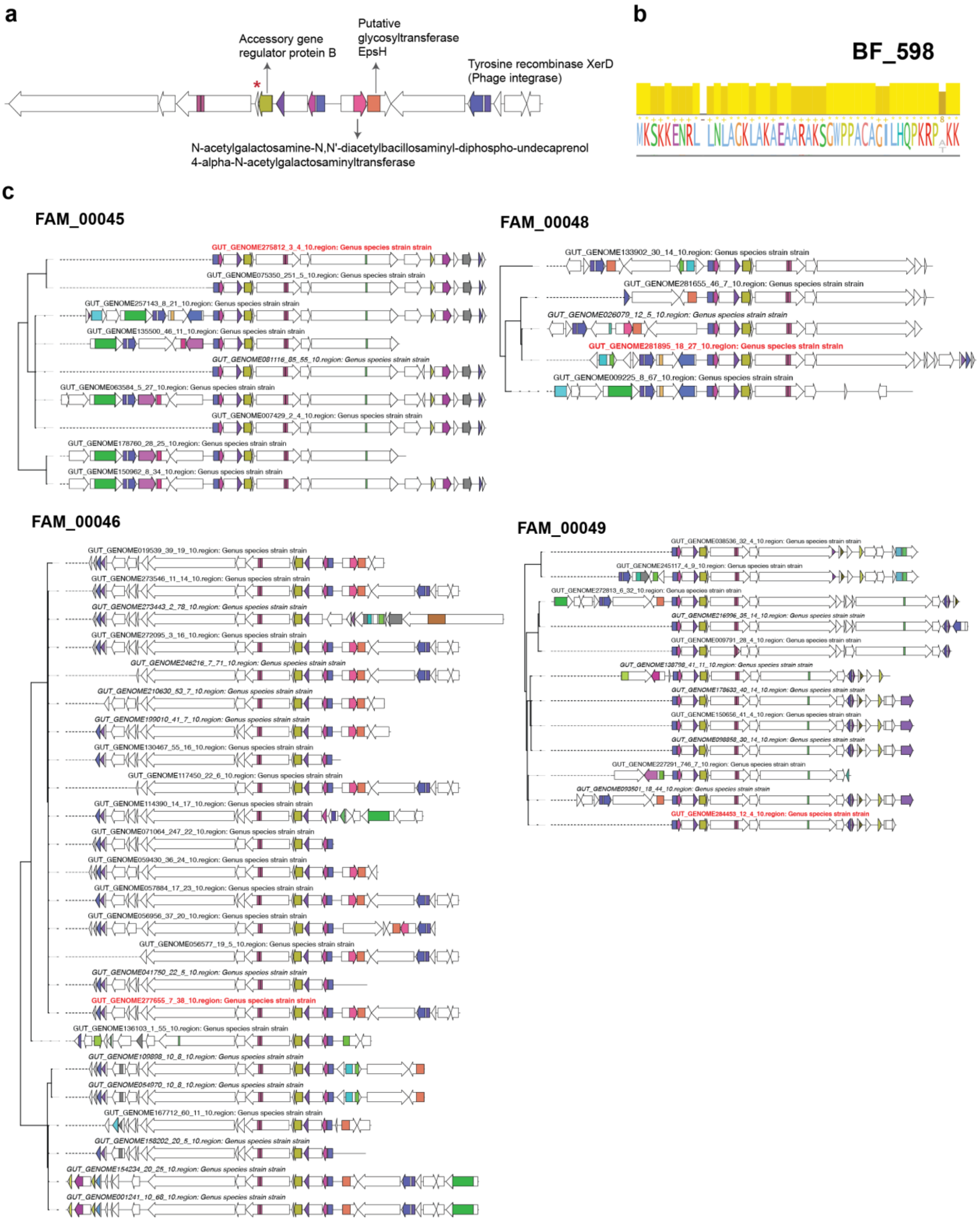

#### Supplementary Figure 16 Representative genomic contexts of RiPP family BF_598

**a** A representative biosynthetic gene cluster was chosen to show the conserved domain and protein products annotated by BiG-SCAPE in the genomic neighborhood. The location of the precursor was noted with a red asterisk. **b** The sequence log shows a multiple-sequence alignment of family members' shared amino acid residues (sequence logo) and their conservation (bar plot). **c** The top 50 formative genomic contexts (larger gene sizes) of members in the family were chosen for analysis, which were further annotated and grouped into different clades by BiG-SCAPE.

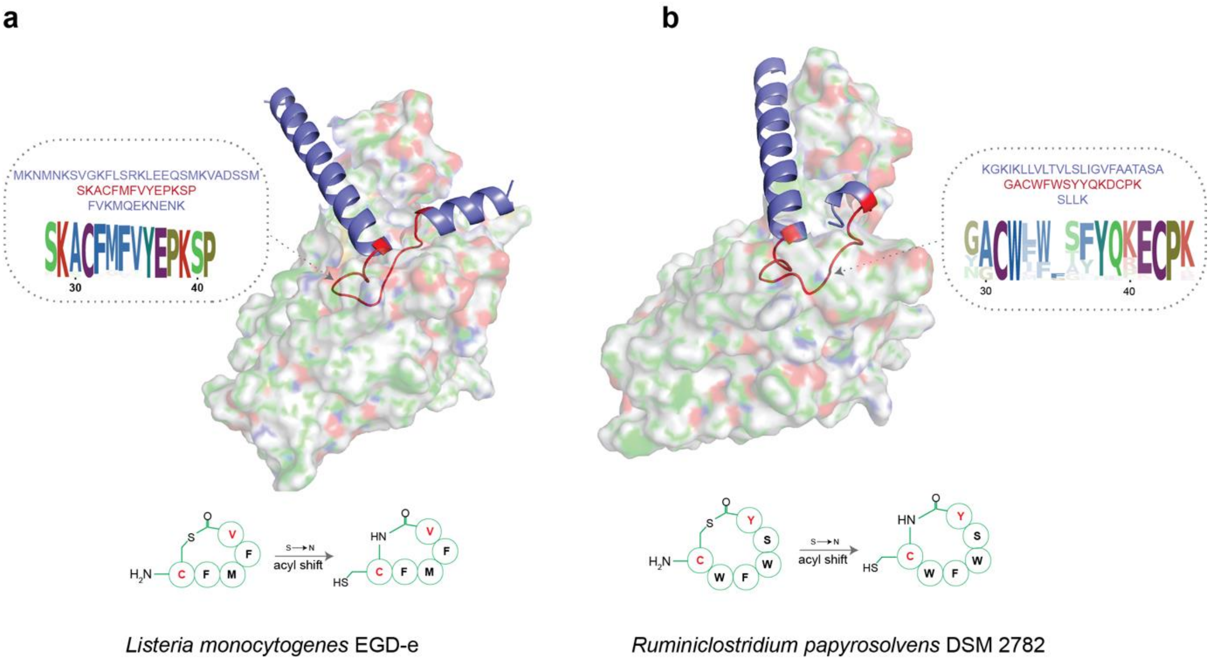

#### Supplementary Figure 17 Reported AIPs from *Listeria monocytogenes* EGD-e and *Ruminiclostridium papyrosolvens* DSM 2782

The upper panel depicts the predicted interaction between the precursor peptide and AgrB enzyme, represented as a complex. The circled region indicates the precursor sequence, while the potentially modified core region is highlighted in red. Sequence logo analysis demonstrates a high level of conservation within this core region across the precursor family. The bottom panel displays the predicted structure. This structure corresponds to experimentally verified mature AIP derived from *Listeria monocytogenes* EGD-e[14-16] **a** and *Ruminiclostridium papyrosolvens* DSM 2782[17] **b.**

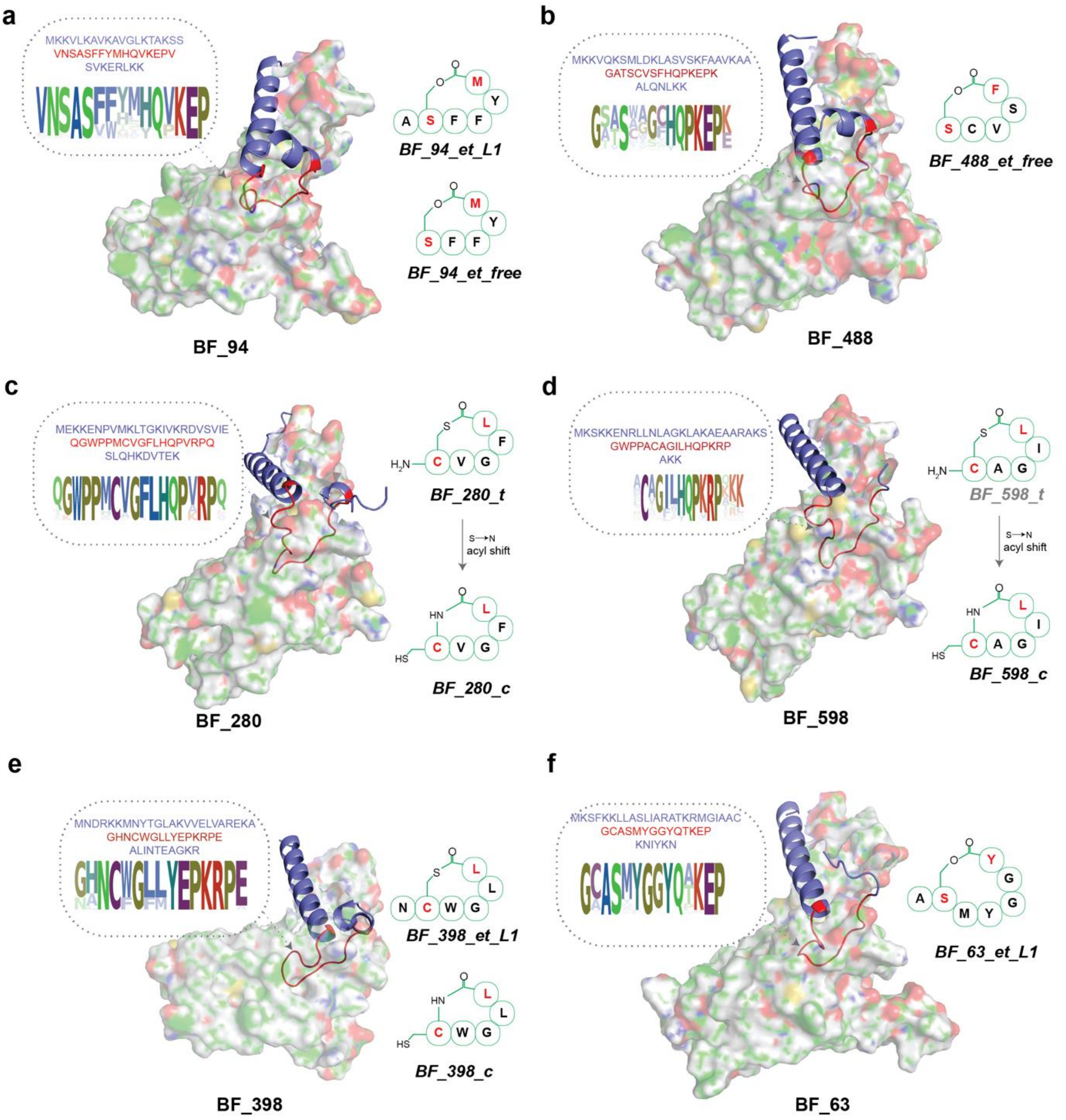

#### Supplementary Figure 18 Putative mature AIPs in this study

The depicted complex represents the predicted interaction between the precursor peptide and AgrB enzyme. The circled region within the complex corresponds to the precursor sequence, while the potentially modified core region is highlighted in red. Sequence logo analysis reveals a high level of conservation within this core region across the precursor family, which aids in predicting the exocyclic amino acids. The structures displayed on the right side represent the putative mature AIP products. Each compound is labeled with a specific suffix to denote its characteristics. For example, “et_L1” indicates that the AIP has an exotail with a length of 1 amino acid, “et_free” signifies an AIP without an exotail, “c” denotes cyclopeptides, and “t” represents exotail-free AIPs with a thiolactone group.

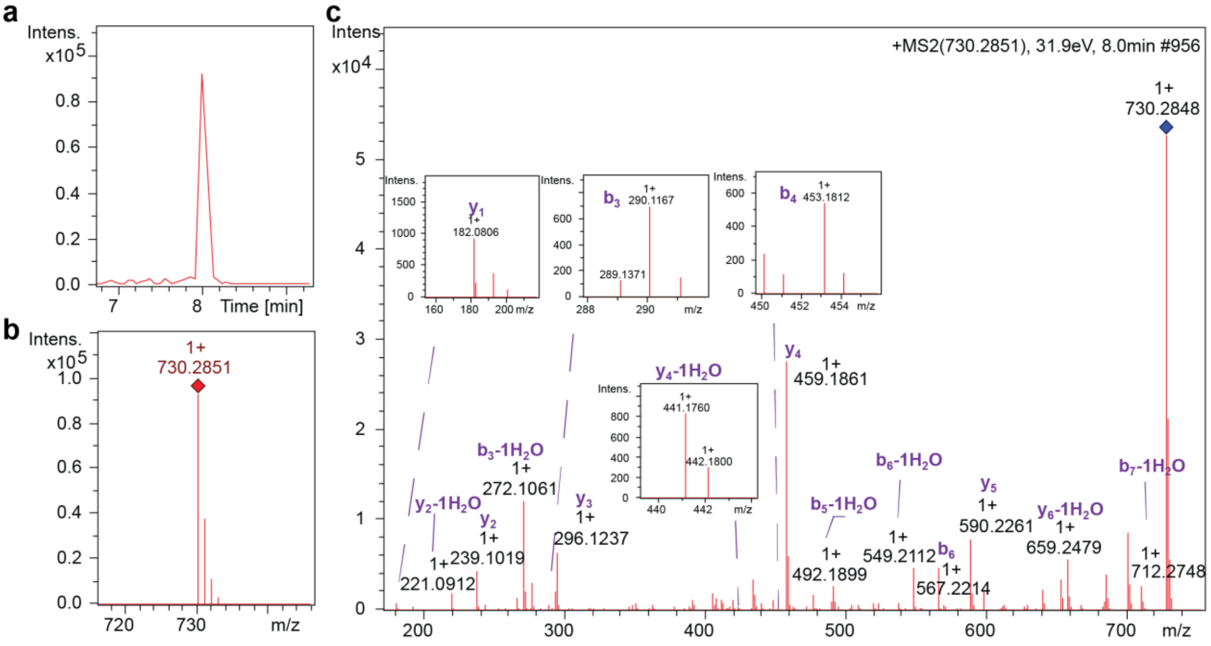

d

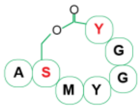

| ions | fragment | Observed m/z | Calculated m/z | charge | ppm |
|---|---|---|---|---|---|
| y1 | Y | 182.0806 | 182.0812 | 1 | 2.97 |
| y2-1$H_2O$ | GY | 221.0912 | 221.0921 | 1 | 3.82 |
| y2 | GY | 239.1019 | 239.1026 | 1 | 2.88 |
| y3 | GGY | 296.1237 | 296.1241 | 1 | 1.37 |
| y4-1$H_2O$ | YGGY | 441.1760 | 441.1769 | 1 | 1.88 |
| y4 | YGGY | 459.1861 | 459.1874 | 1 | 2.90 |
| y5 | MYGGY | 590.2261 | 590.2279 | 1 | 3.06 |
| y6-1$H_2O$ | SMYGGY | 659.2479 | 659.2494 | 1 | 2.25 |
| b3-1$H_2O$ | ASM | 272.1061 | 272.1063 | 1 | 0.87 |
| b3 | ASM | 290.1167 | 290.1169 | 1 | 0.61 |
| b4 | ASMY | 453.1812 | 453.1802 | 1 | 2.15 |
| b5-1$H_2O$ | ASMYG | 492.1899 | 492.1911 | 1 | 2.51 |
| b6-1$H_2O$ | ASMYGG | 549.2112 | 549.2126 | 1 | 2.60 |
| b6 | ASMYGG | 567.2214 | 567.2232 | 1 | 3.11 |
| b7-1$H_2O$ | ASMYGGY | 712.2748 | 712.2759 | 1 | 1.57 |

#### Supplementary Figure 19 UPLC-HRMS/MS analysis of synthesized BF_63_et_L1

The retention time **a**, MS1 spectra **b**, and HRMS2-based fragmentations **c** of synthetic peptide were shown. **d** Listed are the paired ions and possible fragments. The structure of synthetic AIP is displayed on the left side.

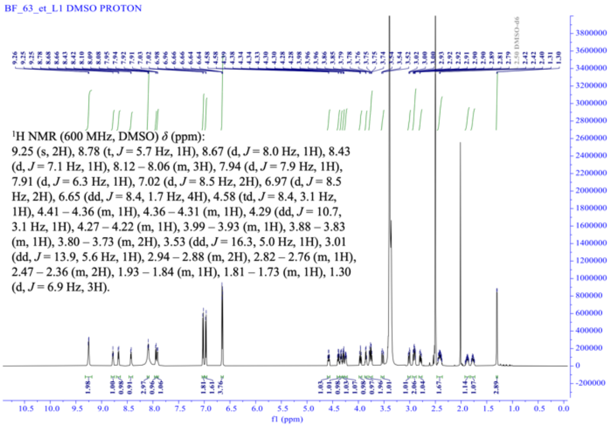

$^{1}$H NMR spectrum (600 MHz, DMSO-$d_6$) of BF_63_et_L1.

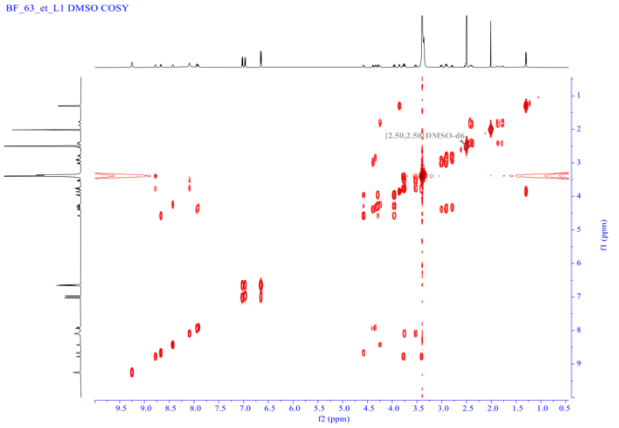

COSY NMR spectrum (600 MHz, DMSO-$d_6$) of BF_63_et_L1.

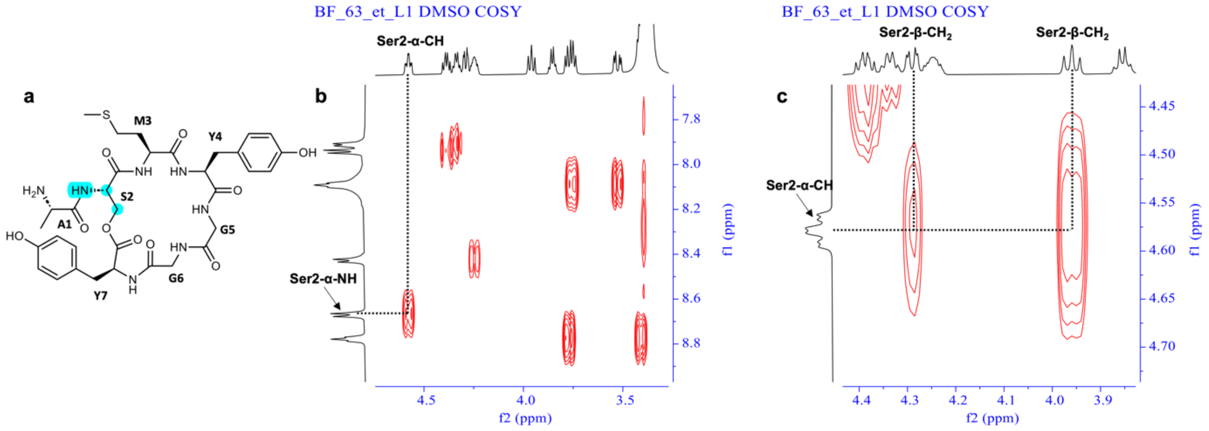

#### Supplementary Figure 20 NMR assignment of BF_63_et_L1

Key COSY correlations for the proton in the serine residue of BF_63_et_L1. **a** The chemical structure of BF_63_et_L1, and the groups in the blue background corresponds to the proton in the key serine residue. **b** Key COSY correlations for Ser2-α-NH/Ser2-α-CH. **c** Key COSY correlations for Ser2-α-CH/Ser2-β-$CH_2$.

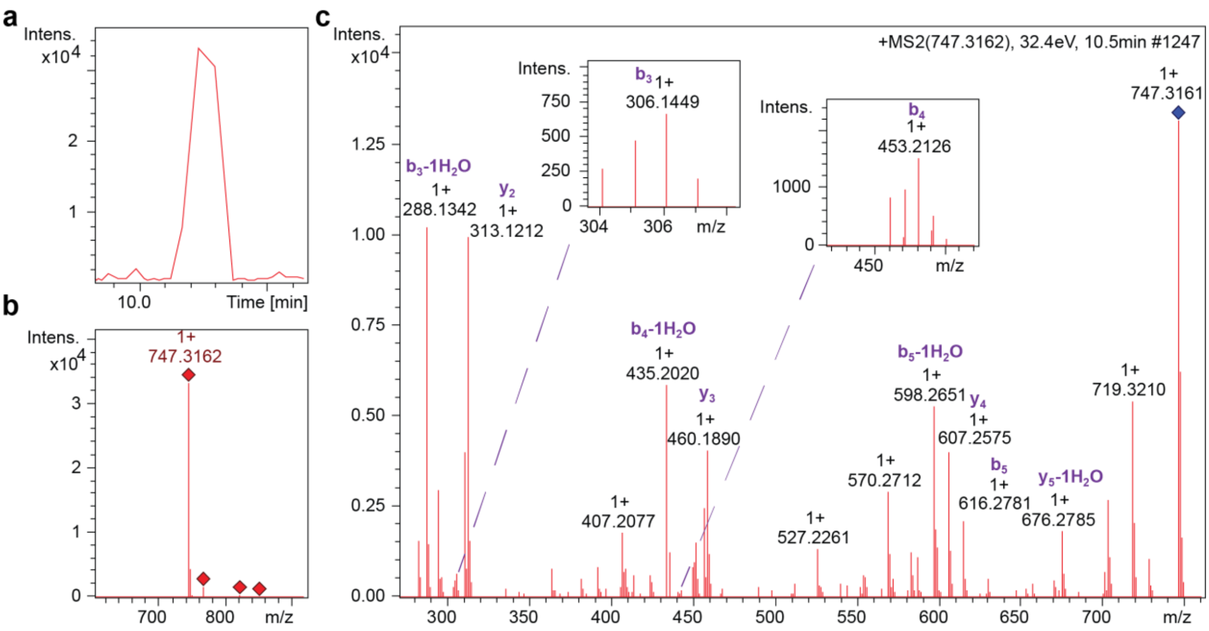

d

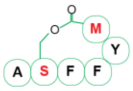

| ions | fragment | Observed m/z | Calculated m/z | charge | ppm |
|---|---|---|---|---|---|
| $y_2$ | YM | 313.1212 | 313.1217 | 1 | -1.45 |
| $y_3$ | FYM | 460.1890 | 460.1901 | 1 | -2.32 |
| $y_4$ | FFYM | 607.2575 | 607.2585 | 1 | -1.62 |
| $y_5$-1$H_2O$ | SFFYM | 676.2785 | 676.2799 | 1 | -2.14 |
| $b_3$-1$H_2O$ | ASF | 288.1342 | 288.1343 | 1 | -0.24 |
| $b_3$ | ASF | 306.1449 | 306.1448 | 1 | 0.22 |
| $b_4$-1$H_2O$ | ASFF | 435.2020 | 435.2027 | 1 | -1.57 |
| $b_4$ | ASFF | 453.2126 | 453.2132 | 1 | -1.41 |
| $b_5$-1$H_2O$ | ASFFY | 598.2651 | 598.2660 | 1 | -1.52 |
| $b_5$ | ASFFY | 616.2781 | 616.2766 | 1 | 2.47 |

**Supplementary Figure 21 UPLC-HRMS/MS analysis of synthesized BF_94_et_L1**

The retention time **a**, MS1 spectra **b**, and HRMS2-based fragmentations **c** of synthetic peptide. **d** Listed are the paired ions and possible fragments. The structure of synthetic AIP is displayed on the left side.

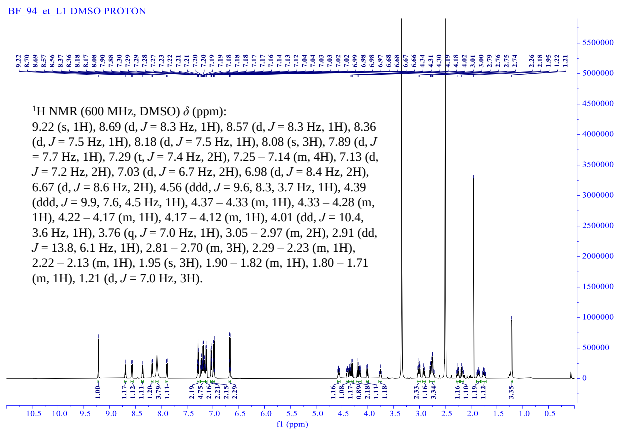

$^{1}$H NMR spectrum (600 MHz, DMSO-$d_6$) of BF_94_et_L1.

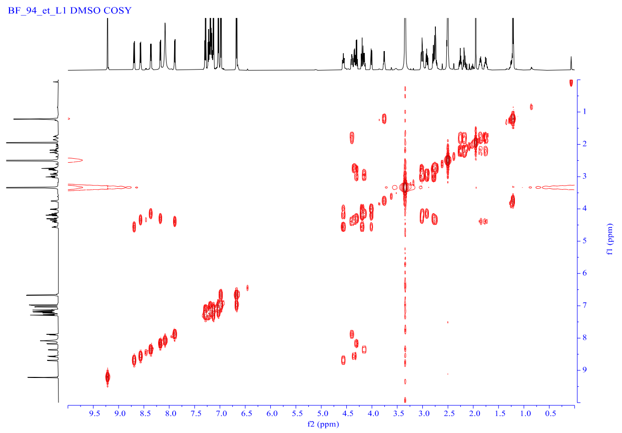

COSY NMR spectrum (600 MHz, DMSO-$d_6$) of BF_94_et_L1.

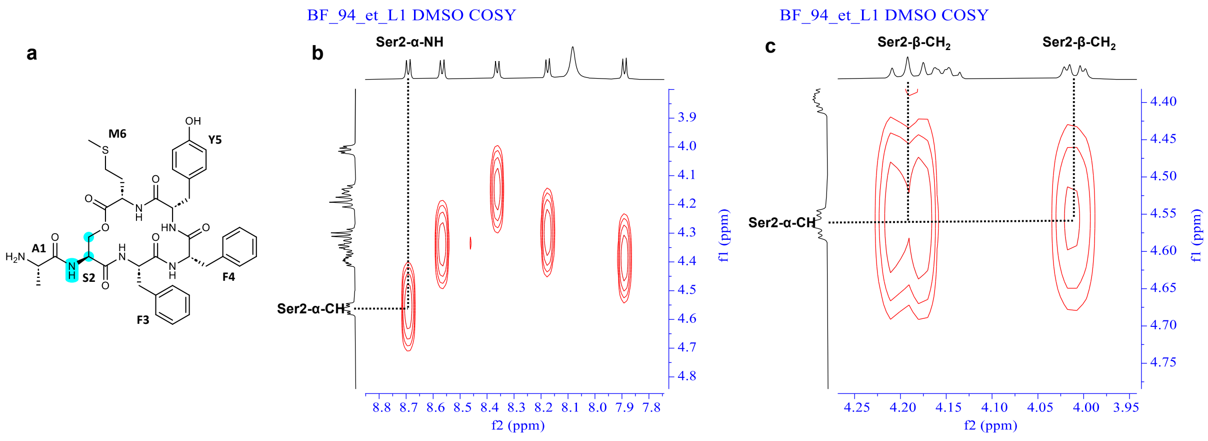

#### Supplementary Figure 22 NMR assignment of BF_94_et_L1

Key COSY correlations for the proton in the serine residue of BF_94_et_L1. **a** The chemical structure of BF_94_et_L1 and the groups in the blue background correspond to the proton in the key serine residue. **b** Key COSY correlations for Ser2-α-NH/ Ser2-α-CH. **c** Key COSY correlations for Ser2-α-CH/Ser2-β-CH2.

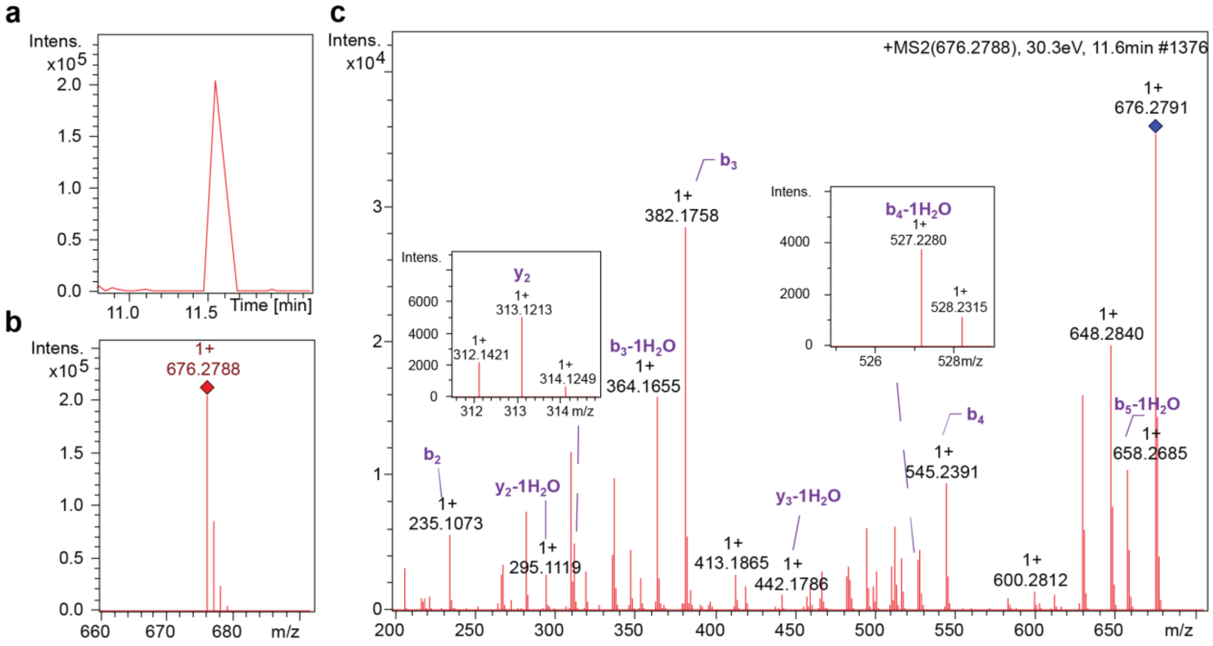

**d**

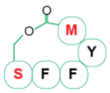

| ions | fragment | Observed m/z | Calculated m/z | charge | ppm |
|---|---|---|---|---|---|
| b2 | SF | 235.1073 | 235.1077 | 1 | 1.68 |
| b3 | SFF | 382.1758 | 382.1761 | 1 | 0.97 |
| b3-1$H_2O$ | SFF | 364.1655 | 364.1656 | 1 | 0.32 |
| b4 | SFFY | 545.2391 | 545.2395 | 1 | 0.71 |
| b4-1$H_2O$ | SFFY | 527.2280 | 527.2289 | 1 | 1.76 |
| b5-1$H_2O$ | SFFYM | 658.2685 | 658.2694 | 1 | 1.28 |
| y2 | YM | 313.1213 | 313.1217 | 1 | 0.99 |
| y2-1$H_2O$ | YM | 295.1119 | 295.1111 | 1 | 2.68 |
| y3-1$H_2O$ | FYM | 442.1786 | 442.1795 | 1 | 2.00 |

#### Supplementary Figure 23 UPLC-HRMS/MS analysis of synthesized BF_94_et_free

The retention time **a,** MS1 spectra **b**, and HRMS2-based fragmentations c of synthetic peptide. **d** Listed are the paired ions and possible fragments. The structure of synthetic AIP is displayed on the left side.

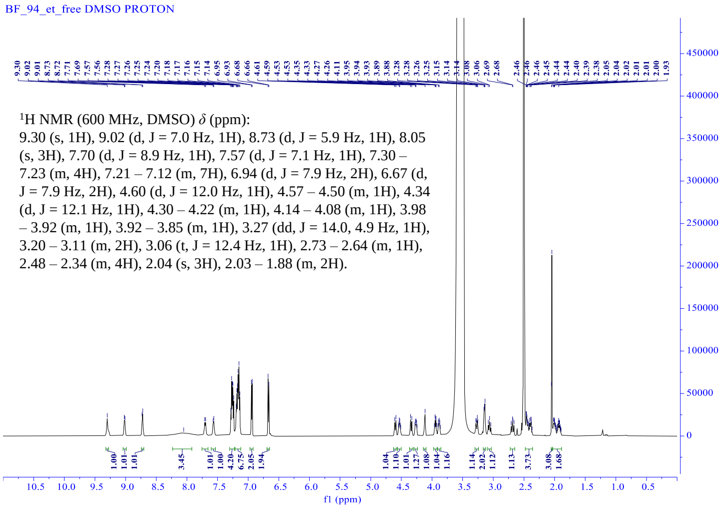

$^1$H NMR spectrum (600 MHz, DMSO-$d_6$) of BF_94_et_free.

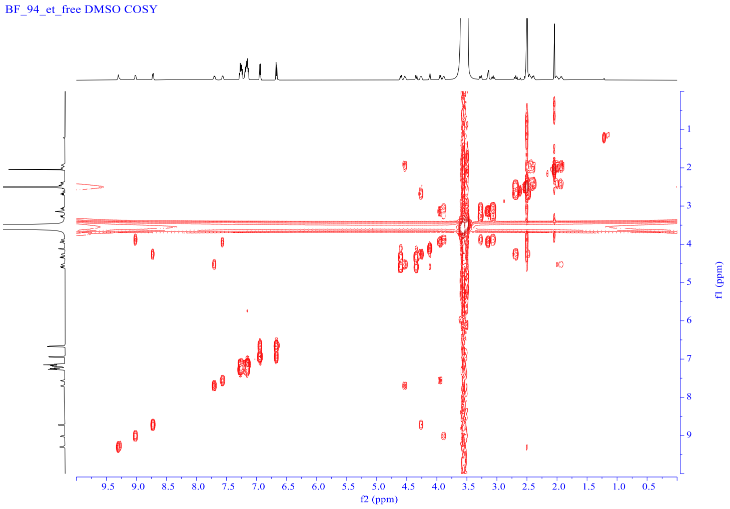

COSY NMR spectrum (600 MHz, DMSO-$d_6$) of BF_94_et_free.

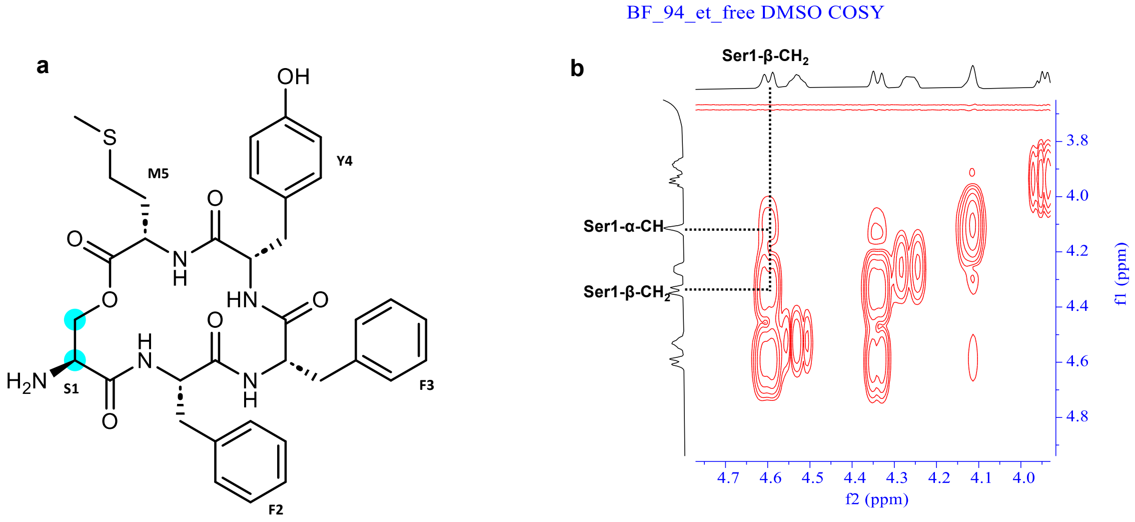

#### Supplementary Figure 24 NMR assignment of BF_94_et_free

Key COSY correlations for the proton in the serine residue of BF_94_et_free. **a** The chemical structure of BF_94_et_free and the groups in the blue background correspond to the proton in the key serine residue. **b** Key COSY correlations for Ser1-α-CH/Ser1-β-$CH_2$.

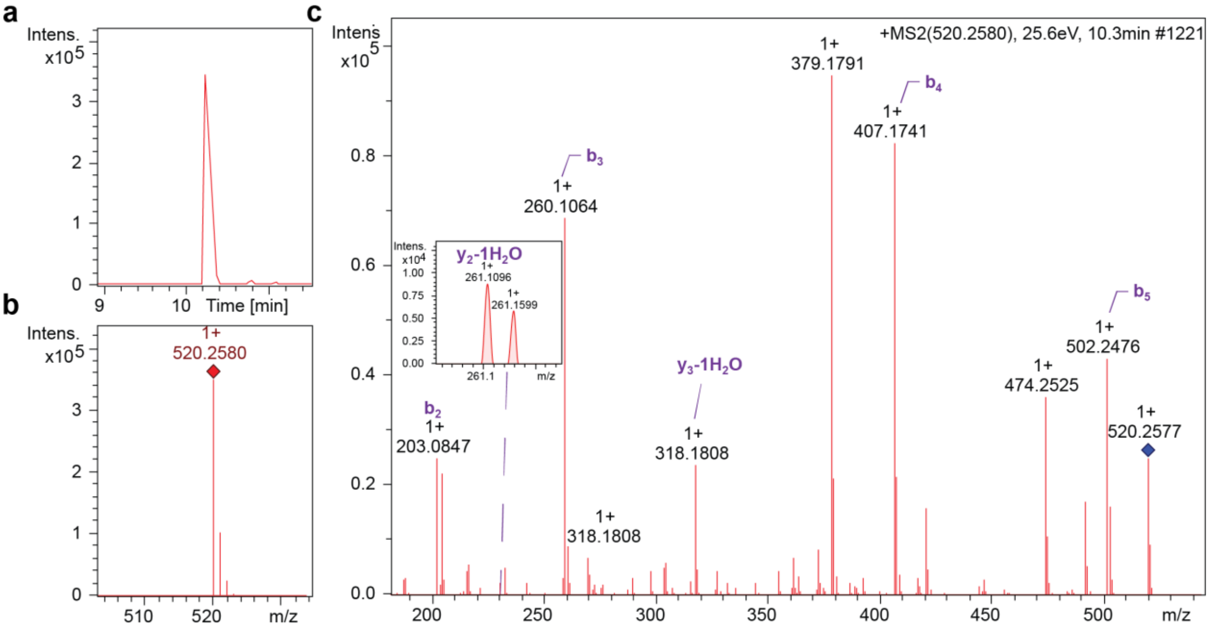

d

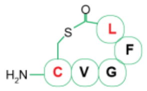

| ions | fragment | Observed m/z | Calculated m/z | charge | ppm |
|---|---|---|---|---|---|
| b2 | CV | 203.0847 | 203.0849 | 1 | 0.63 |
| b3 | CVG | 260.1064 | 260.1063 | 1 | 0.12 |
| b4 | CVGF | 407.1741 | 407.1748 | 1 | 1.52 |
| b5 | CVGFL | 502.2476 | 502.2483 | 1 | 1.34 |
| y2-1$H_2O$ | FL | 261.1599 | 261.1598 | 1 | 0.48 |
| y3-1$H_2O$ | GFL | 318.1808 | 318.1812 | 1 | 1.38 |

**Supplementary Figure 25 UPLC-HRMS/MS analysis of synthesized BF_280_t**

The retention time **a**, MS1 spectra **b,** and HRMS2-based fragmentations **c** of synthetic peptide. **d** Listed are the paired ions and possible fragments. The structure of synthetic AIP is displayed on the left side.

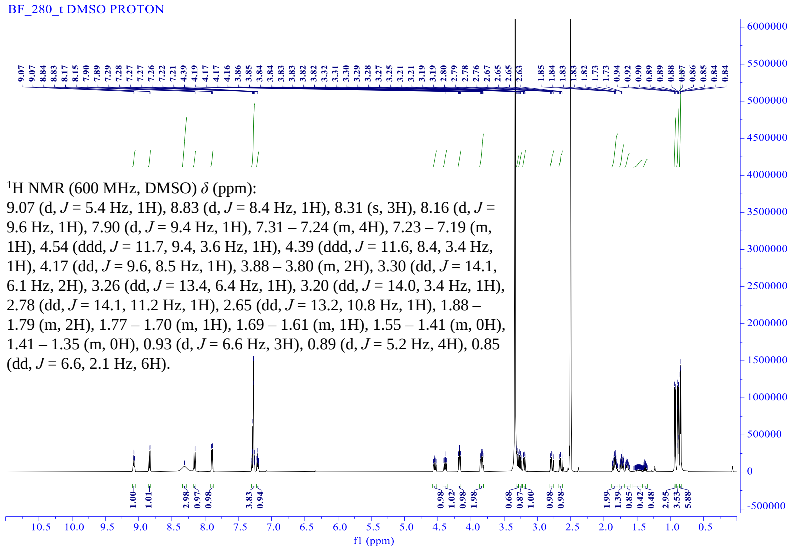

$^1$H NMR spectrum (600 MHz, DMSO-$d_6$) of BF_280_t.

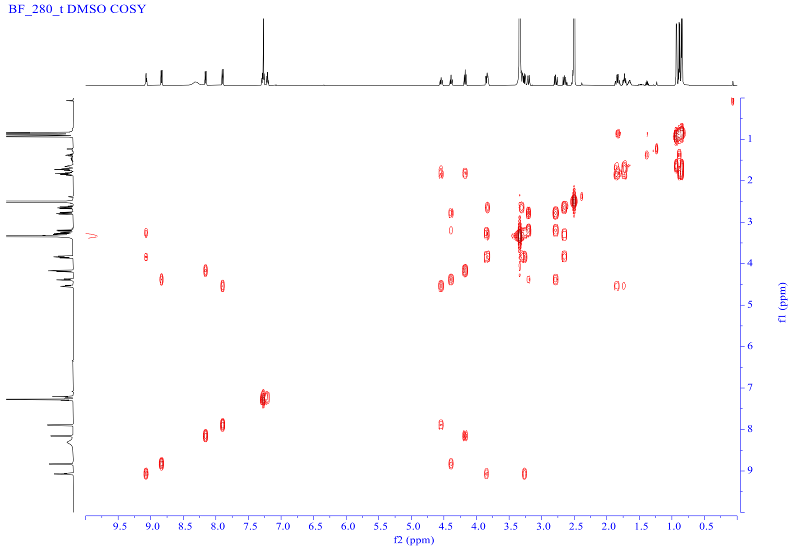

COSY NMR spectrum (600 MHz, DMSO-$d_6$) of BF_280_t.

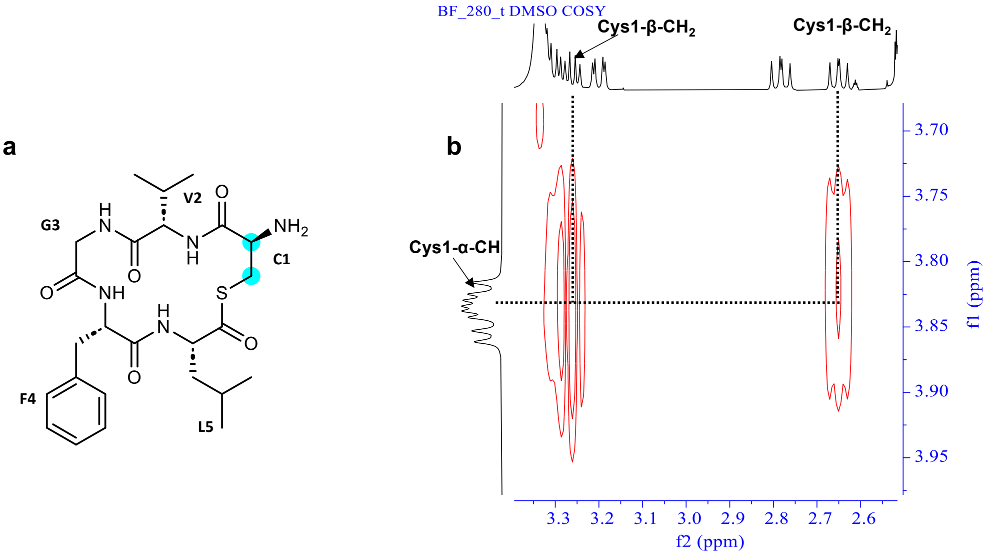

#### Supplementary Figure 26 NMR assignment of BF_280_t

Key COSY correlations for the proton in the cysteine residue of BF_280_t. **a** The chemical structure of BF_280_t and the groups in the blue background correspond to the proton in the key cysteine residue. **b** Key COSY correlations for Cys1-α-CH/Cys1-β-$CH_2$.

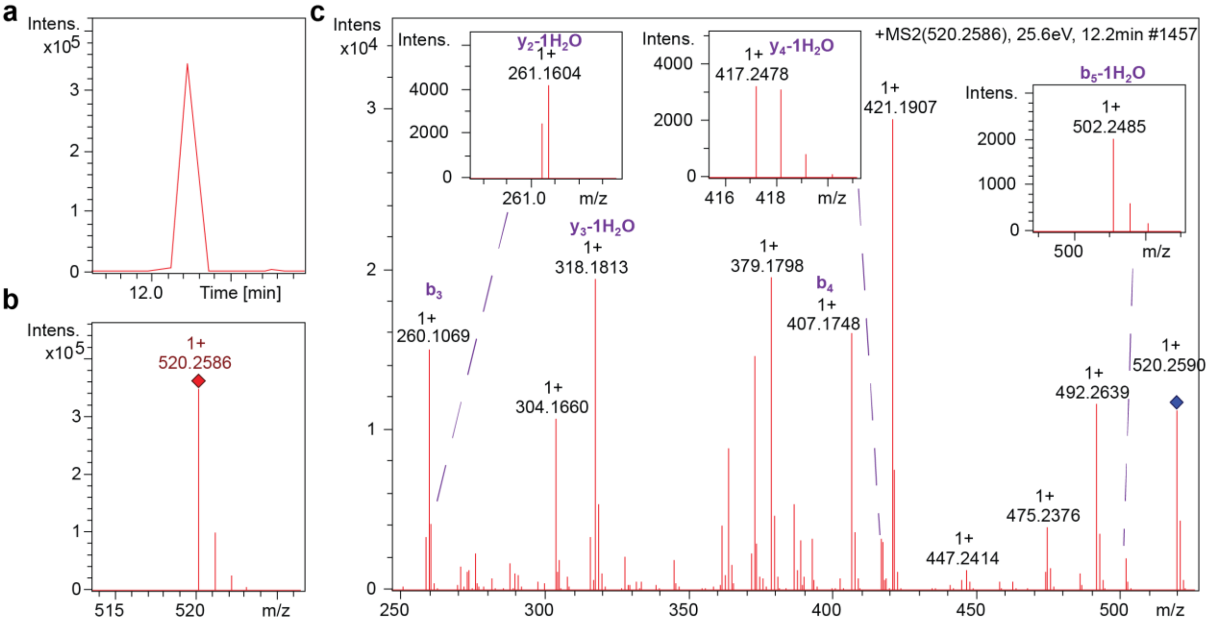

**d**

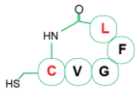

| ions | fragment | Observed m/z | Calculated m/z | charge | ppm |
|---|---|---|---|---|---|
| $y_2$-1$H_2O$ | FL | 261.1604 | 261.1598 | 1 | 2.47 |
| $y_3$-1$H_2O$ | GFL | 318.1813 | 318.1812 | 1 | 0.26 |
| $y_4$-1$H_2O$ | VGFL | 417.2478 | 417.2496 | 1 | -4.39 |
| $b_3$ | CVG | 260.1069 | 260.1063 | 1 | 2.16 |
| $b_4$ | CVGF | 407.1748 | 407.1748 | 1 | 0.12 |
| $b_5$-1H2O | CVGFL | 502.2485 | 502.2483 | 1 | 0.49 |

#### Supplementary Figure 27 UPLC-HRMS/MS analysis of synthesized BF_280_c

The retention time **a**, MS1 spectra **b,** and HRMS2-based fragmentations **c** of synthetic peptide. **d** Listed are the paired ions and possible fragments. The structure of synthetic AIP is displayed on the left side.

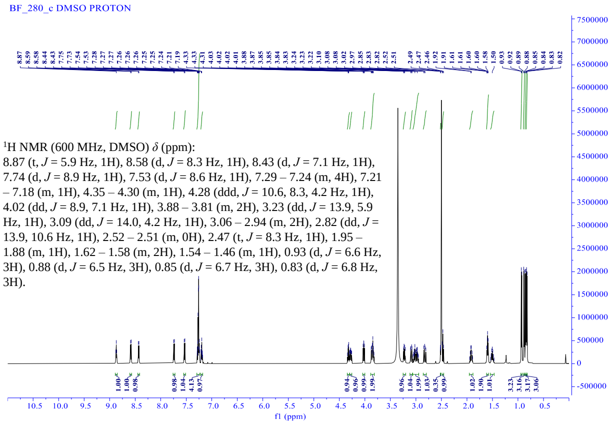

$^{1}$H NMR spectrum (600 MHz, DMSO-$d_6$) of BF_280_c.

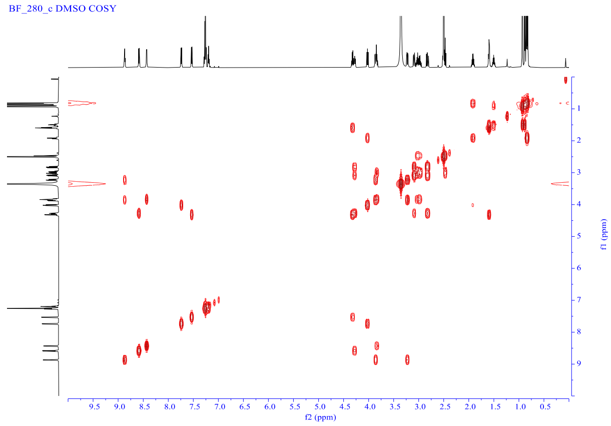

COSY NMR spectrum (600 MHz, DMSO-$d_6$) of BF_280_c.

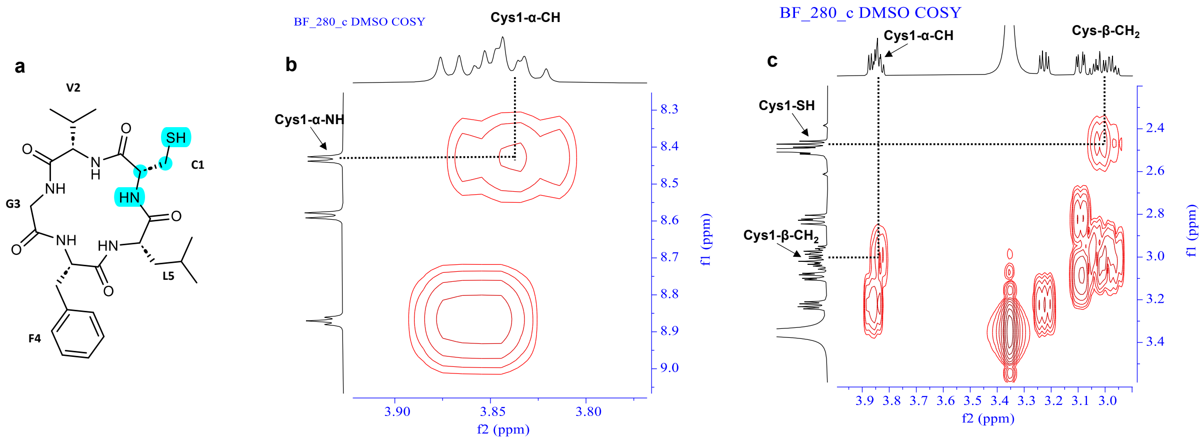

#### Supplementary Figure 28 NMR assignment of BF_280_c

Key COSY correlations for the proton in the cysteine residue of BF_280_c. **a** The chemical structure of BF_280_c, and the groups in the blue background corresponds to the proton in the key cysteine residue. **b** Key COSY correlations for Cys1-α-NH/Cys1-α-CH. **c** Key COSY correlations for Cys1-α-CH/Cys1-β-$CH_2$ and Cys1-SH/Cys1-β-$CH_2$.

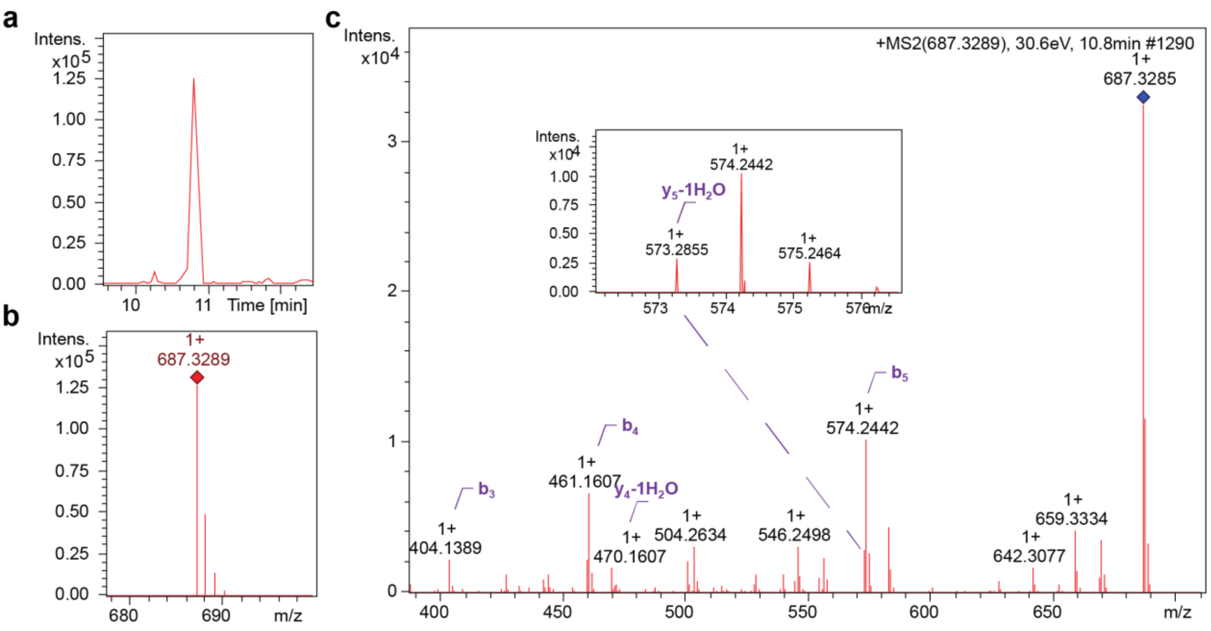

d

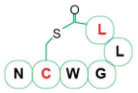

| ions | fragment | Observed m/z | Calculated m/z | charge | ppm |
|---|---|---|---|---|---|
| b3 | NCW | 404.1389 | 404.1387 | 1 | 0.45 |
| b4 | NCWG | 461.1607 | 461.1602 | 1 | 1.22 |
| b5 | NCWGL | 574.2442 | 574.2442 | 1 | 0.14 |
| y4-1H2O | WGLL | 470.2754 | 470.2762 | 1 | 1.69 |
| y5-1H2O | CWGLL | 573.2855 | 573.2854 | 1 | 0.15 |

#### Supplementary Figure 29 UPLC-HRMS/MS analysis of synthesized BF_398_et_L1

The retention time a, MS1 spectra b**,** and HRMS2-based fragmentations c of synthetic peptide. **d** Listed are the paired ions and possible fragments. The structure of synthetic AIP is displayed on the left side.

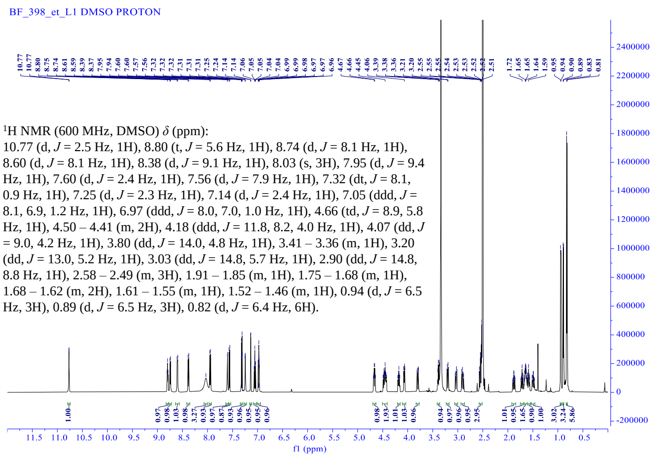

$^1$H NMR spectrum (600 MHz, DMSO-$d_6$) of BF_398_et_L1.

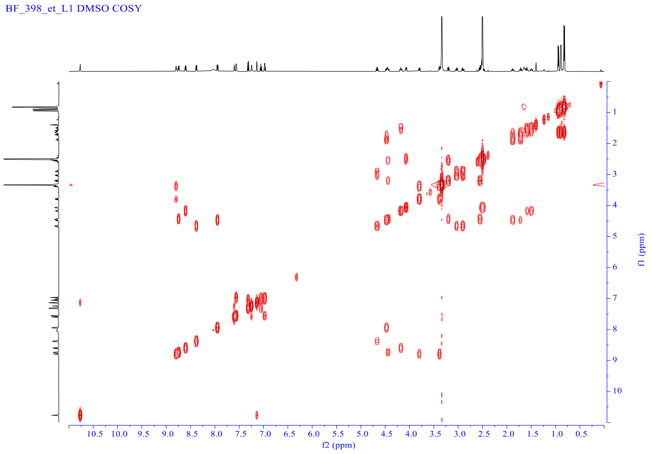

COSY NMR spectrum (600 MHz, DMSO-$d_6$) of BF_398_et_L1.

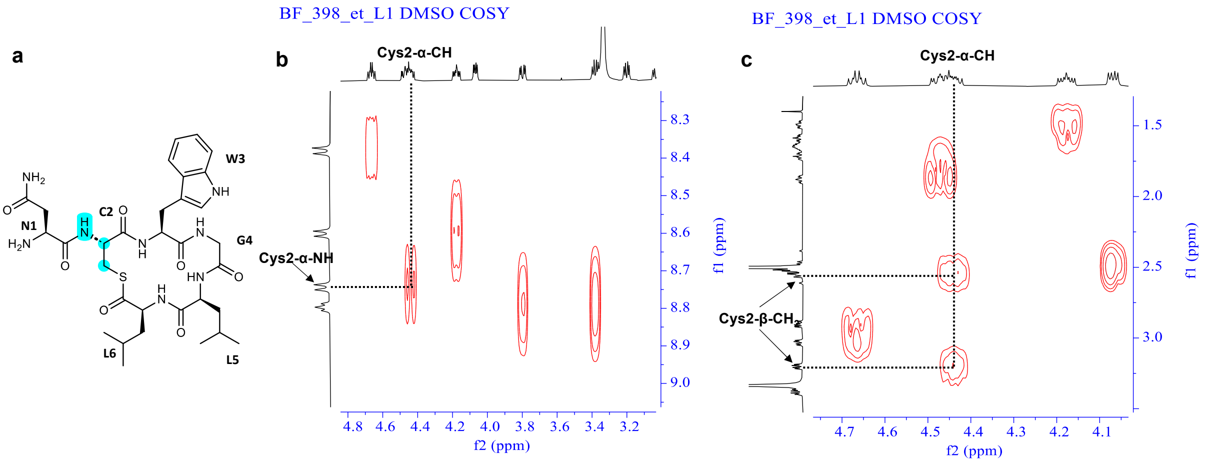

#### Supplementary Figure 30 NMR assignment of BF_398_et_L1

Key COSY correlations for the proton in the cysteine residue of BF_398_et_L1. **a** The chemical structure of BF_398_et_L1 and the groups in the blue background correspond to the proton in the key cysteine residue. **b** Key COSY correlations for Cys2-α-NH/Cys2-α-CH. **c** Key COSY correlations for Cys2-α-CH/Cys2-β-

CH2.

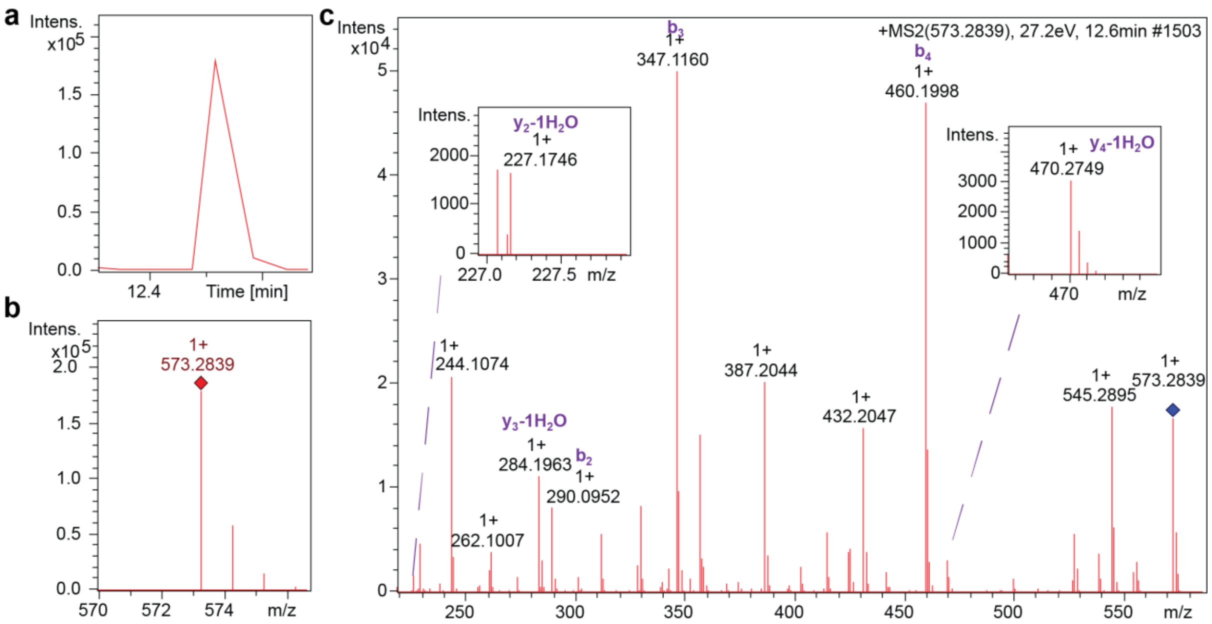

d

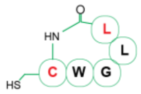

| ions | fragment | Observed m/z | Calculated m/z | charge | ppm |
|---|---|---|---|---|---|
| $y_2$-1$H_2O$ | LL | 227.1749 | 227.1754 | 1 | -2.22 |
| $y_3$-1$H_2O$ | GLL | 284.1963 | 284.1969 | 1 | -2.00 |
| $y_4$-1$H_2O$ | WGLL | 470.2749 | 470.2762 | 1 | -2.72 |
| $b_2$ | CW | 290.0952 | 290.0958 | 1 | -1.98 |
| $b_3$ | CWG | 347.1160 | 347.1172 | 1 | -3.57 |
| $b_4$ | CWGL | 460.1998 | 460.2013 | 1 | -3.26 |

**Supplementary Figure 31 UPLC-HRMS/MS analysis of synthesized BF_398_c**

The retention time **a**, MS1 spectra **b**, and HRMS2-based fragmentations c of synthetic peptide. **d** Listed are the paired ions and possible fragments. The structure of synthetic AIP is displayed on the left side.

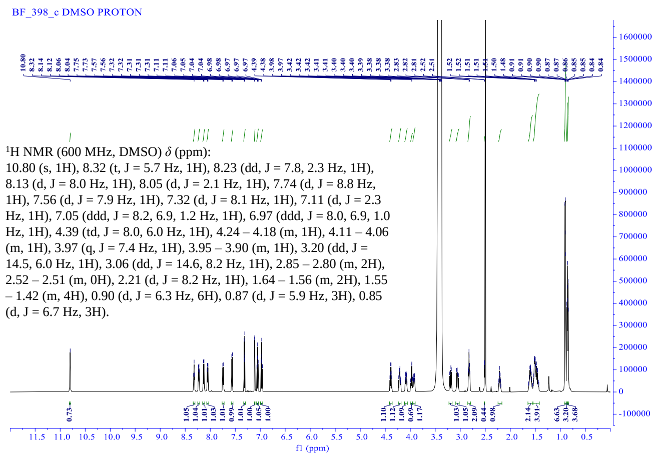

$^{1}H$ NMR spectrum (600 MHz, DMSO-$d_6$) of BF_398_c.

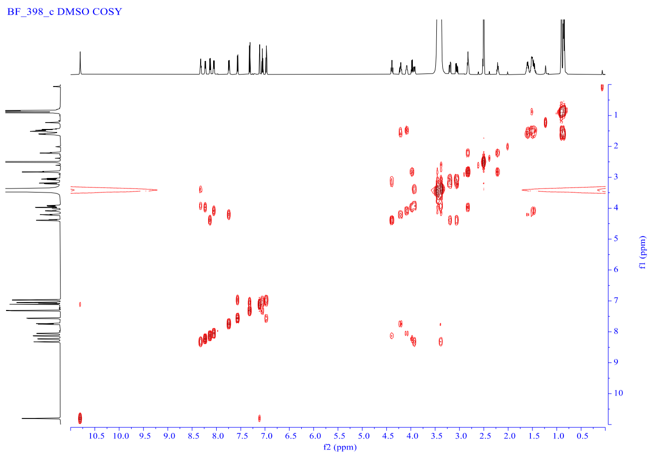

COSY NMR spectrum (600 MHz, DMSO-$d_6$) of BF_398_c.

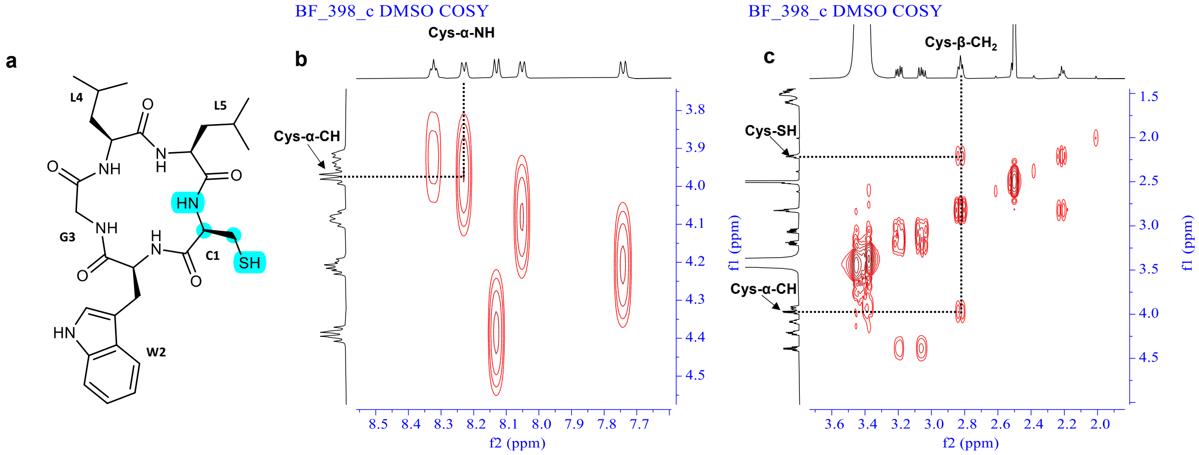

#### Supplementary Figure 32 NMR assignment of BF_398_c

Key COSY correlations for the proton in the serine residue of BF_398_c. **a** The chemical structure of BF_398_c, and the groups in the blue background corresponds to the proton in the key cysteine residue. **b** Key COSY correlations for Cys1-α-NH/Cys1-α-CH. c Key COSY correlations for Cys1-α-CH/Cys1-β-$CH_2$

and Cys1-SH/Cys1-β-$CH_2$.

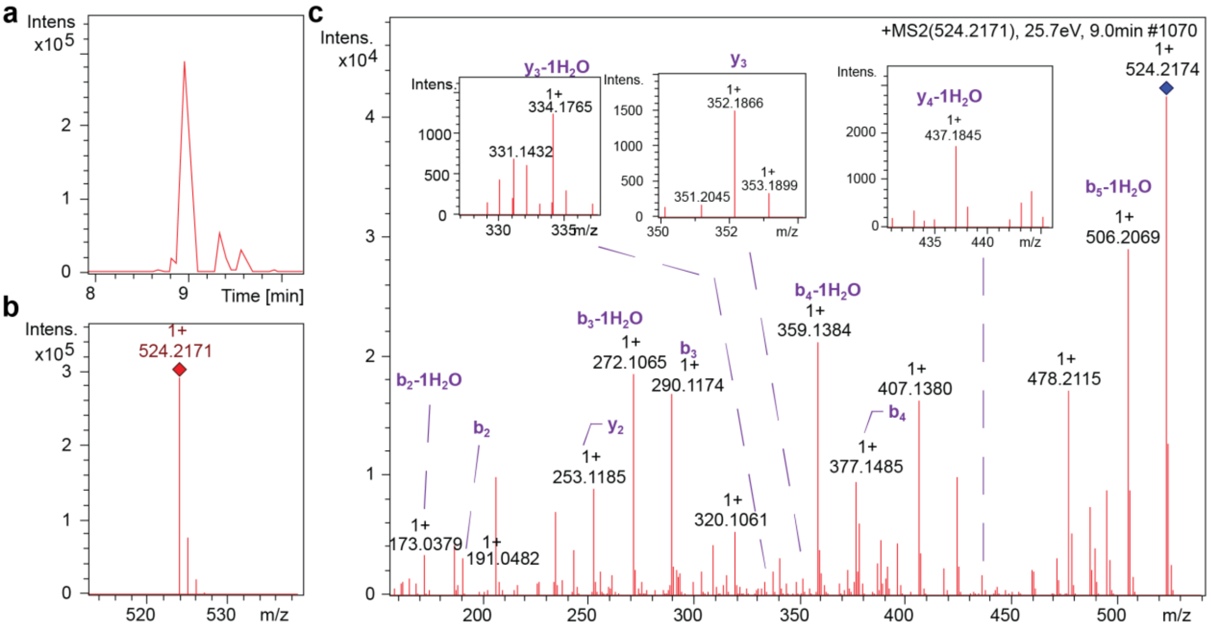

**d**

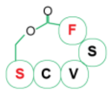

| ions | fragment | Observed m/z | Calculated m/z | charge | ppm |
|---|---|---|---|---|---|
| b2 | SC | 191.0482 | 191.0485 | 1 | 1.48 |
| b2-1$H_2O$ | SC | 173.0379 | 173.0379 | 1 | 0.26 |
| b3 | SCV | 290.1174 | 290.1169 | 1 | 1.71 |
| b3-1$H_2O$ | SCV | 272.1065 | 272.1063 | 1 | 0.69 |
| b4 | SCVS | 377.1485 | 377.1489 | 1 | 1.13 |
| b4-1$H_2O$ | SCVS | 359.1384 | 359.1384 | 1 | 0.03 |
| b5-1$H_2O$ | SCVSF | 506.2069 | 506.2068 | 1 | 0.24 |
| y2 | SF | 253.1185 | 253.1183 | 1 | 0.78 |
| y3 | VSF | 352.1866 | 352.1867 | 1 | 0.29 |
| y3-1$H_2O$ | VSF | 334.1765 | 334.1761 | 1 | 1.18 |
| y4-1$H_2O$ | CVSF | 437.1845 | 437.1853 | 1 | 1.92 |

#### Supplementary Figure 33 UPLC-HRMS/MS analysis of synthesized BF_488_et_free

The retention time **a**, MS1 spectra **b**, and HRMS2-based fragmentations **c** of synthetic peptide. **d** Listed are the paired ions and possible fragments. The structure of synthetic AIP is displayed on the left side.

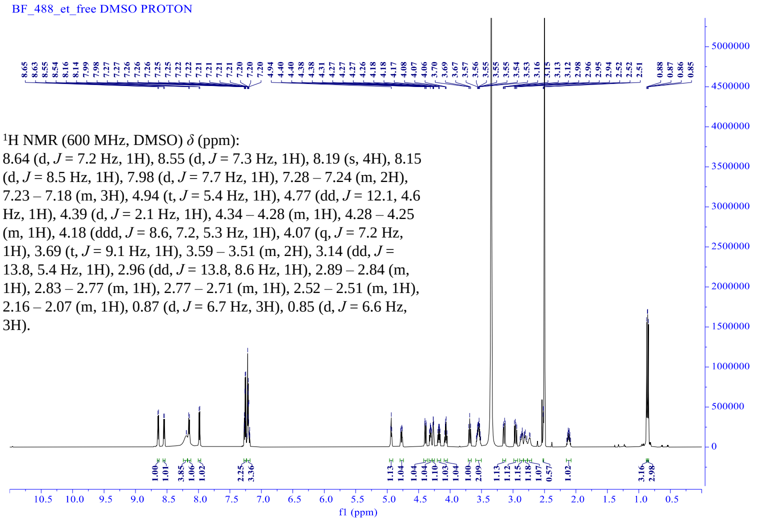

$^{1}$H NMR spectrum (600 MHz, DMSO-$d_6$) of BF_488_et_free.

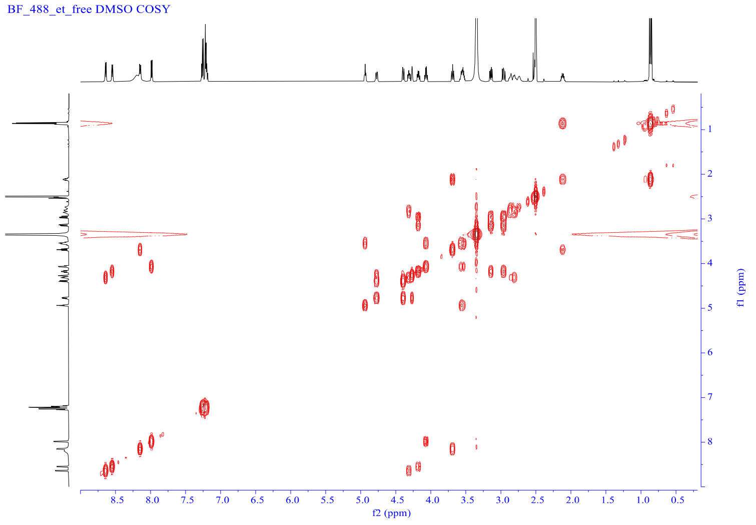

COSY NMR spectrum (600 MHz, DMSO-$d_6$) of BF_488_et_free.

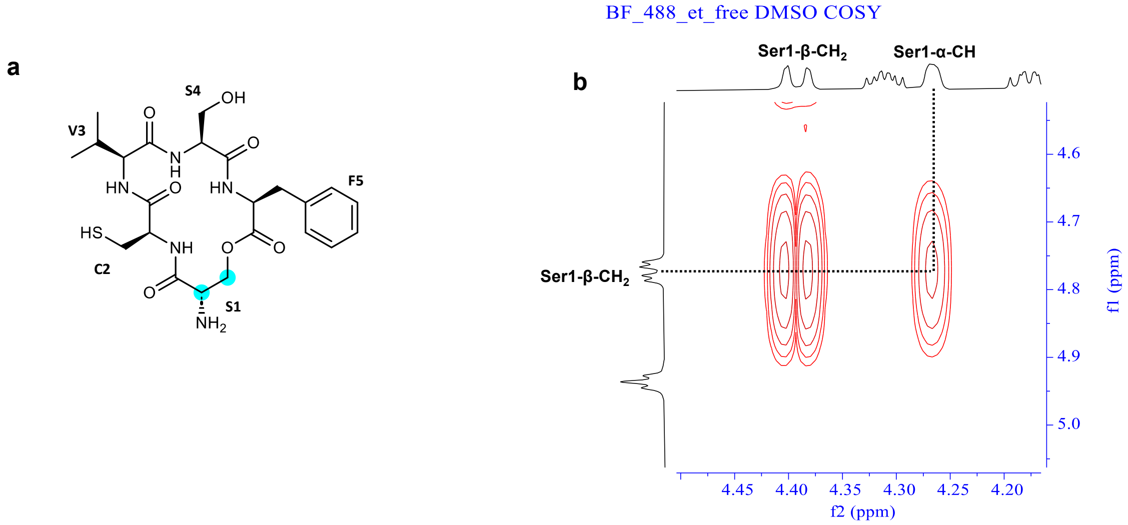

#### Supplementary Figure 34 NMR assignment of BF_488_et_free

Key COSY correlations for the proton in the serine residue of BF_488_et_free. **a** The chemical structure of BF_488_et_free and the groups in the blue background correspond to the proton in the key serine residue. **b** Key COSY correlations for Ser1-α-CH/Ser1-β-$CH_2$.

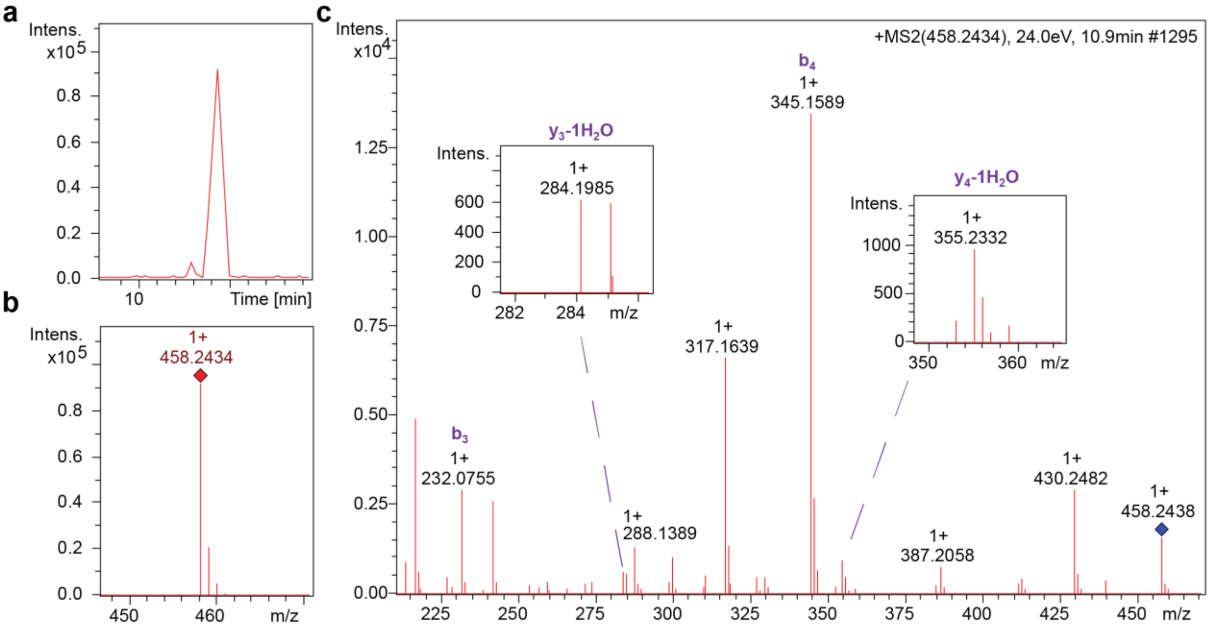

**d**

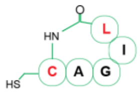

| ions | fragment | Observed m/z | Calculated m/z | charge | ppm |
|---|---|---|---|---|---|
| $y_3$-1$H_2O$ | GIL | 284.1985 | 284.1969 | 1 | 5.74 |
| $y_4$-1$H_2O$ | AGIL | 355.2332 | 355.2340 | 1 | -2.20 |
| $b_3$ | CAG | 232.0755 | 232.0750 | 1 | 1.99 |
| $b_4$ | CAGI | 345.1589 | 345.1591 | 1 | -0.59 |

#### Supplementary Figure 35 UPLC-HRMS/MS analysis of synthesized BF_598_c

The retention time **a**, MS1 spectra **b**, and HRMS2-based fragmentations **c** of synthetic peptide. **d** Listed are the paired ions and possible fragments. The structure of synthetic AIP is displayed on the left side.

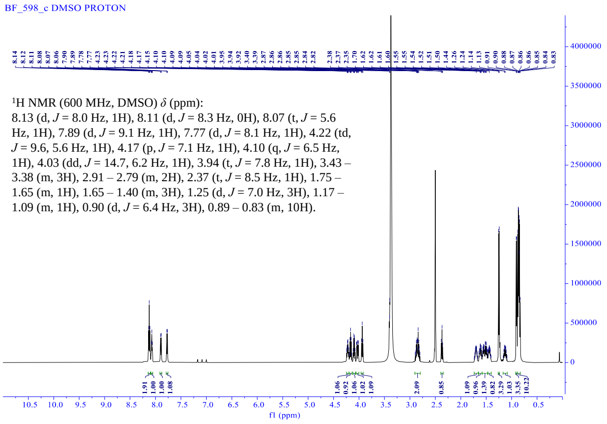

$^{1}$H NMR spectrum (600 MHz, DMSO-$d_6$) of BF_598_c.

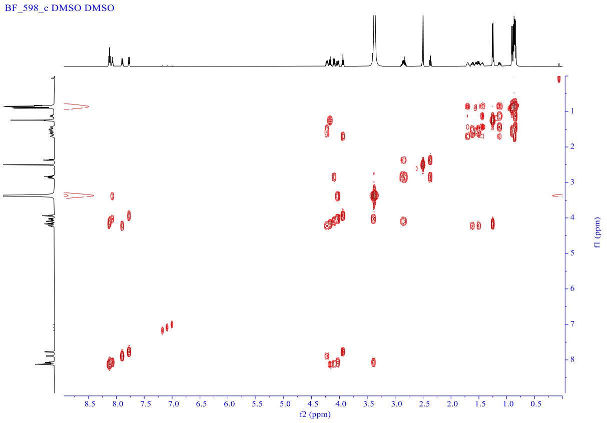

COSY NMR spectrum (600 MHz, DMSO-$d_6$) of BF_598_c.

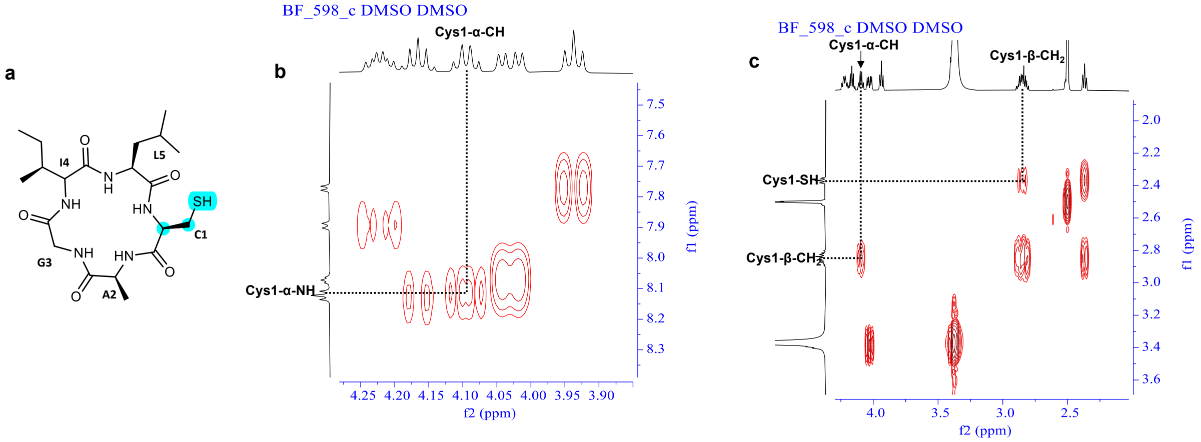

#### Supplementary Figure 36 NMR assignment of BF_598_c

Key COSY correlations for the proton in the cysteine residue of BF_598_c. **a** The chemical structure of BF_598_c and the groups in the blue background correspond to the proton in the key cysteine residue. **b** Key COSY correlations for Cys1-α-NH/Cys1-α-CH. **c** Key COSY correlations for Cys1-α-CH/Cys1-β-$CH_2$ and Cys1-SH/Cys1-β-$CH_2$.

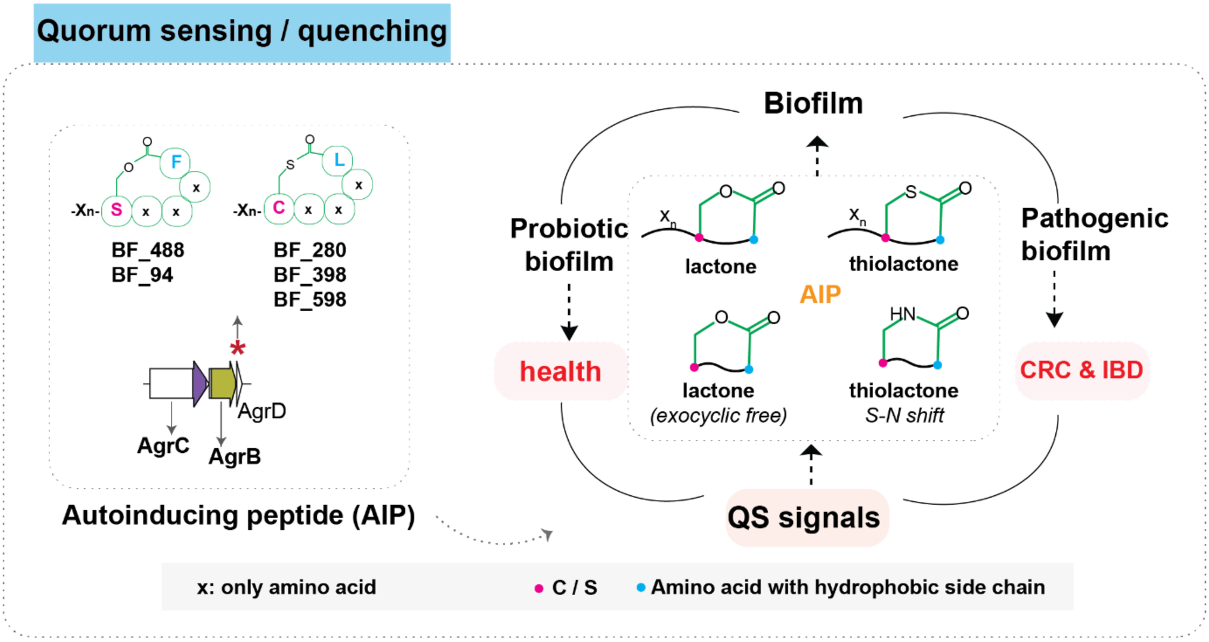

**Supplementary Figure 37 Potential quorum sensing or quorum quenching function of AIPs in mediating biofilm formation**

Given the significant impact of gut microbial biofilm on both disease pathogenesis[18], as in the case of pathogenic biofilm, and human health[19], as in the case of probiotic biofilm, we have directed our attention toward the study of AIPs that often involve in mediating biofilm formation. We thus aimed to validate six AIP families (BF_63, BF_94, BF_280, BF_398, BF_488, and BF_598) with an enrichment in the healthy microbiome compared to the diseased microbiome, particularly in CRC and IBD **(Fig. 4b, Supplementary Figure 18)**. **Left Panel**: The genomic context and potential structures of autoinducing peptide (AIP) families that are negatively associated with CRC and/or IBD are depicted. These families, exemplified by BF_488, harbored complete biosynthetic elements in their genomic context for biological activity[20]: membrane-bound endopeptidase AgrB, a sensor histidine kinase AgrC, and/or precursor AgrD domain. Moreover, the precursor sequence features indicated their potential to form a thiolactone (C-Xn-L) or lactone (S-Xn-L/F) functional group[21,22], which were required for potent biological activity. **Right Panel**: The functional groups of reported AIPs are shown in the inner circle. AIPs may be involved in biofilm development, which is related to human diseases. Xn represents the exocyclic tail of AIP with variable length.

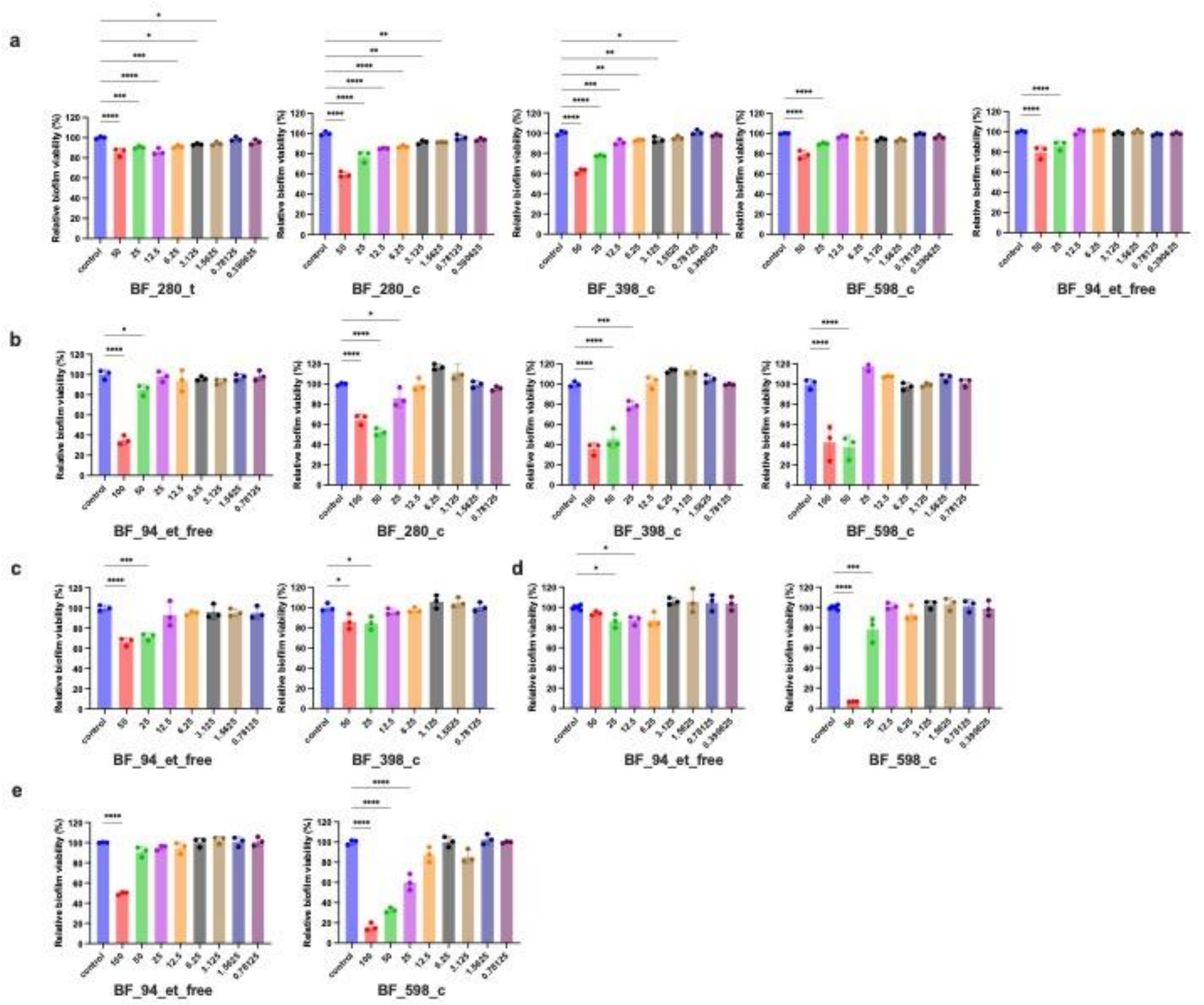

#### Supplementary Figure 38 AIPs inhibit clinical pathogenic biofilm formation

Crystal violet assay for antibiofilm activity against **a** *Staphylococcus aureus* ATCC43300, **b** *Listeria monocytogenes* ATCC19115, **c** *Peptostreptococcus stomatis* DSM17678, **d** *Streptococcus gallolyticus* subsp. Gallolyticus DSM 16831, **e** *Candida albicans* ATCC10231 biofilm formation during 24 h incubation. n = 3 biologically independent samples. Significance was determined using a student t-test. Bars represent mean ± standard error. For all *p* values: * 0.05 < $p$ < 0 .01, ** 0.001 < $p$ < 0 .01 and **** $p$ < 0.0001, mean significant difference compared with the control group.

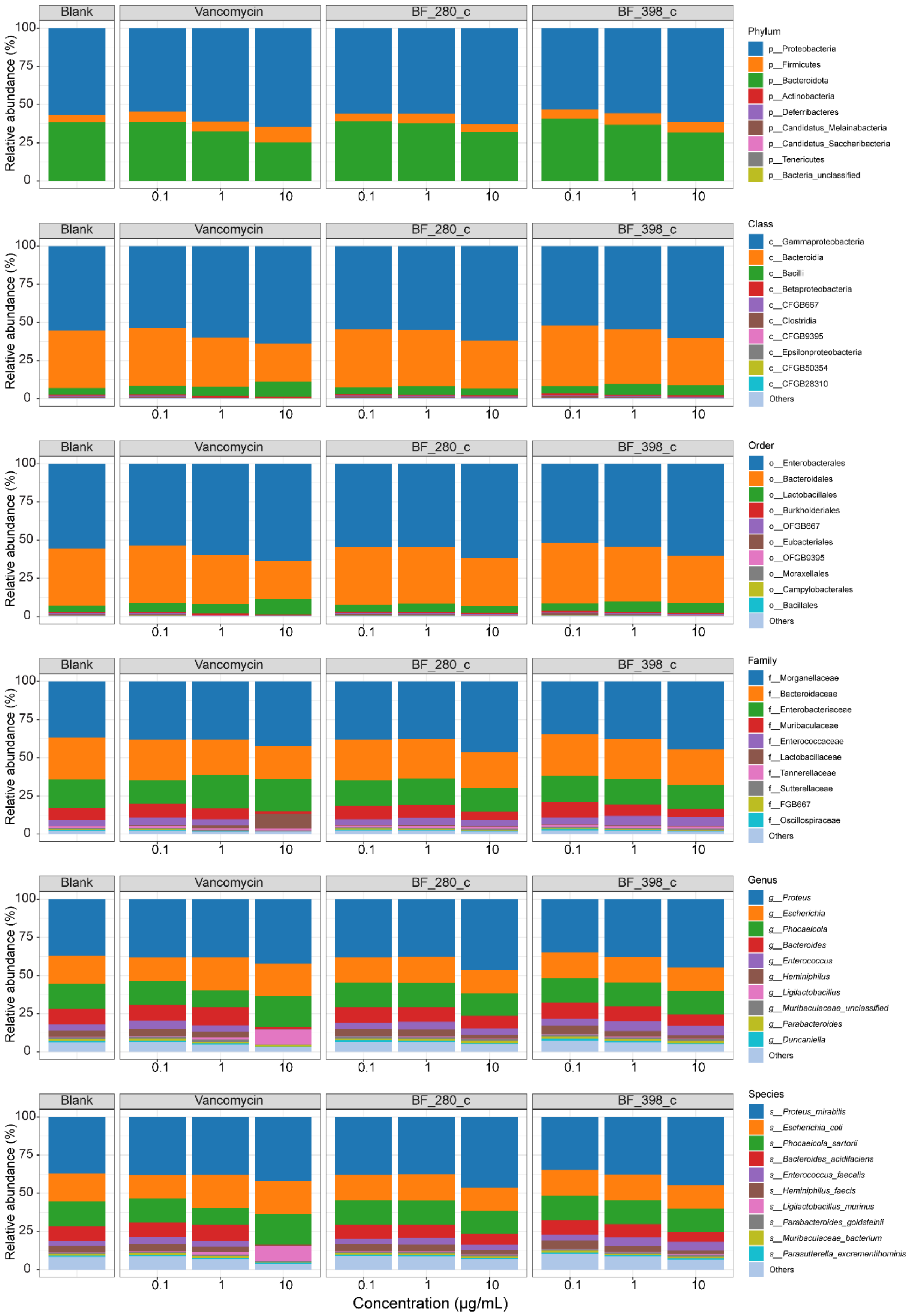
Blank
Vancomycin
BF_280_c
BF_398_c
Relative abundance (%)
100
75
50
25
0
0.1
1
10
Concentration (μg/mL)
Phylum
p__Proteobacteria
p__Firmicutes
p__Bacteroidota
p__Actinobacteria
p__Deferribacteres
p__Candidatus_Melainabacteria
p__Candidatus_Saccharibacteria
p__Tenericutes
p__Bacteria_unclassified
Class
c__Gammaproteobacteria
c__Bacteroidia
c__Bacilli
c__Betaproteobacteria
c__CFGB667
c__Clostridia
c__CFGB9395
c__Epsilonproteobacteria
c__CFGB50354
c__CFGB28310
Others
Order
o__Enterobacterales
o__Bacteroidales
o__Lactobacillales
o__Burkholderiales
o__OFGB667
o__Eubacteriales
o__OFGB9395
o__Moraxellales
o__Campylobacterales
o__Bacillales
Others
Family
f__Morganellaceae
f__Bacteroidaceae
f__Enterobacteriaceae
f__Muribaculaceae
f__Enterococcaceae
f__Lactobacillaceae
f__Tannerellaceae
f__Sutterellaceae
f__FGB667
f__Oscillospiraceae
Others
Genus
g__Proteus
g__Escherichia
g__Phocaeicola
g__Bacteroides
g__Enterococcus
g__Heminiphilus
g__Ligilactobacillus
g__Muribaculaceae_unclassified
g__Parabacteroides
g__Duncaniella
Others
Species
s__Proteus_mirabilis
s__Escherichia_coli
s__Phocaeicola_sartorii
s__Bacteroides_acidifaciens
s__Enterococcus_faecalis
s__Heminiphilus_faecis
s__Ligilactobacillus_murinus
s__Parabacteroides_goldsteinii
s__Muribaculaceae_bacterium
s__Parasutterella_excrementihominis
Others

#### Supplementary Figure 39 Relative abundance at the bacterial taxonomic levels of different treatments

Stacked bar plot showing the average relative abundance of bacterial phylum, class, order, family, genus and species (from top to bottom) in indicated samples. Only the top 10 abundant taxa are shown, and other taxa are grouped into “Others”.

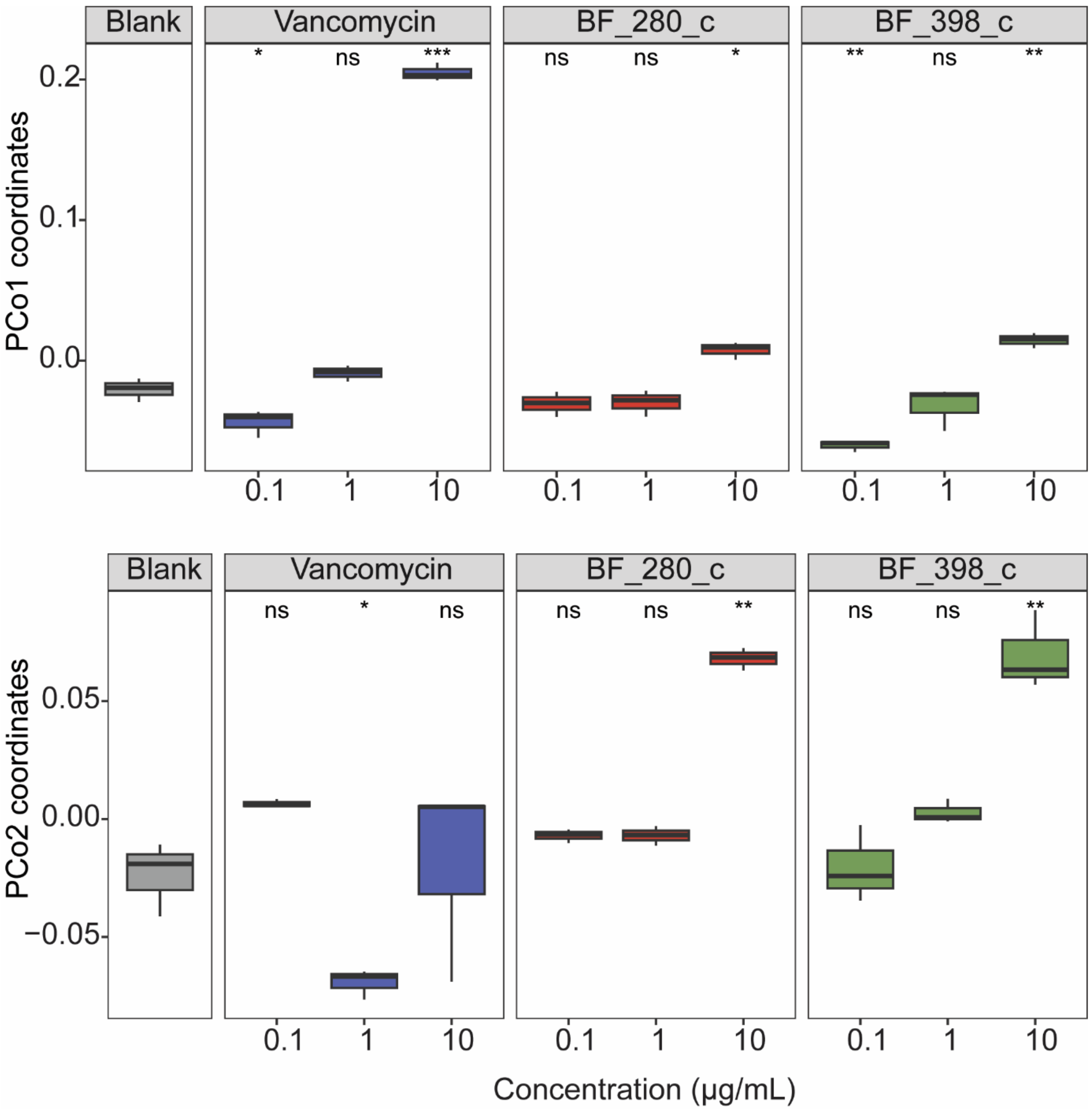

**Supplementary Figure 40 Coordinates of PCo1 and PCo2 axes of PCoA plot**

Boxplot showing the coordinates of two axes of the PCoA plot in Fig. 5d. Significances between treatment groups and blank group were indicated by using Welch's *t*-test. *$p < 0.05$; **$p < 0.01$; ***$p < 0.001$; ns, not significant.

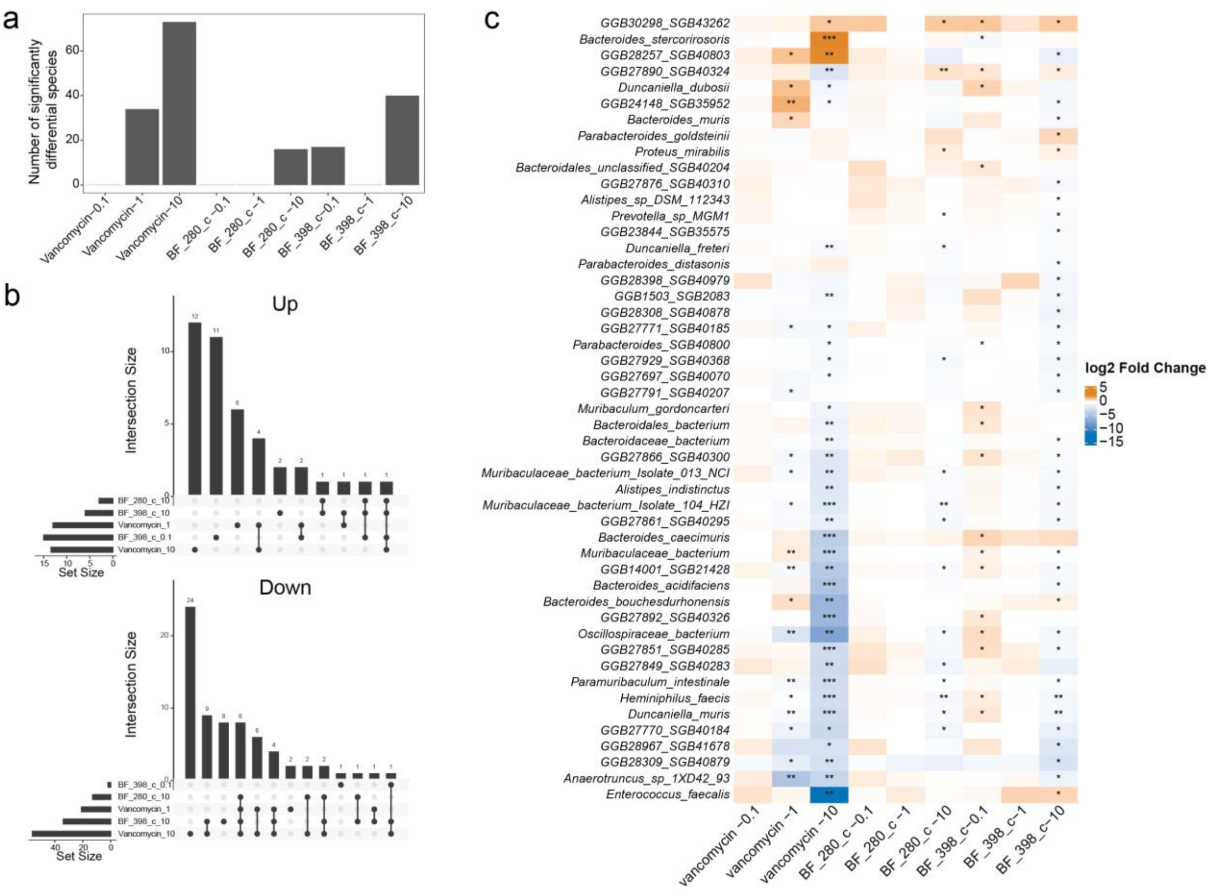

#### Supplementary Figure 41 Differentially abundant species between blank and treatment groups

**a** The number of significantly differentially abundant species between the blank and treatment groups, identified using MaAsLin2. **b** UpsetR visualization illustrating the intersection of differentially abundant species across various groups. The left bar plot indicates the count of species that showed increased (Up) or decreased (Down) abundance compared with blank group. The top bar plot represents the number of differentially abundant species in each intersection. Connecting lines are included for intersections present in multiple groups. c Heatmap displaying the log2 fold change in species abundance between the blank and treatment groups. Significance, as determined by MaAsLin2, is denoted by a plus sign. The differentially abundant species identified between the AIP and blank groups are highlighted. *$p$ < 0.05; **$p$ < 0.01; ***$p$ < 0.001; ns indicates non-significance.
